# Solute selectivity and functional divergence of aquaporins in reef-building corals

**DOI:** 10.64898/2026.10.13.758710

**Authors:** Ayumi Nagashima, Akira Kato

## Abstract

Reef-building corals are increasingly threatened by environmental stressors, including ocean warming, acidification, and osmotic stress caused by fluctuations in seawater salinity, highlighting the need for a better understanding of coral physiology. However, the molecular mechanisms by which corals regulate water and solute transport remain poorly understood. To investigate the role of aquaporins (Aqps) in coral physiology, we examined the evolutionary history and transport properties of the Aqp family in reef-building corals. Comparative genomic analyses across cnidarians revealed that the nine-member core hexacorallian *aqp* repertoire, comprising *aqp4L1.1*, *aqp4L1.2*, *aqp4L1.3*, *aqp4L2*, *aqp4L3*, *glp1*, *glp2.1*, *glp2.2*, and *aqp12L1*, has been highly conserved since the common ancestor of reef-building corals and sea anemones, whereas *aqp8L* was consistently absent across all examined hexacorallian species. Conservation of synteny and exon–intron structures further confirmed that major components of the *aqp* family architecture originated deep in metazoan evolution. Functional characterization of *Acropora millepora* Aqp4L and Glp proteins demonstrated that most transport properties were consistent with phylogenetic predictions, with the majority of Aqp4L proteins exhibiting water-selective permeability, whereas Glp proteins transported both water and small solutes such as glycerol. Unexpectedly, Aqp4L1.2, which belongs to the water-specific aquaporin lineage, transported not only water but also glycerol, urea, and boric acid, suggesting functional co-option after gene duplication and underscoring the evolutionary flexibility of the Aqp4L lineage. These findings suggest that the hexacorallian *aqp* family evolved through a combination of deep evolutionary conservation and lineage-specific functional innovation, providing new insight into the molecular foundations of coral physiology.

**Significance Statement:** How reef-building corals regulate water and solute transport at the molecular level remains largely unknown. Here, comparative genomic and functional analyses revealed the evolutionary history and transport properties of the *aqp* family in reef-building corals. Reef-building corals retain a conserved *aqp* repertoire shared with sea anemones, comprising the *aqp4-like*, *aquaglyceroporin*, and *aqp12-like* subfamilies, whereas the *aqp8-like* subfamily is entirely absent. Phylogenetic analyses further showed that these genes form hexacorallian-specific clades distinct from homologs in other cnidarian lineages, indicating that the fundamental architecture of the hexacorallian *aqp* repertoire was established in their common ancestor. Unexpectedly, we identified a duplicated water-channel gene that evolved the capacity to transport multiple solutes, revealing a previously unrecognized example of functional co-option in coral aquaporins. These findings show that coral aquaporin evolution has been driven by both long-term conservation and lineage-specific innovation, providing a genomic and functional framework for understanding the molecular basis of coral physiology.

## Introduction

Coral reefs are among the most biodiverse marine ecosystems and are increasingly threatened by environmental change. Climate change-driven ocean warming and acidification are major causes of coral bleaching, impaired calcification, and reef degradation (Hughes et al. 2017; Pandolfi et al. 2003; Tambutte et al. 2015). In addition to thermal and acidic stress, fluctuations in seawater salinity represent another critical, yet poorly understood, driver of coral degradation. Although reef-building corals are generally regarded as stenohaline osmoconformers (Aguilar et al. 2019; Ferrier-Pagès et al. 1999; Mayfield and Gates 2007), many reef ecosystems are located in shallow coastal environments where corals can experience substantial fluctuations in seawater salinity (Kerswell and Jones 2003; Lan et al. 2025). Both hypo- and hypersaline conditions have been associated with coral bleaching, mortality, and alterations in physiological performance (Bahr et al. 2015; Coles 2003; Jokiel et al. 1993; Porter et al. 1999; Rodgers et al. 2021). Recent studies have shown that the model sea anemone *Aiptasia* responds to hypersaline conditions, thereby enhancing its thermotolerance and mitigating heat-induced bleaching (Gegner et al. 2019). In addition, osmotic stress influences the stress tolerance of both cnidarian hosts and their Symbiodiniaceae symbionts (Ochsenkuhn et al. 2017). However, the precise cellular and molecular mechanisms by which the coral host regulates intracellular osmolarity and facilitates the transport of low-molecular-weight compounds remain to be elucidated.

In diverse organisms ranging from microbes and plants to metazoans, the aquaporin (Aqp) water channel superfamily plays a crucial role in maintaining cellular volume and osmotic homeostasis (Finn et al. 2014). Aqps are typically 250–300 amino acid membrane proteins that assemble into functional tetramers and are classified according to their solute selectivity into classical water-specific Aqps, aquaglyceroporins, ammoniaporins, and unorthodox Aqps (Takata et al. 2004). Genomic and transcriptomic surveys have identified representatives of each of these major subfamilies in reef-building corals, including water-specific Aqps (Aqp4-like), aquaglyceroporins (Glp), ammoniaporins (Aqp8-like), and unorthodox Aqps (Aqp12-like). Whereas Glp proteins are predicted to transport water, glycerol, and urea, and Aqp8-like proteins are predicted to transport water, ammonia, and urea, the transport properties of many coral Aqps and Glps remain unknown (Finn et al. 2014). While vertebrate Aqps have been extensively studied with respect to gene duplication, subfunctionalization, neofunctionalization, and diversification of solute permeability (Imaizumi et al. 2024; Nagashima et al. 2025), the precise functional and evolutionary configurations of these solute-transporting channels in basal metazoans like stony corals remain largely unexplored.

Over the past two decades, high-throughput sequencing technologies have generated high-quality genome assemblies for diverse anthozoans, including sea anemones, reef-building corals, and soft corals. In 2007, the initial sequencing of the sea anemone *Nematostella vectensis* genome established a foundation for basal metazoan genomics (Putnam et al. 2007), which has recently been updated to high-quality chromosome-level assemblies such as jaNemVect1.1 by the Wellcome Sanger Institute. Since then, Shinzato et al. (2011) reported the genome of a reef-building coral, *Acropora digitifera* (Adig_1.1), which provided critical insights for understanding coral symbiosis and environmental stress responses (Shinzato et al. 2011). Genomic resources for reef-building corals have expanded rapidly, spanning major families and genera; these include high-quality assemblies of *Acropora millepora* (Amil_v2.1; (Fuller et al. 2020), *Porites lutea* (jaPorLute2.1; Wellcome Sanger Institute), *Stylophora pistillata* (Voolstra et al. 2017), *Pocillopora damicornis* (Flot and Tillier 2007), and *Orbicella faveolata* (Smithsonian Tropical Research Institute). In addition, the genomes of sea anemones such as *Exaiptasia diaphana* (Baumgarten et al. 2015) and *Actinia tenebrosa* (Surm et al. 2019), as well as the soft corals *Dendronephthya gigantea* (Jeon et al. 2019) and *Xenia* sp. (Hu et al. 2020), have been successfully decoded. Although these genomic resources have revealed numerous putative *aqp*s across anthozoans, many annotations remain incomplete and the transport properties of most predicted proteins have not been experimentally validated.

Despite growing evidence that salinity stress influences coral performance and survival, the molecular mechanisms governing water and solute transport in corals remain poorly understood. Aquaporins are prime candidates for mediating these processes, yet their evolutionary history and physiological functions in reef-building corals have not been comprehensively investigated. The objective of this study was to elucidate the evolutionary diversification and physiological functions of the Aqp superfamily in reef-building corals through comprehensive identification of Aqp repertoires and functional characterization of their encoded proteins. First, a comparative genomic analysis was conducted to examine whether the genetic architecture of Aqp is conserved between reef-building corals and other cnidarian lineages. Next, the Aqp4L and Glp genes from the reef-building coral *Acropora* were expressed in *Xenopus* oocytes, and their transport activities were subsequently measured. This comprehensive functional characterization provides crucial molecular insights that enhance our understanding of how reef-building corals manage osmotic stress in a rapidly changing marine environment. Throughout this study, gene names are shown in lowercase italics, whereas the corresponding proteins are indicated by non-italicized names beginning with a capital letter.

## Results

### *aqp* genes in cnidarians and basal metazoans

To elucidate the genomic composition and evolutionary history of the *aqp* gene family in Hexacorallia, we comprehensively identified *aqp*s from representative hexacorallian genomes together with those of other cnidarians, including hydrozoans, and basal metazoan outgroups, including ctenophores and poriferans (**Table 1**). Predicted amino acid sequences and corresponding accession numbers are provided in **Table 2**, and **Tables S1**–**S3**, and are indicated in the phylogenetic tree (**Figure 1**). Phylogenetic analyses revealed a highly conserved core *aqp* repertoire across Hexacorallia. Reef-building corals and sea anemones consistently possessed orthologs of *aqp4L*, *glp*, and *aqp12L*, and all corresponding protein sequences grouped into well-supported hexacorallian clades (**Figure 1**). These results indicate strong conservation of the core *aqp* family architecture within the subclass. In contrast, *aqp* sequences from octocorals, scyphozoans, and hydrozoans formed distinct lineage-specific clades, whereas reef-building corals and sea anemones retained clear one-to-one orthologous relationships within each of the *aqp4L*, *glp*, and *aqp12L* lineages (**Figure 1**).

**Fig. 1.**
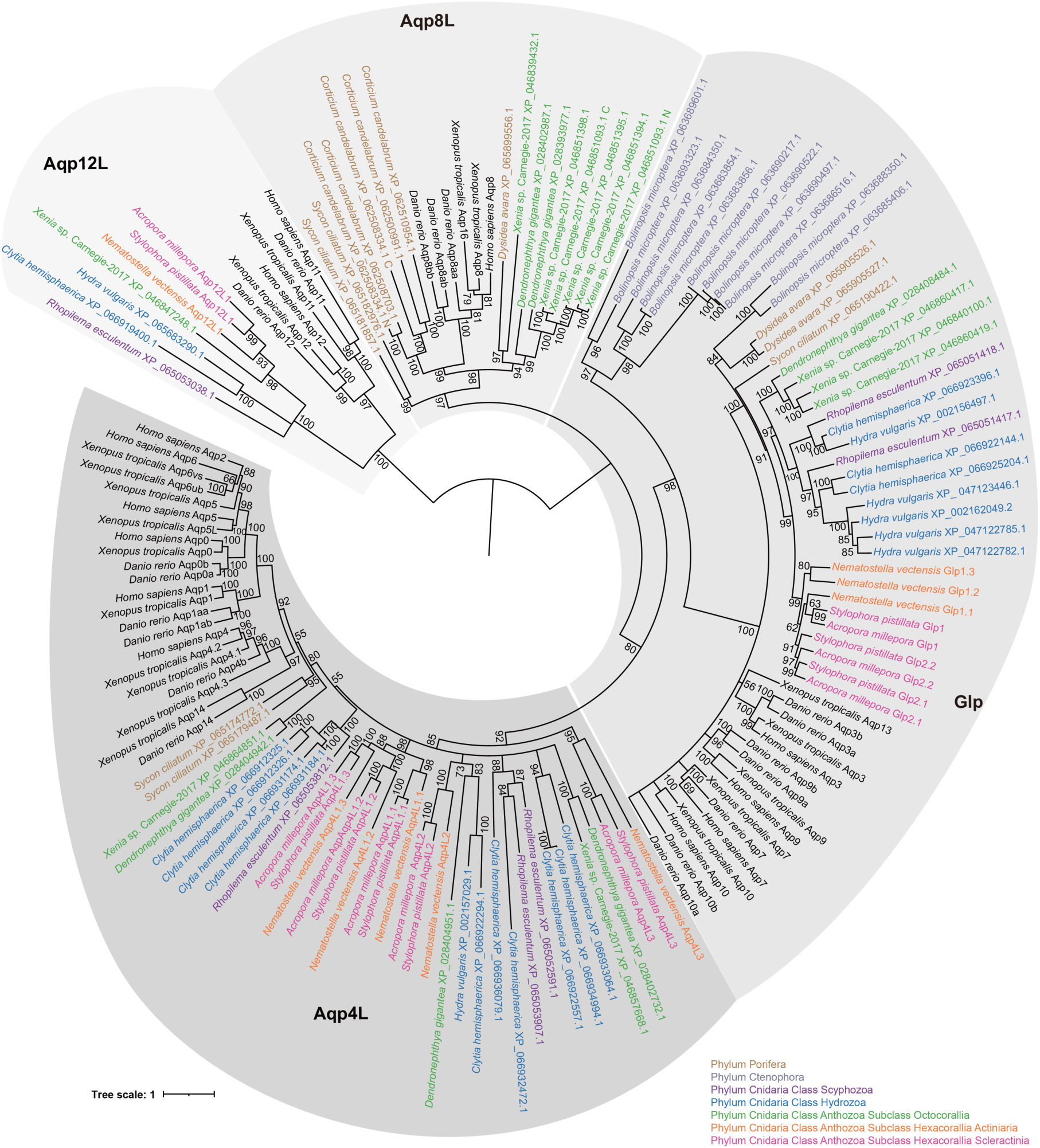
Phylogenetic relationship of Aqp sequences from Porifera, Ctenophora, Cnidaria, and Chordata. Phylogenetic tree and classification of Aqp members. Numbers indicate bootstrap values. Predicted amino acid sequences and corresponding accession numbers are provided in **Table 2**, and **Tables S1-S3**.

**Table 1.**
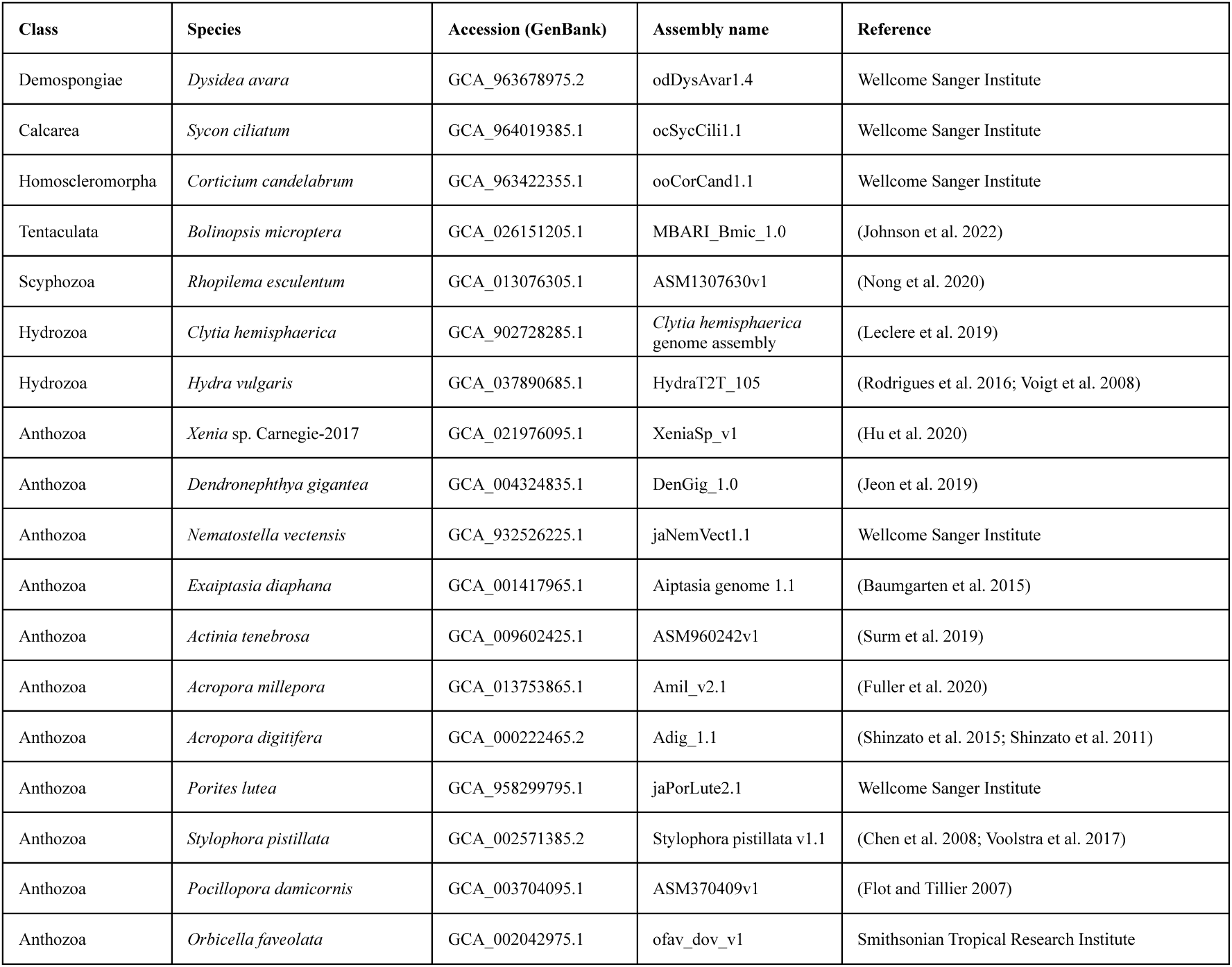
Genome databases analyzed in this study.

**Table 2.**
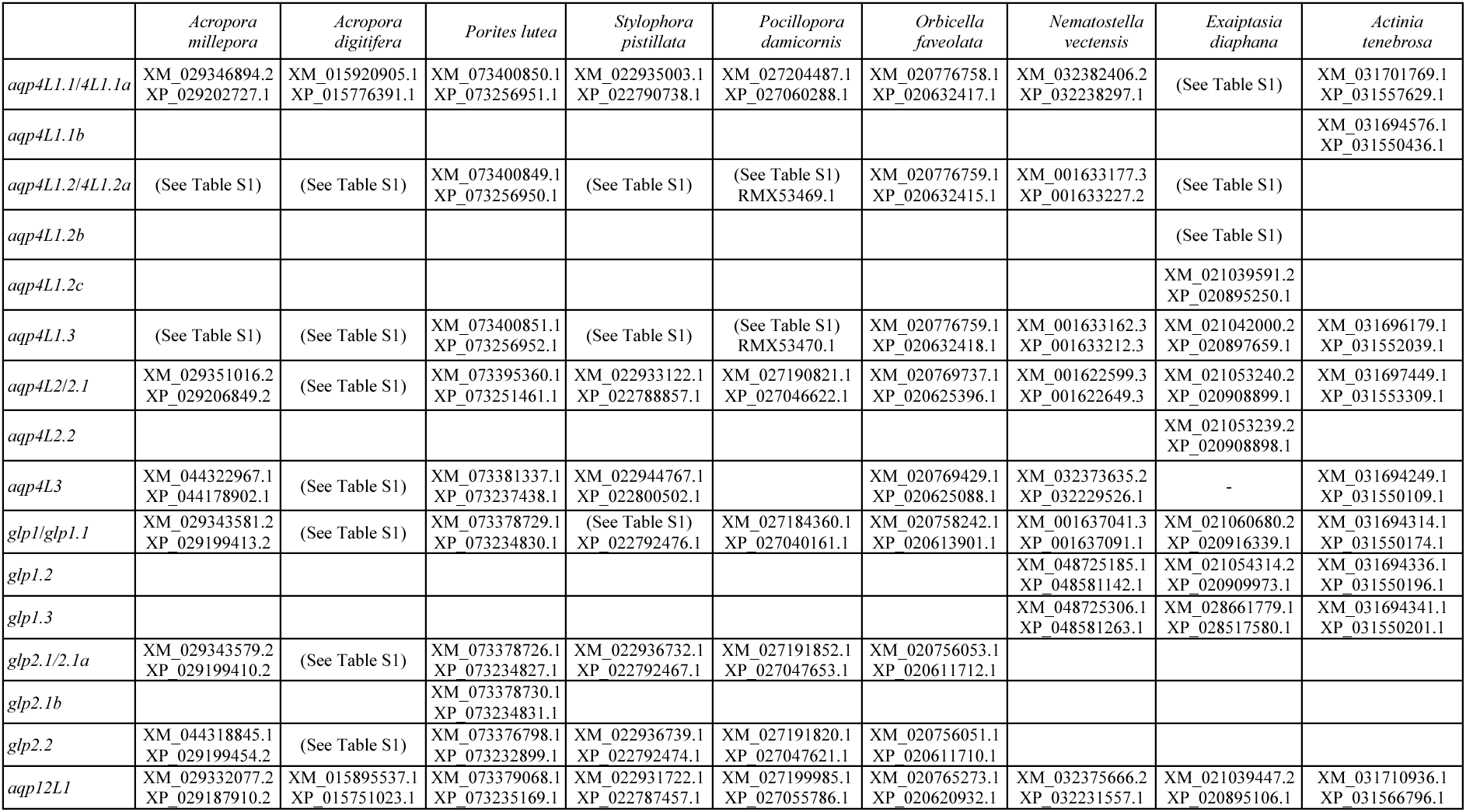
Accession numbers for the *aqp* genes and their encoded proteins in Scleractinia and Actiniaria.

|  | *Acropora millepora* | *Acropora digitifera* | *Porites lutea* | *Stylophora pistillata* | *Pocillopora damicornis* | *Orbicella faveolata* | *Nematostella vectensis* | *Exaiptasia diaphana* | *Actinia tenebrosa* |
| --- | --- | --- | --- | --- | --- | --- | --- | --- | --- |
| *aqp4L1.1/4L1.1a* | XM_029346894.2<br>XP_029202727.1 | XM_015920905.1<br>XP_015776391.1 | XM_073400850.1<br>XP_073256951.1 | XM_022935003.1<br>XP_022790738.1 | XM_027204487.1<br>XP_027060288.1 | XM_020776758.1<br>XP_020632417.1 | XM_032382406.2<br>XP_032238297.1 | (See Table S1) | XM_031701769.1<br>XP_031557629.1 |
| *aqp4L1.1b* |  |  |  |  |  |  |  |  | XM_031694576.1<br>XP_031550436.1 |
| *aqp4L1.2/4L1.2a* | (See Table S1) | (See Table S1) | XM_073400849.1<br>XP_073256950.1 | (See Table S1) | (See Table S1)<br>RMX53469.1 | XM_020776759.1<br>XP_020632415.1 | XM_001633177.3<br>XP_001633227.2 | (See Table S1) |  |
| *aqp4L1.2b* |  |  |  |  |  |  |  | (See Table S1) |  |
| *aqp4L1.2c* |  |  |  |  |  |  |  | XM_021039591.2<br>XP_020895250.1 |  |
| *aqp4L1.3* | (See Table S1) | (See Table S1) | XM_073400851.1<br>XP_073256952.1 | (See Table S1) | (See Table S1)<br>RMX53470.1 | XM_020776759.1<br>XP_020632418.1 | XM_001633162.3<br>XP_001633212.3 | XM_021042000.2<br>XP_020897659.1 | XM_031696179.1<br>XP_031552039.1 |
| *aqp4L2/2.1* | XM_029351016.2<br>XP_029206849.2 | (See Table S1) | XM_073395360.1<br>XP_073251461.1 | XM_022933122.1<br>XP_022788857.1 | XM_027190821.1<br>XP_027046622.1 | XM_020769737.1<br>XP_020625396.1 | XM_001622599.3<br>XP_001622649.3 | XM_021053240.2<br>XP_020908899.1 | XM_031697449.1<br>XP_031553309.1 |
| *aqp4L2.2* |  |  |  |  |  |  |  | XM_021053239.2<br>XP_020908898.1 |  |
| *aqp4L3* | XM_044322967.1<br>XP_044178902.1 | (See Table S1) | XM_073381337.1<br>XP_073237438.1 | XM_022944767.1<br>XP_022800502.1 |  | XM_020769429.1<br>XP_020625088.1 | XM_032373635.2<br>XP_032229526.1 | - | XM_031694249.1<br>XP_031550109.1 |
| *glp1/1.1* | XM_029343581.2<br>XP_029199413.2 | (See Table S1) | XM_073378729.1<br>XP_073234830.1 | (See Table S1)<br>XP_022792476.1 | XM_027184360.1<br>XP_027040161.1 | XM_020758242.1<br>XP_020613901.1 | XM_001637041.3<br>XP_001637091.1 | XM_021060680.2<br>XP_020916339.1 | XM_031694314.1<br>XP_031550174.1 |
| *glp1.2* |  |  |  |  |  |  | XM_048725185.1<br>XP_048581142.1 | XM_021054314.2<br>XP_020909973.1 | XM_031694336.1<br>XP_031550196.1 |
| *glp1.3* |  |  |  |  |  |  | XM_048725306.1<br>XP_048581263.1 | XM_028661779.1<br>XP_028517580.1 | XM_031694341.1<br>XP_031550201.1 |
| *glp2.1/2.1a* | XM_029343579.2<br>XP_029199410.2 | (See Table S1) | XM_073378726.1<br>XP_073234827.1 | XM_022936732.1<br>XP_022792467.1 | XM_027191852.1<br>XP_027047653.1 | XM_020756053.1<br>XP_020611712.1 |  |  |  |
| *glp2.1b* |  |  | XM_073378730.1<br>XP_073234831.1 |  |  |  |  |  |  |
| *glp2.2* | XM_044318845.1<br>XP_029199454.2 | (See Table S1) | XM_073376798.1<br>XP_073232899.1 | XM_022936739.1<br>XP_022792474.1 | XM_027191820.1<br>XP_027047621.1 | XM_020756051.1<br>XP_020611710.1 |  |  |  |
| *aqp12L1* | XM_029332077.2<br>XP_029187910.2 | XM_015895537.1<br>XP_015751023.1 | XM_073379068.1<br>XP_073235169.1 | XM_022931722.1<br>XP_022787457.1 | XM_027199985.1<br>XP_027055786.1 | XM_020765273.1<br>XP_020620932.1 | XM_032375666.2<br>XP_032231557.1 | XM_021039447.2<br>XP_020895106.1 | XM_031710936.1<br>XP_031566796.1 |

Comparison of *aqp* gene repertoires among representative metazoan lineages further supported this pattern (**Figure 2**). Consistent with the evolutionary framework proposed by Finn et al. (2014) (Finn et al. 2014), *aqp8* orthologs were not detected in any examined hexacorallian species. In contrast, *aqp8* was retained in octocorals, consistent with a loss prior to the diversification of extant Hexacorallia. Together, these findings are consistent with the hypothesis that the ancestral hexacorallian *aqp* repertoire primarily comprised *aqp4L*, *glp*, and *aqp12L* genes.

**Fig. 2.**
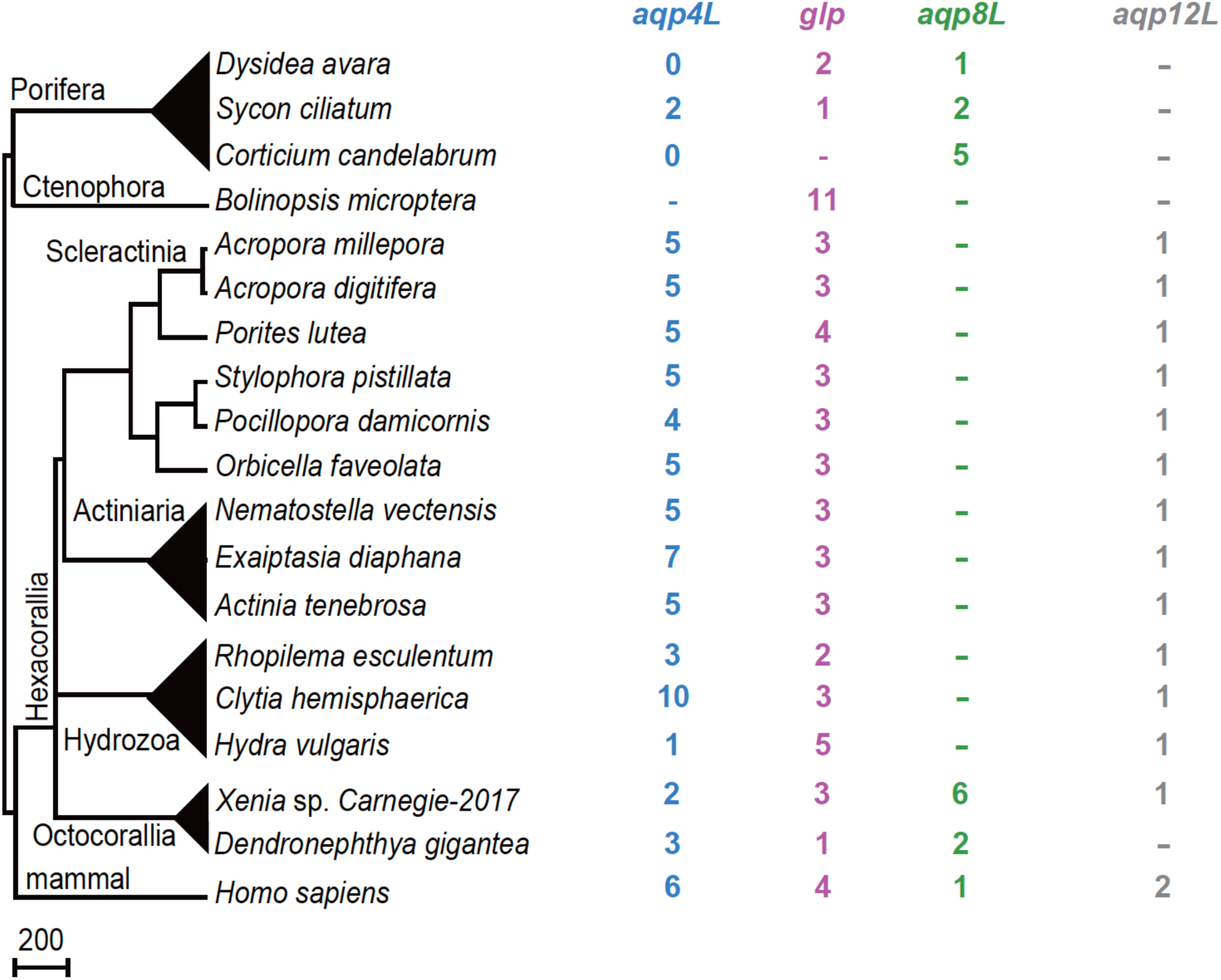
Comparison of *aqp* repertoires among representative metazoan species. The species phylogeny is shown on the left. Numbers indicate the copy number of *aqp4L*, *glp*, *aqp8L*, and *aqp12L* subfamilies based on the phylogenetic analyses shown in Fig. 1. Dashes indicate the absence of an identified ortholog.

To further resolve relationships among hexacorallian *aqp*s, we reconstructed a phylogeny using a hexacorallian-specific dataset (**Figure 3a**). All sequences used in this analysis and their corresponding accession numbers are listed in **Tables 2** and **Table S1**. Our expanded taxonomic sampling also provided an opportunity to refine several *aqp* annotations proposed by Finn et al. (2014) (Finn et al. 2014). Based on the phylogenetic relationships recovered here, the genes previously designated *aqp4L1*, *aqp4L2*, *aqp4L3*, and *aqp4L4* were reassigned as *aqp4L1.1*, *aqp4L1.2*, *aqp4L1.3*, and *aqp4L2*, respectively. Notably, *aqp4L1.3* was not reported in the previous study and was newly identified in our analysis. In addition, the unorthodox *aqp*s subfamily previously designated *12L* is here standardized as *aqp12L1*. The expanded phylogenetic framework also prompted refinement of the *glp* nomenclature to better reflect the inferred evolutionary relationships. Accordingly, the *glp* copy broadly conserved across reef-building corals and sea anemones (formerly *glp2*) was designated *glp1*, whereas the reef-building coral-specific duplicate (formerly *glp1*) was designated *glp2*, reflecting the inferred evolutionary order of gene duplication.

**Fig. 3.**
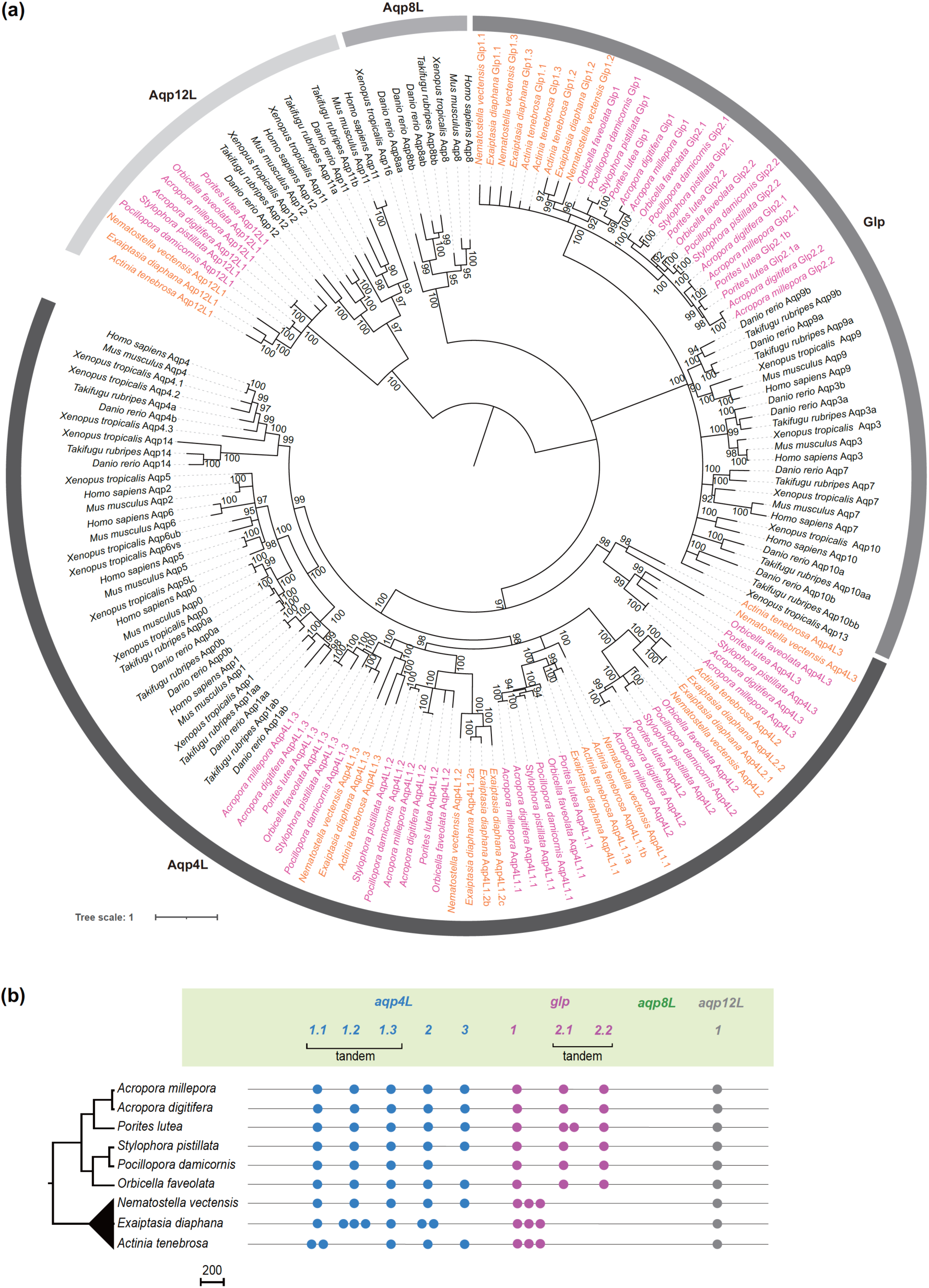
Phylogenetic relationships and gene repertoires of hexacorallian Aqps. (a) Phylogenetic relationships of Aqps reconstructed using an expanded Hexacorallia-specific dataset, including reef-building corals and sea anemones. Gene names were revised based on the phylogenetic relationships recovered in this study. All sequences used in this analysis and their corresponding accession numbers are listed in **Table 2** and **Table S1**. (b) Distribution of *aqp* orthogroups across representative hexacorallian species. The species phylogeny is shown on the left. Numbers indicate the copy number of genes assigned to each orthogroup based on the phylogenetic analysis shown in panel (a). Dashes indicate the absence of an identified ortholog.

Taken together, these results indicate that the core *aqp* repertoire comprising *aqp4L*, *glp*, and *aqp12L* genes was established in the common ancestor of Hexacorallia and has been largely conserved throughout subsequent diversification (**Figure 3b**).

### Conservation of *aqp* synteny across Hexacorallia

While **Figure 3** suggested a general similarity in *aqp* gene repertoires within Hexacorallia, the extent of conservation in their genomic organization remained unclear. To address this, we performed comparative synteny analyses and BLAST searches across representative hexacorallian genomes (**Figure 4**). Based on the phylogenetic relationships recovered in **Figure 1** and **Figure 3a**, we hypothesized that closely related reef-building corals and sea anemones would retain a highly conserved *aqp* genomic synteny despite minor lineage-specific duplication events. Detailed information on the genomic loci and accession numbers used for the synteny analyses is provided in **Tables S4-S8**.

**Fig. 4.**
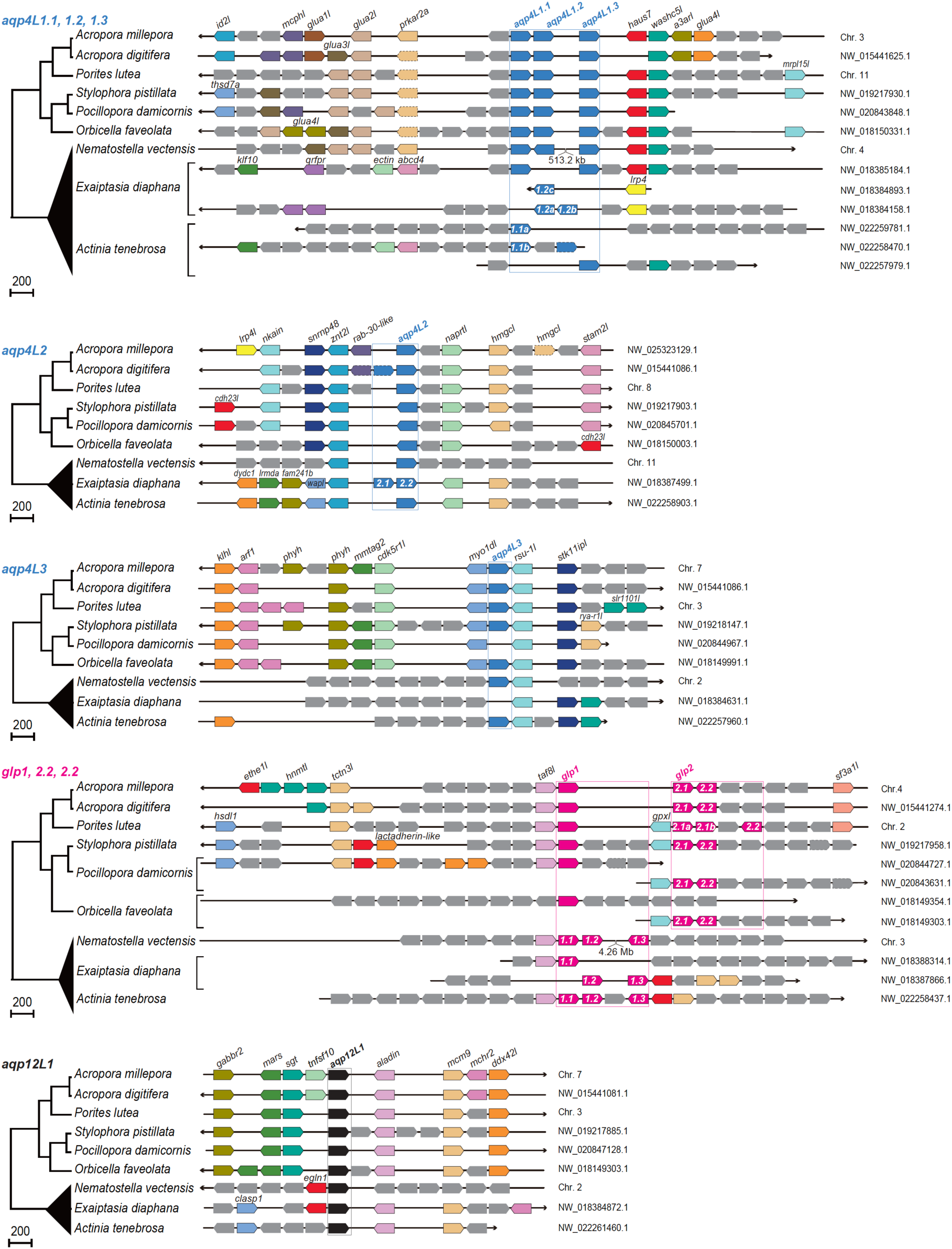
Synteny analyses of *aqp*s in Hexacorallia. Genomic regions surrounding *aqp4L*, *glp*, and *aqp12L* loci are partially conserved across representative hexacorallian species. *aqp4L* genes are shown as blue pentagonal blocks, *glp* genes as magenta pentagonal blocks, and *aqp12L* genes as black pentagonal blocks. Genes with dashed outlines represent putative pseudogenes. Colored blocks indicate genes shared among species, whereas gray blocks represent lineage-specific genes or genes of unknown function. A simplified species phylogeny is shown on the left. Accession numbers of genes are listed in **Table S4-S8**.

Comparative synteny analyses revealed partial conservation of gene order surrounding *aqp* and *glp* loci among representative hexacorallian species. Most examined genomes consistently possessed nine *aqp* genes distributed across five chromosomal loci, comprising five *aqp4L* genes (water-specific aquaporins), three *glp* genes (aquaglyceroporins), and one *aqp12L* gene (unorthodox aquaporin) (**Figure 4**). Moreover, synteny surrounding each *aqp* and *glp* locus was partially conserved across representative hexacorallian species, supporting the orthologous relationships inferred from the phylogenetic analyses. Given that our molecular phylogenetic analyses placed these reef-building coral Aqps in the same clades as their sea anemone orthologs (**Figure 1**), these findings suggest that the core composition and genomic architecture of the *aqp* family were already established in the common ancestor of Hexacorallia.

### Evolutionary conservation of exon–intron structures in *aqp* genes

To gain insights into the evolutionary lineage of the Hexacorallia *aqp* family, we examined the exon-intron structures of identified *aqp* genes and compared them with those of humans (*Homo sapiens*) (**Figure 5, Figure S1**). As a reference, exon-intron organizations of human *AQP*s were reanalyzed using publicly available genome annotations (**Figure S1**). Because reef-building corals lack *aqp8*, additional *aqp8* sequences from other metazoans were included to trace the evolutionary history of this subfamily (**Figure S2**). Consistent with previous studies, analysis of human *AQP*s revealed distinct exon-intron architectures between the classical *AQP* and *GLP* subfamilies (Lu et al. 1996; Tsukaguchi et al. 1999).

**Fig. 5.**
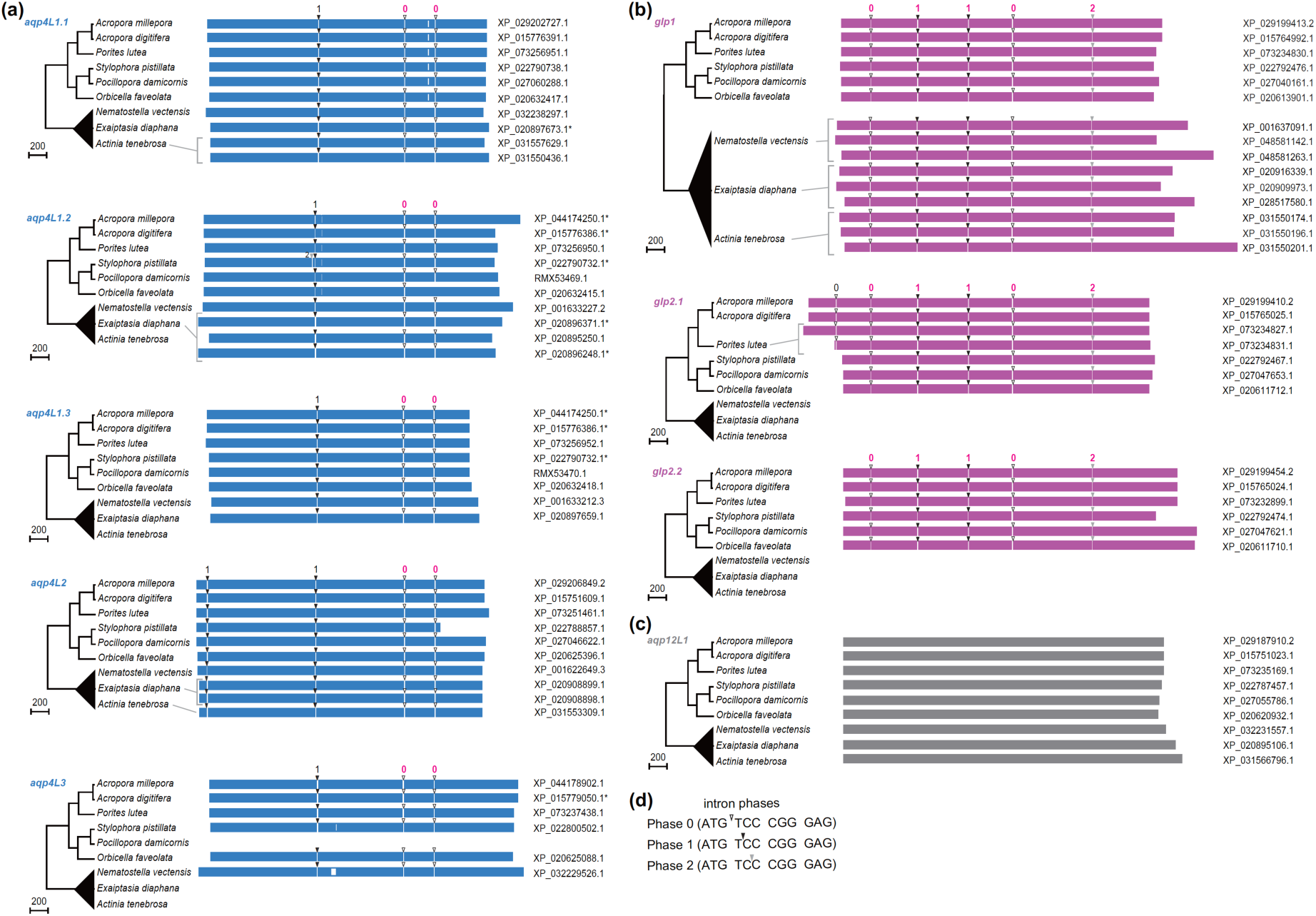
Exon–intron structures of hexacorallian *aqp* genes. Exon–intron structures of (a)*aqp4L*, (b)*glp*, and (c)*aqp12L* subfamilies. Exons are shown as colored boxes. White regions within exons represent alignment gaps. Arrowheads indicate intron positions, and numbers denote intron phases (0, 1, or 2). Colored numbers indicate intron positions and phases conserved with the corresponding human AQP subfamilies (**Fig. S1**). Alignments are shown in **Fig. S3-S5**. Species phylogenies are shown on the left. Asterisks (*) indicate full-length sequences determined in this study (**Table S1**). (d) Definition of intron phases.

Comparison of exon-intron structures revealed partial conservation between coral and human *AQP* genes (**Figure 5, Figure S1**). Within both the *aqp4L* and *glp* subfamilies, several intron positions and phases were shared between reef-building corals and humans despite the lack of clear one-to-one orthologous relationships (**Figure 5a, 5b**). In contrast, all examined coral *aqp12L* genes consisted of a single exon, whereas their human counterparts, *AQP11* and *AQP12*, contained multiple exons (**Figure 5c**). Analysis of *aqp8L*s from other metazoans further revealed that several intron positions were conserved with those of human *AQP8* (**Figure S2**). These conserved intron positions were supported by amino acid sequence alignments (**Figure S3-S6**). These findings suggest that portions of the ancestral exon-intron architecture of the *aqp* family predate the divergence of cnidarians and bilaterians and have been maintained over extensive evolutionary timescales despite substantial sequence diversification within individual *aqp* lineages.

### Functional characterization of coral *aqps*

To determine the permeability profiles of Hexacorallia Aqp4L and Glp, swelling assays were performed using *Acropora millepora* Aqp4L and Glp proteins expressed in *Xenopus* oocytes (**Figure 6**). Based on their phylogenetic positions, Aqp4L proteins were predicted to function primarily as water-specific Aqps, whereas Glp proteins were predicted to transport both water and small solutes. Consistent with these predictions, all examined Aqp4L proteins except Aqp4L1.2 exhibited water-selective permeability and showed no detectable transport of glycerol, urea, or boric acid (**Figure 6a**). Unexpectedly, however, Aqp4L1.2, which belongs to the water-specific Aqp clade, exhibited permeability not only to water but also to glycerol, urea, and boric acid. In contrast, the examined Glp proteins facilitated the transport of multiple low-molecular-weight compounds in addition to water, as predicted from their phylogenetic placement. The boric acid permeabilities were correlated with those of urea permeability (*r* = 0.995 *p* < 0.0001, **Figure 6b**). The estimated bottleneck radius of AmiAqp4L1.2 was 1.5 Å, which was slightly larger than that of AmiAqp4L1.3 (0.8 Å) (**Figure 6c**). These results indicate that solute permeability has diversified within the Aqp4L lineage and is consistent with functional co-option of a duplicated water-specific Aqp toward the transport of multiple low-molecular-weight solutes.

**Fig. 6.**
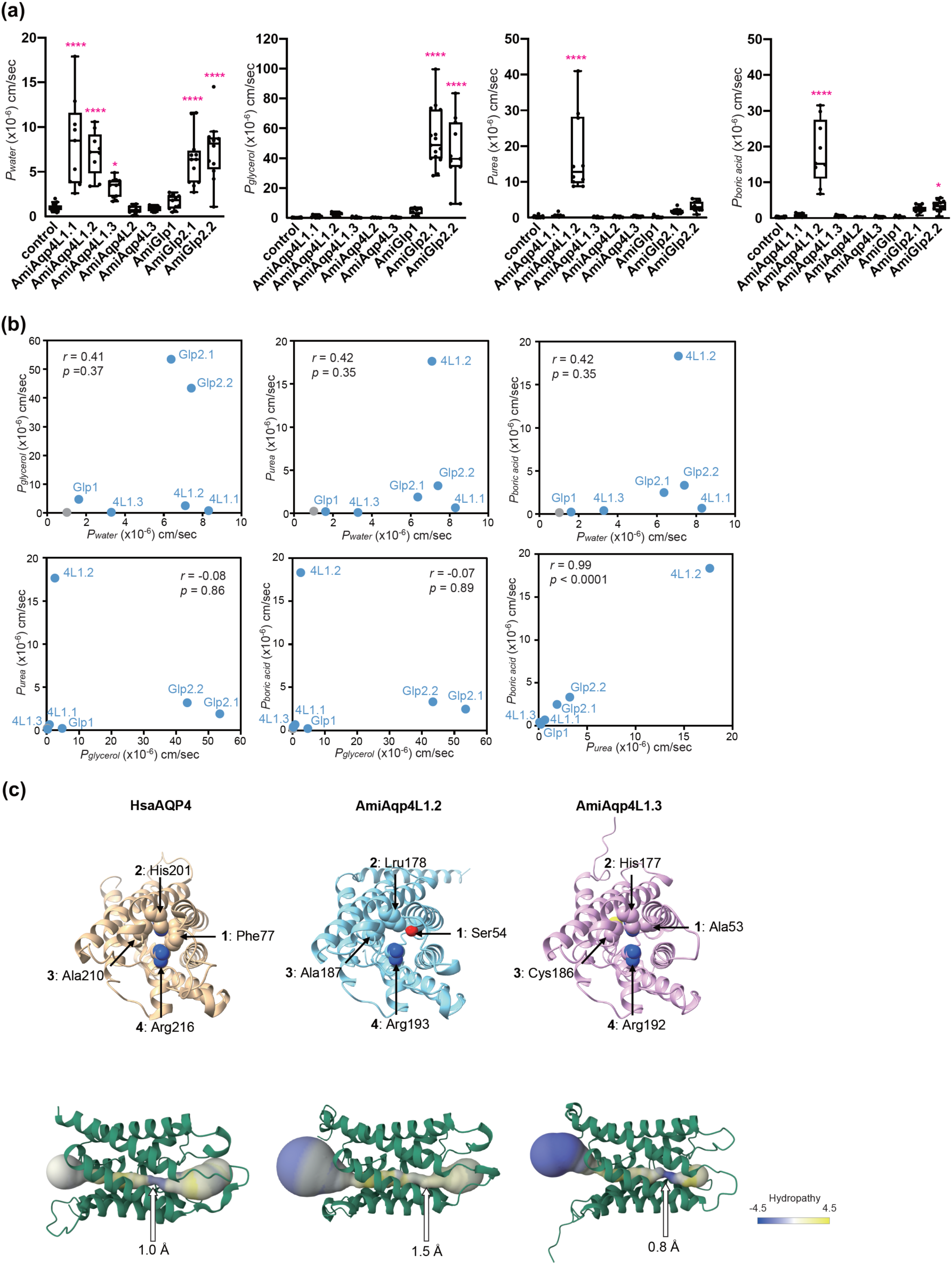
Functional divergence of solute permeabilities in AmiAqps. (a) Water (*P*_water_), glycerol (*P*_glycerol_), urea (*P*_urea_), and boric acid (*P*_boric acid_) permeabilities of *Acropora millepora* Aqps (AmiAqps) expressed in *Xenopus laevis* oocytes. Permeabilities were estimated from oocyte volume changes. Statistical significance was assessed by one-way ANOVA followed by Dunnett’s multiple-comparisons test (\**p* < 0.05, \*\*\*\**p* < 0.0001). All samples are listed in **Table S9 and S10.** (b) Scatter plots showing relationships among *P*_water_, *P*_glycerol_, *P*_urea_, and *P*_boric acid_ of AmiAqps. Water-injected oocytes were plotted in gray. Pearson’s correlation coefficients (*r*) are indicated. (c) Structural characterization of AmiAqp4L1.2 and AmiAqp4L1.3. Upper panels show the crystal structure of human AQP4 (HsaAQP4; PDB ID: 3GD8) and AlphaFold 3-predicted structures of AmiAqp4L1.2 and AmiAqp4L1.3. Residues forming the ar/R selectivity filter are shown as spheres. Lower panels show channel profiles predicted using MOLEonline. Arrows indicate the positions of the ar/R selectivity filters. Estimated bottleneck radii are shown below each channel. Channel surfaces are colored according to hydropathy.

## Discussion

Our integrative analyses of gene repertoires, synteny, exon-intron structures, and solute permeability provide a comprehensive evolutionary framework for the Aqp family in Hexacorallia. The results consistently indicate that the core hexacorallian *aqp* repertoire, comprising *aqp4L*, *glp*, and *aqp12L*, was established in the common ancestor of reef-building corals and sea anemones and has remained largely conserved throughout subsequent diversification. Conservation of synteny and exon-intron structures further supports the conclusion that the fundamental organization of the hexacorallian *aqp* family has been maintained. At a broader evolutionary scale, conservation of intron positions and phases between cnidarians and bilaterians indicates that key components of the ancestral exon-intron architecture of the *aqp* family predate the divergence of these lineages. These ancestral features have persisted over hundreds of millions of years despite extensive lineage diversification. Together, these findings indicate that the fundamental framework of the hexacorallian *aqp* family was established in the common ancestor of Hexacorallia and subsequently maintained through long-term conservation of gene repertoires, syntenic organization, and exon-intron structure.

Gene duplication is widely recognized as a major source of evolutionary innovation. Following duplication, gene copies may be retained through redundancy, dosage effects, subfunctionalization, or neofunctionalization (Kuzmin et al. 2022). Against the background of the highly conserved hexacorallian *aqp* repertoire described above, we identified evidence for localized functional innovation within the *aqp4L* lineage. In this study, *aqp4L1.2* represents a likely example of functional divergence following duplication within the hexacorallian *aqp* family. Although phylogenetically nested within the water-specific *aqp4L* clade, Aqp4L1.2 was capable of transporting glycerol, urea, and boric acid in addition to water, whereas all other examined Aqp4L proteins exhibited strict water-selective permeability. It is intriguing that a member of the Aqp4L lineage acquired the ability to permeate glycerol, urea, and boric acid because Aqp4L proteins are generally classified as classical water-specific *aqp*s. Similar evolutionary trajectories have been reported in other metazoans. For example, several insect lineages such as holometabolan insects lacking aquaglyceroporins have independently recruited Aqp4L proteins for glycerol transport, thereby compensating for the loss of the GLP subfamily (Finn et al. 2015). A comparable pattern has been proposed in mollusks. In the Pacific abalone (*Haliotis discus hannai*), HdAqp4L2 and HdAqp4L5 possess amino acid substitutions within the ar/R selectivity filter that are predicted to enlarge the pore and alter solute selectivity, resulting in aquaglyceroporin-like transport properties (Jia et al. 2022). Taken together, these observations suggest that the Aqp4L lineage possesses substantial evolutionary flexibility. Unlike holometabolan insects, reef-building corals retain multiple *glp*s. The acquisition of glycerol, urea, and boric acid permeability by Aqp4L1.2 therefore suggests that functional diversification of Aqp4L proteins can occur even in the presence of dedicated *glp*s. Interestingly, vertebrate Aqp8 proteins are known to transport urea and boric acid in addition to water (Kumagai et al. 2023; Tingaud-Sequeira et al. 2010; Ushio et al. 2022). Given that Hexacorallia lacks the *aqp8L* subfamily, the acquisition of urea and boric acid permeability by Aqp4L1.2 raises the possibility that this duplicated channel partially compensates for functions that were ancestrally mediated by Aqp8L. Although the transport properties of cnidarian Aqp8L proteins remain unknown, the overlap in substrate selectivity between vertebrate Aqp8 and Aqp4L1.2 is consistent with this hypothesis.

The physiological significance of the newly identified permeability profile remains to be fully resolved. Importantly, recent studies have demonstrated that glucose, rather than glycerol, represents the principal photosynthetically derived carbon compound transferred from *Symbiodiniaceae* to the coral host (Burriesci et al. 2012; Ishii et al. 2023). Consequently, the glycerol permeability observed in Aqp4L1.2 is unlikely to function primarily in direct symbiont-to-host carbon transfer. Instead, glycerol transport may contribute to endogenous osmotic regulation within coral holobiont. Under osmotic stress, *Symbiodiniaceae* accumulate the compatible osmolyte floridoside. Because floridoside is a galactosyl-glycerol compound, its synthesis depends on intracellular glycerol metabolism. The glycerol permeability of Aqp4L1.2 therefore raises the possibility that glycerol transport represents an important component of osmotic acclimation within the coral holobiont. Similarly, the observed permeability to urea may facilitate nitrogen recycling within the coral holobiont, while boric acid permeability raises the possibility that Aqps participate in the movement of boron-containing compounds in marine environments. Although the physiological significance of boric acid transport remains unclear, these findings substantially expand the range of candidate functions attributable to coral Aqps.

Our results reveal a striking contrast between evolutionary conservation and functional innovation within the hexacorallian *aqp* family. The remarkable conservation of *aqp* repertoires, syntenic organization, and intron positions across deeply divergent metazoan lineages indicates that the core organization of the *aqp*s family has been retained over extensive evolutionary timescales, while local gene duplications have repeatedly generated opportunities for functional innovation. At the same time, the emergence of a multifunctional Aqp4L protein illustrates how localized changes can generate substantial physiological innovation without major alterations to overall family architecture. The hexacorallian *aqp*s family therefore exemplifies a general evolutionary principle: deep conservation of gene-family architecture can coexist with repeated functional innovation at individual loci through gene duplication and co-option, highlighting the remarkable evolutionary flexibility of the Aqp4L lineage across metazoans.

## Materials and Methods

### Phylogenetic and Synteny Analyses

To examine the evolutionary relationships of Aqps across cnidarians and basal metazoans (**Figure 1**), amino acid sequences were aligned using MAFFT v7.526 (Katoh and Standley 2013) with the E-INS-i algorithm. Multiple sequence alignments were trimmed using trimAl v1.4 (Capella-Gutiérrez et al. 2009) with the gap-threshold option (-gt 0.2). Maximum-likelihood (ML) phylogenetic analyses were conducted using IQ-TREE 3 (Nguyen et al. 2015) under the LG+C20+F+R10 model. Branch support was assessed using 1,000 ultrafast bootstrap (UFBoot) replicates (Hoang et al. 2018; Minh et al. 2013) and 1,000 SH-like approximate likelihood ratio test (SH-aLRT) replicates (Guindon et al. 2010). To further resolve relationships within cnidarian Aqps (**Figure 3**), a second ML phylogenetic analysis was performed using a cnidarian-specific dataset. Amino acid sequences were aligned using MAFFT v7.526 and trimmed with trimAl v1.4 using the -gt 0.2 option. The best-fitting substitution model was selected using ModelFinder (Kalyaanamoorthy et al. 2017) based on the Bayesian Information Criterion (BIC), resulting in selection of the Q.PFAM+R7 model. ML phylogenetic inference was then performed using IQ-TREE 3 with 1,000 UFBoot replicates. All phylogenetic trees were visualized and annotated using iTOL v7 (Letunic and Bork 2024).

Synteny analyses were performed using chromosome-level or scaffold-level genome assemblies of representative hexacorallian species and other cnidarians listed in **Table 1**. The genomic regions surrounding *aqp* loci were manually examined to compare gene order, orientation, and local genomic neighborhoods among species. Orthologous relationships inferred from phylogenetic analyses were further evaluated by comparison of conserved syntenic relationships and BLAST searches.

### Cloning of Acropora millepora Aqps

Full-length cDNAs encoding *Acropora millepora* Aqps (AmiAqps) were chemically synthesized (**Table S1**; Eurofins Genomics, Tokyo, Japan) with 15 bp terminal sequences homologous to the ends of the linearized pGEMHE vector (Liman et al. 1992). The synthesized cDNAs were subcloned into pGEMHE using the NEBuilder HiFi DNA Assembly Master Mix (New England Biolabs, Ipswich, MA, USA). All constructs were verified by Sanger sequencing using an Applied Biosystems 3730xl DNA Analyzer (Thermo Fisher Scientific, Waltham, MA, USA).

### Expression of Aqps in *Xenopus laevis* Oocytes

All aquatic animal procedures were conducted in accordance with the National Institutes of Health “Guide for the Care and Use of Laboratory Animals” and in accordance with a manual approved by the Institutional Animal Experiment Committee of the Institute of Science Tokyo. Adult female *Xenopus laevis* were purchased from Watanabe Zoushoku (Hyogo, Japan). cRNA synthesis, oocyte isolation, microinjection, and incubation were performed as previously described (Kumagai et al. 2023; Ushio et al. 2022). Briefly, stage V–VI *Xenopus laevis* oocytes were injected with either nuclease-free water (control) or cRNA encoding AmiAqps (0.5 ng/nL; 25 ng per oocyte) and incubated in OR3 medium at 16°C for 5 days prior to functional analyses.

### Swelling Assay and Permeability Calculations

Osmotic water and solute permeability assays were performed using the *Xenopus* oocyte swelling method as previously described (Imaizumi et al. 2024; Kumagai et al. 2023). Oocyte volume changes were monitored using a stereomicroscope (SZX9, Evident, Tokyo, Japan) equipped with a CCD camera (DS-Fi2, Nikon, Tokyo, Japan). Oocyte volumes were estimated assuming spherical geometry. Prior to each assay, oocytes were equilibrated in ND96 solution (approximately 200 mOsmol/kg, pH 7.5). Water permeability (*P*_water_) was determined by transferring oocytes to hypotonic ND96 diluted two-fold with distilled water (approximately 100 mOsmol/kg). Glycerol, urea, and boric acid permeabilities were assessed using iso-osmotic substitution assays in which NaCl was replaced with 180 mM glycerol, urea, or boric acid while maintaining a total osmolality of approximately 200 mOsmol/kg. Permeability coefficients were calculated from the initial rates of volume increase. Osmotic water permeability (*P*_water_) and solute permeability (*P*_solute_) were calculated according to the methods of Preston et al. (1992) and Carbrey et al. (2003), respectively (Carbrey et al. 2003; Preston et al. 1992). Each experiment was performed using oocytes obtained from at least three independent *Xenopus laevis* individuals.

### Statistical Analysis

Statistical analyses were performed using GraphPad Prism version 8 (GraphPad Software, San Diego, CA, USA). Statistical significance was evaluated using one-way ANOVA followed by Dunnett’s test (\*\*\*\**p* < 0.0001, \**p* < 0.05).

### Structural Modeling and Pore Analysis

Structural models of AmiAqp4L1.2 and AmiAqp4L1.3 were generated using AlphaFold 3 (Abramson et al. 2024). Structural models were visualized using UCSF ChimeraX version 1.9 (Pettersen et al. 2021). The crystal structure of human AQP4 (PDB ID: 3GD8) was used as a reference (Ho et al. 2009). Pore analyses were performed using MOLEonline (Racek et al. 2025). For AmiAqp4L1.2, residues 233–262 were removed prior to analysis because the predicted structure contained an extended C-terminal region that interfered with pore visualization. AmiAqp4L1.3 was analyzed using the full-length model.

## Data Availability

Data generated/analyzed in this study are available in this article and Supplementary Materials.

## Acknowledgments

We would like to thank Nodoka Nagai, Yoko Yamamoto, Nana Shinohara, the Bioscience Center, and the Open Research Facilities for Life Science and Technology at the Institute of Science Tokyo for their technical assistance. This work was supported by the Japan Society for the Promotion of Science (JSPS) KAKENHI, grant number 24K09067 (to A.N.), the Laboratory for Design of Social Innovation in Global Networks (DLab) Challenge Research Grant from the Institute of Science Tokyo (to A.K.), and the Core Research for Evolutionary Science and Technology (CREST) Program on Carbon-Blue X from the Japan Science and Technology Agency (JST), grant number JPMJCR23J2 (to A.K.).

## Author Contributions

A.N. and A.K. conceived and designed the study, interpreted the experimental results, and drafted and edited the manuscript. A.N. and A.K. performed the experiments. A.N. and A.K. analyzed the data. A.N. and A.K. approved the final version of the manuscript.

