## Supplementary material for "Solute selectivity and functional divergence of aquaporins in reef-building corals": TableS1-S9

Table S1. Coding and amino acid sequences of aquaporins analyzed/identified in this study.

| **Species** | Gene name | Accezsion no. | Coding sequence | Amino acid sequence |
|---|---|---|---|---|
| *Acropora millepora* | *aqp4L1.2* | XM_044318315.1<br>XP_044174250.1 | ATGTTGTTCCAAGAAGGTAGCAACGAATTTCTACGCCGCTCATTTTGGACCGAGGTGTTCGCAGAGTTTTTAGT<br>AACACTCATTTACTTGTTCCTGGTTTGCGGTGCAGTGTTGCCGTGGAATAATACGCCGCCTCCCGTGATTCACGG<br>CGCCTTCATCAGCGGTTTGAGCATCGCCACCTTAGTAATGGCGGTTATGCACATTTCTGGTGGACATCTCAACCC<br>AGTCGTAACCATTTCATTCATGTCTGTTGGAAGGGTATCAGTATTGAAGGCTATTTTCTTTATAGTAGCCCAATGT<br>TTAGGAGCGATATGTGGTGCGGCCATGGTGTATGGTACAACTCCCGAAGAAGTCATCGGTTCCTTGGGAGTGAC<br>GGCTCCTGGAAAAGGCGTTACCACAATGCAGGCGTTTGCGGTCGAGCTTGGCCTCACATTCGTACTGATGTTTT<br>TTGTCATGGCTGTTACGGACCCCACAAGAGGCATGACCGGTTATGGAGTACCGCTTTCCATTGGAATTTGTGTG<br>TTCGTTTGCCTTATGCAGGGGATTCCGTCATCTGGGGCCAGCTTGAATCCTGCGCGTTCCATCGGACCTGCTGTT<br>GTCATGAATTCATGGAGGGATCATTGGATTTACTGGGCTGCTCCAACAGCTGGTGGTCTCCTAGGAACGGGCAC<br>TTATCAACTGGTATTCGCTTCGAAAAGATCCAATCACGATTCAGATAATAACATGATTCATTCTATGGAAGTGAC<br>GGAGAAGGATGATGCTACCAAATTCCTTGATTCCACTTTGTGA | MLFQEGSNEFLRRSFWTEVFAEFLVTLI<br>YLFLVCGAVLPWNNTPPPVIHGAFISGL<br>SIATLVMAVMHISGGHLNPVVTISFMSV<br>GRVSVLKAIFFIVAQCLGAICGAAMVYG<br>TTPEEVIGSLGVTAPGKGVTTMQAFAVE<br>LGLTFVLMFFVMAVTDPTRGMTGYGVP<br>LSIGICVFVCLMQGIPSSGASLNPARSIG<br>PAVVMNSWRDHWIYWAAPTAGGLLGT<br>GTYQLVFASKRSNHDSDNNMIHSMEVT<br>EKDDATKFLDSTL |
| *Acropora millepora* | *aqp4L1.3* | XM_044318315.1<br>XP_044174250.1 | ATGCCAGTTTTTGACACGCTCGCGCTGAGAAAGCGCGAATTTTGGTCCGCCGTCGGTGGTGAATTTATCGCCAC<br>ACCATTTTTGTATTTATGGTTTGTGCTTCGACTTTAACCTGGGAAAACACCTCTTGTGCACTCCTCCGTATTTCGC<br>GACTGCTGGGATGACCATTGCGAGCTTAGCTTTGACCATTGGTCACCTGAGTGGTGGTTTAATGAATCCAGCCG<br>GAACATCGCATTTCTGGTTTCGCGAAGAATTACATTAATTGAAGGAGCTTTCTTCACTGTGGCTCAATTTTGCGG<br>GGCATTTTAGGTGCAGCTCTTTTATACGGTCTTACACCCTCGATGGTGAGAGGAGGACTGGGAGTGACAGCACC<br>GCACACAATCTTAATAATGCCCAGGCCTTTGGAATTGAGCTGTTACTCACGTTTGTACTGGTTCTGACCATATTT<br>GCGCGACAGACCGCGGCAAGAATCACCACGGATATGAAATACCGTTGTCCATTGGGACATGTGTACTCATCTGT<br>ACATGGTCGGGATTCCATTTACCGGCTGCAGTATGAATCCTGCGCGGTCGTTTGGACCGGCCCTAATTATGAACT<br>TTGGAAACAACATTGG | MPVFDTLALRKREFWSAVGGEFIATTIF<br>VFMVCASTLNWENTSCALLRISLTAGM<br>TIASLALTIGHLSGGLMNPAVNIAFLVSQ<br>RITLIEAAFFTVAQFCGGILGAALLYGLT<br>PWMVRGGLGVTAPAHNLNNAQAFGIE<br>LLLTFVLVLTIFGATDRGKNHHGYEIPLS<br>IGTCVLICHMVGIPFTGCSMNPARSFGP<br>ALIMNSWKQHWVYWVGPIPGAIMGAL<br>LYEHVFSASKMAVTPA |
| *Acropora digitifera* | *aqp4L1.2* | XM_015920900.1<br>XP_015776386.1 | ATGTTGTTCCAAGAAGGTAGCAACGAATTTCTACGGCGTTCATTTTGGACCGAGGTGTTCGCAGAGTTTTTGGT<br>AACACTCATTTACTTGTTCCTGGTTTGCGGTGCAGTGTTGCCGTGGAATAACACGCCGCCTCCCGTGATTCACG<br>GCGCCTTCATCAGCGGTTTGAGCATCGCCACCTTAGTTATGGCGGTTATGCACATTTCTGGTGGACATCTCAACC<br>CAGTCGTAACCATTTCATTCATGTCTGTTGGAAGGGTATCAGTATTGAAGGCTATTTTCTTTATAGTAGCCCAGTG<br>TTTAGGAGCGATATGTGGTGCGGCCATGGTGTATGGTACAACTCCCGAAGAAGTCATCGGTTCCTTGGGAGTGA<br>CGGCTCCTGGAAAAGGCGTTACCACAATGCAGGCGTTTGCGGTCGAGGTTGGCCTCACATTCGTACTGATGTTT<br>TTTGTCATGGCTGTTACGGACCCCACAAGAGGCATGACCGGTTATGGAGTACCGCTTTCCATTGGAATTTGTGT<br>GTTCGTTTGCCTTATGCAGGGGATTCCGTCATCTGGGGCCAGCTTGAATCCTGCGCGTTCCATCGGACCTGCTG<br>TTGTCATGAATTCATGGAGGGATCATTGGATTTACTGGGCTGCTCCAACAGCTGGTGGGCTCCTAGGAACGGGC<br>ACTTATCAACTGGTATTCGCTTCGAAAAGATCCAATCACGATTCAGATAATAACATGATTAATTCTATGGAAGTG<br>ATGGACAAGGATGATGCTTCCAAATTCCTTGATTCTACTTTGTGA | MLFQEGSNEFLRRSFWTEVFAEFLVTLI<br>YLFLVCGAVLPWNNTPPPVIHGAFISGL<br>SIATLVMAVMHISGGHLNPVVTISFMSV<br>GRVSVLKAIFFIVAQCLGAICGAAMVYG<br>TTPEEVIGSLGVTAPGKGVTTMQAFAVE<br>VGLTFVLMFFVMAVTDPTRGMTGYGV<br>PLSIGICVFVCLMQGIPSSGASLNPARSI<br>GPAVVMNSWRDHWIYWAAPTAGGLLG<br>TGTYQLVFASKRSNHDSDNNMINSMEV<br>MDKDDASKFLDSTL |
| *Acropora digitifera* | *aqp4L1.3* | XM_015920900.1<br>XP_015776386.1 | ATGCCCGTTTTTGACACGCTTGCGCTGAGAAAGCGCGAATTTTGGTCCGCCGTCGGTGGTGAATTTATCGCCAC<br>TACCATTTTTGTGTTTATGGTTTGTGCTTCGACTTTAACCTGGGAAAACACCTCTTATGCACTTCTCCGTACTTCG<br>CTGACTGCTGGGATGACCATTGCGAGCTTAGCTTTGACGATTGGTCACCTGAGTGGTGGTTTAATGAATCCAGC<br>CGTGAACATCGCATTTCTGGTTTCGCGAAGAATTACATTAATTGAAGGAGCTTTCTTCACTGTGGCTCAATTTTG<br>CGGGGGCATTTTAGGTGCAGCTCTTTTATACGGACTTACACCCTCGATGGTGAGAGGAGGACTGGGAGTGACA<br>GCACCAGCACACAACCTTAATAATGCCCAGGCCTTTGGAATTGAGTTGTTACTCACGTTTGTACTGGTTCTGAC<br>CATATTTGGCGCGACAGACCGCGGCAAGAATCACCACGGATATGAAATACCGTTGTCCATTGGGACATGTGTAC<br>TCATCTGTCACATGGTCGGGATTCCATTTACCGGCTGCAGTATGAATCCTGCACGGTCGTTTGGACCGGCCCTG<br>ATTATGAACTCTTGGAAACAACATTGGGTTTACTGGGTTGGCCCTATTCCAGGTGCAATAATGGGTGCGCTGCT<br>CTATGAGCACGTTTTTTCTGCTAGCAAGATGGCTGTGACTCCAGCTTAA | MPVFDTLALRKREFWSAVGGEFIATTIF<br>VFMVCASTLTWENTSYALLRTSLTAGM<br>TIASLALTIGHLSGGLMNPAVNIAFLVSR<br>RITLIEGAFFTVAQFCGGILGAALLYGLT<br>PSMVRGGLGVTAPAHNLNNAQAFGIEL<br>LLTFVLVLTIFGATDRGKNHHGYEIPLSI<br>GTCVLICHMVGIPFTGCSMNPARSFGPA<br>LIMNSWKQHWVYWVGPIPGAIMGALL<br>YEHVFSASKMAVTPA |
| *Acropora digitifera* | *aqp4L2* | XM_015896123.1<br>XP_015751609.1 | ATGGCGCGCACATTTCGTTATCTTCGAGTTGGACTGAATGAAGCATTTCATACAGACTTCTGGTTAGCCATTTTG<br>GCCGAGTATCTTGCTACATTGTTTCTCTTGTTCGCTGTATCCGGTTCGTCGCTTCGATGGGAAGGAGCCCCAACA<br>ATTTTCCAACAAGCTCTCGCTGCAGGGCTTTCCACTGCATCTGCTATTCAAGCATTCAGATCAGTTAGCTTGCCC<br>CTTGTACATGGGAATCCAGCCGTTTCGCTCGCAGCGTTCATCACCGGGGATGCTAGCTTGGTTCGCGTAACTTG<br>CTCCATCATCGCGCAGATGTTTGGAGCGATAACCGGTGCGGGAATCATAATGGCAATCAGTCCAGAACACGCCC<br>GTGGGAAGCTTGGCGCGACTATTCCAGCAAAAGACGTGAGCGGTGCCCAGGCCTTCGGGGTTGAGCTCGTTG<br>CATCATTCTTACTGGTACTAACAATGCTGGCCTCTATTGAAGCGAACAAAGACGCCGCTCGAAGGGATTATGGG<br>ACAGCTGCAGCGTTGGGTGTGGTAGTTACTCTGTGCTACCTTATCGCGTTGCCAATCACCGGTTGTGGCGTGAA<br>TCCTGCGAGGTCTTTTGCTCCAGCGGTTTTAGCAGACACCTGGAAAGAACACTGGGTTTACTGGGCTGGGCCG<br>CTAATGGCATCGTTGTTGGCAGGATTCCTTCATTGTTTGTTGTTCAAAAATGCTGAACGTCGCCATGAACGTTCT<br>GCTAGCACGGAAAACGAGCTTGATGAGGCCATTATTCAGTAA | MARTFRYLRVGLNEAFHTDFWLAILAE<br>YLATLFLLFAVSGSSLRWEGAPTIFQQA<br>LAAGLSTASAIQAFRSVSLPLVHGNPAV<br>SLAAFITGDASLVRVTCSIIAQMFGAITG<br>AGIIMAISPEHARGKLGATIPAKDVSGA<br>QAFGVELVASFLLVLTMLASIEANKDAA<br>RRDYGTAAALGVVVTLCYLIALPITGC<br>GVNPARSFAPAVLADTWKEHWVYWAG<br>PLMASLLAGFLHCLLFKNAERRHERSA<br>STENELDEAIIQ |
| *Acropora digitifera* | *aqp4L3* | XM_015923564.1<br>XP_015779050.1 | ATGGGACGTTGGACTGAGGCCTTGGTCGTTTTTGTCCCCTGTTTGGTTGAATTAGTCTGCACAATATTTCTTGTT<br>TTTTTGGGATGCGGCTCGACCTTAAATTGGCCAAGAGAAGAAACTTCTCCAACTGTCTTAACCATATCGTTGTC<br>TTTTGGGTTCTTGTATGCATTTCTGGGGAACATCTCCACTCTTGTCAGTGGTGGTTTTTTTAATCCTGCAATAACA<br>GCGGCTTTGGCGACGATAAAAACTATTCCAATCAAAAGAGCGGTCTGCTTGATTTCGCGCAACTGACAGGAG<br>CTGTTATTGGAGCAGGACTCCTGTACTTCGTCGTCCCTGATGACAATCGTGAGTATTTGGGAGTGACGCGAATG<br>AGGTCAGTCTCTTGTCTCACTGGATGTGGCGTGGAAGTATTGATGACCGCAGCTTTGACGCTAACTGTGTTGCT<br>AACATCTGATGGAGGCCAAAGAACTATCGATTCTTCAGCATGCATGGTGATAGGGATGGCAGTTAGTGCATGCC<br>ACTTGTTTGGGTACCCTATAAGTGGATGTGGTATCAACCCAGTGCGTTCTTTTGGACCTGCTGTCATTATGAACA<br>TATGGAAAAATCACTGGGTGGTTTATTGGATTGGTCCAATCATTGGTGCACAACTGGCAGCAGCCATTCATCATC<br>TCGTCAACAATTTTCCAGTTAAAACAGCTGACGAGCTTGAAGTCAGGGGAAGACAACTCAAAGACAAAGTGG<br>AAATGCCATCTTTTGGATTGAACATTAGTGAACACCACCCCCCAATTCCAACCCAGGTTACTACACAAGGTGAC<br>TCTCCCAGTGAGCTTGCTACTTTCCTGTGA | MGRWTEALVVFVPCLVELVCTIFLVFLG<br>CGSTLNWPREETSPTVLTISLSFGFLYAF<br>LGNISTLVSGGFFNPAITAALATIKTIPIK<br>RAVCLIFAQLTGAVIGAGLLYFVVPDDN<br>REYLGVTRMRSVSCLTGCGVEVLMTA<br>ALTLTVLLTSDGGQRTIDSSACMVIGMA<br>VSACHLFGYPISGCGINPVRSFGPAVIMN<br>IWKNHWVVYWIGPIIGAQLAAAIHHLV<br>NNFPVKTADELEVRGRQLKDKVEMPSF<br>GLNISEHHPPIPTQVTTQGDSPSELATFL |
| *Acropora digitifera* | *glp1* | XM_015909506.1<br>XP_015764992.1 | ATGCCAAGTAGATTTCAGGGAAGACTTTCGCCTTTGTCGAGAGAAGCACTTGCCGAATTCATTTCTACATTTATT<br>TTAGTGTCATTTGGAGTTGGATCAGTTGCCCAGAAGGTCCTCAGCCACGGAGAATTTGGCACCTTCTTTTCCAT<br>AAACTTCGGCTGGGGTATTGGGGTTACCCTCGGGTGTTATTGGGCCGGAGGGGTATCAGGTGCTCACATGAATC<br>CTGCAGTTACTCTCGCGTCTGCTGTTGTTCGGCGTTTGGACTGGGTGAAGGTTCCCGTGTATTGTATTGCTCAGT<br>TCTTTGGAGCCTTCATCGCGTCTGCAGTTGTCTATGGAATCTACCATGATTCTCTGAATGCCTTTGACGGTGGGG<br>TTCGATACGTGTTGGGTAAAAAAGGTACTGCAGGGATCTGGGCCACCTATCCCCAGGCCCATGTATCCACAAGT<br>AATGGATTCGCGGATCAGCTGTTTGGCACAGCCCTTCTCGTCAGTTGTATTTTTGCCATCTTGGACCGAAATAAC<br>AACGCACCAAACAAGGGAGTGGCTCCGGTCATGATCGGTTTGGTGGTCTTTGTGATTGGCACCTCTTTTGGCTC<br>CAACTGCGGCTACGCCATAAATCCTGCCAGGGATCTGGGACCAAGGGTATTTACTGCCATGGCAGGGTGGGGT<br>GGCGAGGTGTTCACGGCTGCCAATCATTGGGCTTGGGTACCAGTTGTTGCTTGTTCGCTTGGAGGAGTACTGG<br>GTGCCATTGTGTATATGATTTTTATAGAGCTGCATCATCCATTGACGAATGATCAGCATAACTCTACAGACAATCA<br>ATACCTGCCAATCAATCCCAGAGAGGACGATGAAGATGAACGTAGTGCAACAGTGTAA | MPSRFQGRLSPLSREALAEFISTFILVSF<br>GVGSVAQKVLSHGEFGTFFSINFGWGIG<br>VTLGCYWAGGVSGAHMNPAVTLASAV<br>VRRLDWVKVPVYCIAQFFGAFIASAVV<br>YGIYHDSLNAFDGGVRYVLGKKGTAGI<br>WATYPQAHVSTSNGFADQLFGTALLVS<br>CIFAILDRNNNAPNKGVAPVMIGLVVFV<br>IGTSFGSNCGYAINPARDLGPRVFTAMA<br>GWGGEVFTAANHWAWVPVVACSLGG<br>VLGAIVYMIFIELHHPLTNDQHNSTDNQ<br>YLPINPREDDEDERSATV |
| *Acropora digitifera* | *glp2.1* | XM_015909539.1<br>XP_015765025.1 | ATGTCTAACAATATGGATAAAGATAAAACTGCGGAGAAGCTCGATAAAGATGACAATATCGATAAACTTCAACA<br>ATGTAAGCAAGTTTTTTCCATGGCTAAGCGATTTCGTCTATCTCCCTTGATGAGGGAAGCCGTCGCTGAGTTTTG<br>TGGTACCTTCATTCTACTGATATTTGGCATAGCTTCTGTGGCGCAGATGGTGCTAAGCAATTGGAAGTTTGGAAA<br>CTTTTTCTCCGTCAATTTCGGCTGGGGTATTGGTGTCACCTTGGGGTGTTACTGGGCCGGCGGTGTATCAGGCG<br>CTCACATGAATCCAGCAGTTACTGTCGCGTTTGCGGTTGTGAAGAGATTACCCTTGAGAAAAGTTCCTGTGTAT<br>GTTGCAGCACAAATGCTCGGTGCGTTTGTAGCTTCAGCCTGCGTTTATGGAGTGTATTACGATGCCCTCAATGC<br>ATTTGATGGAGGCGTCCGCCAAGTCCTGGGTCACAAGGGAACAGCTGGGATCTGGGCAACATACCCACAGGA<br>ATACTTATCAATCGGCAGCGGTTTCGCTGACCAGTTCCTTGGCACAGCTCTGCTAGTTGGATGTATTTTCGCTAT<br>CACTGATAAAAAGAACAACTCTCCTAACAAGGGTGTTGCACCTGTCATGATCGGTCTGGTGGTTTTTGTCATCG<br>GAGCAACCTTTGGATTCAACTGCGGCTATGCTATTAACCCTGCCAGGGATCTGGGACCAAGAATCTTCACTGCA<br>ATGGCAGGGTGGGGTGGTGAGGTGTTCACGGCGGGTAATAACTGGTCCTGGGTTCCAGTGATCGCATGCCTCA<br>TGGGAGGTGTCACAGGTGCCTTTATCTACATCGGTCTTATCGAGATGCACCATGATTCAGATGAACCAAACGAA<br>AACGGGAGAGAACTGGAGAGGATTATCATAAATGAATAA | MSNNMDKDKTAEKLDKDDNIDKLQQC<br>KQVFSMAKRFRLSPLMREAVAEFCGTFI<br>LLIFGIASVAQMVLSNWKFGNFFSVNFG<br>WGIGVTLGCYWAGGVSGAHMNPAVTV<br>AFAVVKRLPLRKVPVYVAAQMLGAFVA<br>SACVYGVYYDALNAFDGGVRQVLGHK<br>GTAGIWATYPQEYLSIGSGFADQFLGTA<br>LLVGCIFAITDKKNNSPNKGVAPVMIGL<br>VVFVIGATFGFNCGYAINPARDLGPRIFT<br>AMAGWGGEVFTAGNNWSWVPVIACL<br>MGGVTGAFIYIGLIEMHHDSDEPNENG<br>RELERIIINE |
| *Acropora digitifera* | *glp2.2* | XM_015909538.1<br>XP_015765024.1 | ATGGCCATTCGTTTTCGATTGTCTACTCTACAAAAAGAAGCACTCGCTGAGTTCTGCTCTACATTTATCTTGGTG<br>ATATTTGGGGTAGGCTCAATTGCACAGATGGTGCTAAGTGAGTGGAAGTTTGGGAACTTTTTTTCCGTCAATTTT<br>AGCTGGGGAATTGGTGTGACATTAGGCTGTTACTGGGCCGGCGGAATCTCAGGCGCCCACATGAACCCTGCGG<br>TGACTCTTGCCTTTGCTGTGACGAGAAGACTCGCTTGGAAGAAAGTTCCCGTATATTGGGCGGCGCAAATGGT<br>TGGCTCTTTTGTGGCCTCGGCTTGTGTCTACGGAGTATACTACGATGCCCTGAGTGCATTTGATGGTGGCCATCG<br>TCAAGTATTGGGTCCAAAAGGCACCGCTGGAATATGGGCGACGTATCCTAAACCCTTTTTGTCAATAGCGAGTG<br>GCATCGCCGACCAGGTTCTCGGAACTTCTTTGCTTGTGAGCTGTATCTTCGCGATCAACGATAAAAATAATAATT<br>CCCCCGACCAGGGCGTGGCGCCCGTTGTGATAGGGTTGGTGGTGTTTGTTATTGGTGCAACGTTTGGATTTAAC<br>TGCGGCTACGCCATCAATCCCGCAAGGGATCTGGGACCAAGATGTTTCACTGCTATTGCAGGGTGGGGCATTGA<br>AGTGTTTACAGCTGCCAATAACTGGTGGTGGGTCCCTGTGGTTGGCGGTTTCCTGGGCGGCGTTATTGGAGGAC<br>TAATTTATGTATGTTTTATTGGAATGCACCTCAGTTCTGATAGCGGAGGGGAGTCAGGGTATCAACTTGAAAATA<br>TCATCACGACCAAGGCTGAGTTTAAGACCGACGATTCTGAAATGGGCAAGGGAAACGTGGCTTTTAGCGATGT<br>AGTCGTAGCTAATGTAAAATAA | MAIRFRLSTLQKEALAEFCSTFILVIFGV<br>GSIAQMVLSEWKFGNFFSVNFSWGIGV<br>TLGCYWAGGISGAHMNPAVTLAFAVTR<br>RLAWKKVPVYWAAQMVGSFVASACV<br>YGVYYDALSAFDGGHRQVLGPKGTAGI<br>WATYPKPFLSIASGIADQVLGTSLLVSCI<br>FAINDKNNNSPDQGVAPVVIGLVVFVIG<br>ATFGFNCGYAINPARDLGPRCFTAIAGW<br>GIEVFTAANNWWWVPVVGGFLGGVIG<br>GLIYVCFIGMHLSSDSGGESGYQLENIIT<br>TKAEFKTDDSEMGKGNVAFSDVVVAN<br>VK |
| *Stylophora pistillata* | *aqp4L1.2* | XM_022934997.1<br>XP_022790732.1 | ATGTTTCTCCGCGATGGGAAAATGGAATTCCGCTCTCGCAAATTGTGGACCGAAGTCTTCGCGGAATTTTTGGC<br>CACACTTTTGTTCCTGTTCATGGTCTGTGGCACAGTGTTACCGTGGAATAACACACCACCATCCGTTATACACAT<br>TGCTTTCAGTCATGGACTAAGTATAGCGACCTTAGCGATGGCGGTGTTTCACATTTCGGGAGGACAGCTTAACC<br>CAGCCGTGACCATCAGTATGATGGCAGTTGGAAGGGTTTCAGTCTTAAAAGCTGTGTTTTTCATCATTGCTCAC<br>GAAGCTATTTGTGGAGCAGCTATAGTTTTCTACCTAACCCCAAATGACTCTGTTGGTTCATTGGGTGTTACAGCT<br>CCTTCAAATGGTGTCACTACTGCGCAGGCGTTTGGAGTGGAACTCATGTTGACTTTCATATTGGTTTTTGTGATA<br>TTCGCTGCGACTGACCCCAACAGAGACATGGCAGGATATGGGGTACCGCTGGCCATAGGAATTTGTGTGTTCAT<br>TTGTCTGATGCATGGGATCCCTTCATCTGGAGCCAGTTTGAATCCAGCTCGTTCCCTTGGTCCTGCCGTTGTTAT<br>GAATTTGTGGAGAGACCATTGGATTTACTGGGTGGCACCTATTGTTGGCGGTCTCCTAGCAACGTGCACCTACC<br>AGCTGTTTTTCGCCCGGAGGAGAATCAACCAAAACACTTTGGAAGATGACAACTACAATGTGGAACTCAATGA<br>CAACGATACATCGAAGTT TATGGTCTCCCCTTTGTAA | MFLRDGKMEFRSRKLWTEVFAEFLATL<br>LFLFMVCGTVLPWNNTPPSVIHIAFSHG<br>LSIATLAMAVFHISGGQLNPAVTISMMA<br>VGRVSVLKAVFFIIAHEAICGAAIVFYLT<br>PNDSVGSLGVTAPSNGVTTAQAFGVEL<br>MLTFILVFVIFAATDPNRDMAGYGVPLA<br>IGICVFICLMHGIPSSGASLNPARSLGPAV<br>VMNLWRDHWIYWVAPIVGGLLATCTY<br>QLFFARRRINQNTLEDDNYNVELNDND<br>TSKFMVSPL |

| | | | | |
|---|---|---|---|---|
| *Stylophora pistillata* | *aqp4L1.3* | XM_022934997.1<br>XP_022790732.1 | ATGACAGATTCCGCGATCCGTAAGCGCAAATTTTGGATTTGCGTCGGGGCAGAATTTCTGGCGACGACAATTTT<br>TGTTTTTGTGGTTTGCGGCTCGACGCTAACCGTGCACTCGAACGAGTCGGTCGGCGTGTTACATATCTCGCTTA<br>CTGCAGGTCTAACGGTGGCGACACTAGCGATGGCGACAGGACATCTAACTGGAGGCATGGTAAATCCTGCGCT<br>AACCATTGCGTTGATGGCAACTAAAAAGATTTCGGTGTTACAAGGAGTGTTTTACACAGTCGCACAATTTCTTG<br>GTGCGGTTTTAGGAGCAGCACTTCTATATGGTCTAACACCTAGCCAAATACGCGGGGCACTGGGAGCGACTACA<br>GTTGGTAGTGGCCTAAATGCTGGCCAAGCCTTCGGATTGGAGCTCTTTCTTACGTGTATATTGGTGTTCACCATA<br>TTTGCCGCCACTGATCCGGGGAAGGAGCTTCGGGGATATGACATACCGCTGTCCATAGGAGTGTGTGTGTTCAT<br>CTGTCACATGTGCGGGATTCCTTTCACTGGCTGCAGCATGAACCCTGCTAGGTCTTTCGGTCCCGCCCTGATTA<br>GTAACATCTGGAAAGATCACTGGGTTTACTGGGCCGGTCCCATACCGGGAGGAATAATAGCCGCATTCTTGTAT<br>GAATACGTCTTCTCCTCAAGCAAGACCGGAGTTTCTCCATCTTAA | MTDSAIRKRKFWICVGAEFLATTIFVFV<br>VCGSTLTVHSNESVGVLHISLTAGLTVAT<br>LAMATGHLTGGMVNPALTIALMATKKI<br>SVLQGVFYTVAQFLGAVLGAALLYGLT<br>PSQIRGALGATTVGSGLNAGQAFGLELF<br>LTCILVFTIFAATDPGKELRGYDIPLSIGV<br>CVFICHMCGIPFTGCSMNPARSFGPALIS<br>NIWKDHWVYWAGPIPGGIIAAFLYEYV<br>FSSSKTGVSPS |
| *Stylophora pistillata* | *glp1* | XM_022936741.1<br>XP_022792476.1 | ATGGCTAATAACATATCGAGAGGCCGATTGCCGCCATTGATGAGAGAAGCGCTGGCTGAGTTTATGGCAACATT<br>TATCTTAGTGTCATTTGGTCTTGGCTCCGTGGCCCAATTTACTCTTAGCCGCAGTCAATCTGGAACGTTCTTCTC<br>GGTAAATTTCAGCTGGGGAGTTGCAGTTACGATGGGTTGTTACTGGGCTGGGGGAATATCAGGTGCACATATGA<br>ATCCTGCTGTGACCCTGGCCCTTGCTGTGGTGCGGCGATTGAGGTGGACTAAAGTTCCTGCATACTGCCTCGCT<br>CAACTGCTTGGATCTTTTTTTGCGTCAACTATTGTTTATGGAATTTATCGCGATGCTCTTGACGCATTCGACGGTG<br>GTGTTCGTCAAGTGACAGGCCCCAACGCTACCGCAGGGGTCTTTGCAACGTATCCAAAACCTTTCCTGACTAC<br>TGCAAATGGATTCGCTGATCAGTTTTTTGGCACTGCTCTCCTTGTCAGCTGTATATTCGCTATCCTTGACCGTAAC<br>AACAATGCCCCCGACCAGGGGGTGGCACCGGTTATGATTGGTCTTGTCGTGTTTTTGATTGGCACCACCTTTGG<br>ATTTAACTGCGGTTACCCCATCAACCCTGCGAGGGATCTGGGACCAAGAATTTTCACCGCAATGGCCGGATGGG<br>GATCTGAAGTGTTTACGGCATATAATTATTGGGCGTGGGTCCCAATCATAGCCTGTTCACTAGGGGGAGTACTGG<br>GCGCCATTGTCTACATCACTACCATCGAGTTTCATCACTCAGCATCGGGAGAGACAGAATATCCTTATGAACCCA<br>TTGTTACAGCGGCCGTCACAACAATAAAGAAATGA | MANNISRGRLPPLMREALAEFMATFILV<br>SFGLGSVAQFTLSRSQSGTFFSVNFSWG<br>VAVTMGCYWAGGISGAHMNPAVTLAL<br>AVVRRLRWTKVPAYCLAQLLGSFFASTI<br>VYGIYRDALDAFDGGVRQVTGPNATAG<br>VFATYPKPFLTTANGFADQFFGTALLVS<br>CIFAILDRNNNAPDQGVAPVMIGLVVFL<br>IGTTFGFNCGYPINPARDLGPRIFTAMA<br>GWGSEVFTAYNYWAWVPIIACSLGGVL<br>GAIVYITTIEFHHSASGETEYPYEPIVTA<br>AVTTIKK |
| *Pocillopora damicornis* | *aqp4L1.2* | XM_027204471.1<br>XP_027060272.1<br>(RMX53469.1) | ATGTATCTCAGTGAAGGCAAAATGGAGTTTAGATCTCGCAAATTTTGGACCGAAGTCTTCGCGGAATTTTTGGC<br>CACACTTCTGTTCCTGTTCATGGTCTGTGGCACAGTGTTACCGTGGAATAACACTCCACCATCCGTTATACACAT<br>TGCTTTCAGTCATGGACTAAGTATAGCGACTTTAGCGATGGCGGTGTTTCACATTTCGGGCGGACAGCTCAACC<br>CAGCCGTGACCATCAGCATGATGGCTGTTGGAAGGGTCTCAGTCTTAAAAGCTGTGTTCTTCATTATTGCCCAG<br>TGTGCCGGAGCTATTTGTGGAGCAGCTATAGTTTACTACATGACCCCAACTGATTCTATTGGCTCATTGGGTGTC<br>ACAGCTCCTTCAAATGGTGTCTCTACTGCGCAGGCGTTTGGAGTGGAACTCATGCTGACTTTCATATTGGTTTTT<br>GTGATATTCGCTGCGACCGACCCCAACAGGGGTATGGCAGGATATGGGGTACCGCTGGCCATAGGGATTTGTGT<br>GTTCATTTGTCTCATGCATGGGATTCCTTCATCTGGAGCCAGCTTGAATCCAGCTCGTTCTCTTGGTCCTGCCGT<br>TGTAATGAATTTGTGGAGAGACCATTGGATTTACTGGGTGGCACCCACTCTCGGCGGTCTCCTGGCAACGTGCA<br>CCTACAAGCTGTTTTTCGCCCAGAGGAGAAACAACCAAAACACTTTGGAAGATGATAATTACAATGTGGAACT<br>CAACGACAACGATACATCGAAGTTTATGATCTCCCCTTTGTAA | MYLSEGKMEFRSRKFWTEVFAEFLATL<br>LFLFMVCGTVLPWNNTPPSVIHIAFSHG<br>LSIATLAMAVFHISGGQLNPAVTISMMA<br>VGRVSVLKAVFFIIAQCAGAICGAAIVY<br>YMTPTDSIGSLGVTAPSNGVSTAQAFGV<br>ELMLTFILVFVIFAATDPNRGMAGYGVP<br>LAIGICVFICLMHGIPSSGASLNPARSLG<br>PAVVMNLWRDHWIYWVAPTVGGLLAT<br>CTYKLFFAQRRNNRNTLEDDNYNVELN<br>DNDTSKFMISPL |
| *Pocillopora damicornis* | *aqp4L1.3* | XM_027204471.1<br>XP_027060272.1<br>(RMX53470.1) | ATGACAGACTTCGCGCTCCGTAAGCGCAAATTTTGGACATGCGTTGGGGCAGAGTTCTTAGCGACAACAATTTT<br>TGTTTTTGCGGTTAGCAGCTCGACGCTAACCGTGCAAGGGGACGCTACTGTCGTTGTGTTGCATGTCTCGCTTA<br>CTGCAGGTCTAACTGTGGCGACATTAGTGATGGCGACAGGACATCTAACGGGAGGCCTTATAAATCCCGCGCTA<br>ACAACTGCGCTAATGGCTACCAAAAAGATTTCGGTGTTAGAAGGAGTGTTTTACATAGTCGCGCAATTTCTTGG<br>AGCGATTTTAGGAGCAGCACTTCTATATGGTCTGACACCTGGCAAAATGCGCGGGGCACTGGGAGCGACCACA<br>ATCAGTAGCGACCTTAGTGCTAGCCAAGCCTTTGGACTGGAGCTCCTTCTCACGTTTGTGTTGGTGTTTACCATA<br>TTTGCCGCTACTGATCCGGGCAGGGAACTTCGAGGATACGACATACCGCTGTCCATAGGAGTGTGTGTGTTCAT<br>CTGTCACATGTGCGGGATTCCTCTCACTGGCTGCAGCATGAACCCCGCCAGGTCTCTCGGTCCTGCCCTAATTA<br>GAAACTTCTGGAAAGACCACTGGGTTTACTGGGTCGGTCCCGTACCAGGAGGAATTATAGCGGCATTCTTGTAT<br>GAATACGTCTTCTCCACAAGGAGTGCTGGAGTTTCTCCAGCTTAA | MTDFALRKRKFWTCVGAEFLATTIFVF<br>AVSSSTLTVQGDATVVVLHVSLTAGLTV<br>ATLVMATGHLTGGLINPALTTALMATKK<br>ISVLEGVFYIVAQFLGAILGAALLYGLTP<br>GKMRGALGATTISSDLSASQAFGLELLL<br>TFVLVFTIFAATDPGRELRGYDIPLSIGV<br>CVFICHMCGIPLTGCSMNPARSLGPALIR<br>NFWKDHWVYWVGPVPGGIIAAFLYEY<br>VFSTRSAGVSPA |
| *Exaiptasia diaphana* | *aqp4L1.1* | XM_021042014.1<br>XP_020897673.1 | ATGGTTTTACAAGAAGTCAAAACCAAAGAGTTCTGGATATGTATGTTGGCGGAATTGCTTGCTACCTTCTTTTTT<br>GTGTTCTTGACGACTGGTTCAACGATTACCTGGGATCCCAACCATCCACCAACGATTGAGCACATTTCCTTGTC<br>GTTTGGTCTCAGCATTGCAACACTGGTTATGTCTATTCTTCACATAAGTGGTGGTCTTATTAACCCTGCTGTCAG<br>CGTGGCTCAAATTGTTACCAGAAAAATATCAATCATTCGTGGAGTTGCTTTTATCGTCTTTCAATCGGCCGGAGG<br>AATTTTAGGATCGGCGTTATTATACGGGCTAACTCCAGCAACAAAGCGAGGGACCCTTGGTGTTACCGTTCCTG<br>GCGATGGTGTAGATGTTGGTCAAGCGTTTGGGATAGAGTTTGTGTTAACATTTCTACTAGTTCTGACCATATTTG<br>CTTGTACCGATCCTAAGAGGAATCACTATGGGTATGAAGTTCCTTTGGCCATTGGATTTTGTGTCGCTGTTTGCC<br>ATATGATTGGAATACGTTTTACTGGATGTGGTATCAATCCTGCTAGATCACTTGGGCCCGCTGTTGTTATGGAAAT<br>CTGGACAGACCACTGGGTATACTGGGTTGGTCCTATGGGTGGAGCACTCGTTGCTAGTTTATTCTATGTGACTGT<br>ATTCCGTGCTCGTACAGAACAAGGCCAATCTGAAGGCAATATGGATGCTGTCGTATTAAGCAATGTTACCATCA<br>ATAATAACAAGTAA | MVLQEVKTKEFWICMLAELLATFFFVF<br>LTTGSTITWDPNHPPTIEHISLSFGLSIAT<br>LVMSILHISGGLINPAVSVAQIVTRKISIIR<br>GVAFIVFQSAGGILGSALLYGLTPATKRG<br>TLGVTVPGDGVDVGQAFGIEFVLTFLLV<br>LTIFACTDPKRNHYGYEVPLAIGFCVAV<br>CHMIGIRFTGCGINPARSLGPAVVMEIW<br>TDHWVYWVGPMGGALVASLFYVTVFR<br>ARTEQGQSEGNMDAVVLSNVTINNNK |
| *Exaiptasia diaphana* | *aqp4L1.2a* | XM_021040589.2<br>XP_020896248.1 | ATGACCAACGACGTAAATTCTTGTTTCTTTCGGTTCAAAAATGAAGTCAAGAGCGGTAAACTCTGGGTTGAAG<br>TATTTGCAGAGTTTCTCATCACAACTCTGTTTATCTCGTTTGTGTGTGGATGTTCGTTGGCATGGAATCCTCCCTC<br>GTCTGTGATGCACATGGCTATAAGTTCAGGTTTAGGAGTTGCTACGTTTGCCTTTACCATGTGGGATGTGTCATC<br>TGGTCTTTTTAATCCTGCTCTTACCGTTGCTTTCTTTGTTGGTGGAAAGAAGACGCTTGCGCAAACGTTGTTTTA<br>CATCATAGCTCAAGTTATTGGAGGTATTTGCGGTGCTGCAGTGATTTATGGGATAGCTCCAGAACAAGCCATCGT<br>TGATTCAAATCTCGCTGTCAATCTTCGCAATCCTGATGTGTCTGTTGGCAAGGCGTTAGGAATGGAAATGTGGA<br>TTACTTTTATTCTTGTTCTAGTGGTGTTTGCTGCTGGTGACCCTGACAGACATCTGTCAGGATATGGACCACCGC<br>TGGCCATTGGATTCGTCGTATTTGTCAATCTGAACCTCTCGATTAACTTATCTGGAGGTGGAATGAATCCAACGC<br>GATCATTTGGGCCAGCGGTTGTCATGAATTCTTGGGAAGACCACTGGATCTACTGGGTTGGCCCGATGATAGGA<br>GCTATTGCGGCAGCGCCGATCTACAACTACGTGTTTTCGCAAAGGGTATACAACAGTCTGCTATCTCAACCACA<br>ACAGTCTGAGAACTCAACCTCAAACCACGATCTTGACAACGAAAATATACAATTTCAACCATGA | MTNDVNSCFFRFKNEVKSGKLWVEVF<br>AEFLITTLFISFVCGCSLAWNPPSSVMH<br>MAISSGLGVATFAFTMWDVSSGLFNPAL<br>TVAFFVGGKKTLAQTLFYIIAQVIGGIC<br>GAAVIYGIAPEQAIVDSNLAVNLRNPDV<br>SVGKALGMEMWITFILVLVVFAAGDPD<br>RHLSGYGPPLAIGFVVFVNLNLSINLSG<br>GGMNPTRSFGPAVVMNSWEDHWIYWV<br>GPMIGAIAAAPIYNYVFSQRVYNSLLSQ<br>PQQSENSTSNHDLDNENIQFQP |
| *Exaiptasia diaphana* | *aqp4L1.2b* | XM_021040712.2<br>XP_020896371.1 | ATGACTAAGAACATAAGTTCTTGTTTCGTTCGGTTTAAAAATGAAGTCAAAAGCGGTAAACTCTGGGTTGAAGT<br>ATTTGCAGAGTTTCTCATCACAACTCTGTTTATCTCATTTGTGTGTGGATGCTCGTTAGCATGGAATCCTCCCTCG<br>TCTGTGATGCACATGGCTATAAGTTCAGGTTTAGGAGTTGCTACGTTTGCCTTTACCATGTGGGATGTGTCATCT<br>GGTCTTTTTAATCCTGCTCTTACCGTTGCTTTCTTTGTTGGTGGAAAGAAGACGCTTGCGCAAACGTTGTTTTAC<br>ATCATAGCTCAAGTTATTGGAGGTATTTGCGGTGCTGCAGTGATTTATGGGATAGCTCCAGAGCAAGCCATAATT<br>GATTCAAATCTGGCTGTCAATCTTCGCAATCCTGATGTGTCTGTTAGCAAGGCGTTAGGAATGGAAATATGGATT<br>ACTTTTATTCTTGTTCTAGTGGTGTTTTCGGCTGGTGACCCTGACAGACATCTGTCAGGATATGGACCACCGCTG<br>GCCATTGGATTCGTCGTATTTGCCAATCTGAACCTCTCGATCAATTTATCTGGAGGTGGAATGAATCCAACGCGA<br>TCATTCGGGCCAGCGGTTGTCATGAATTCTTGGCAGGACCACTGGATCTACTGGGTTGGCCCTATGATAGGAGC<br>TATTGCGGCAGCGCTGATCTATAACTACGTGTTTGCGCAAAGAGTTTACACAAAAATTGCCAAAACATCCTACG<br>ACCTCAACCATGACGACGTTAAGGCAAAGAAGAGCGGATCTGTACATTCCCAGGAAAAGATAAATATGGATATA<br>CAGTAG | MTKNISSCFVRFKNEVKSGKLWVEVFA<br>EFLITTLFISFVCGCSLAWNPPSSVMHM<br>AISSGLGVATFAFTMWDVSSGLFNPALT<br>VAFFVGGKKTLAQTLFYIIAQVIGGICG<br>AAVIYGIAPEQAIIDSNLAVNLRNPDVSV<br>SKALGMEIWITFILVLVVFSAGDPDRHL<br>SGYGPPLAIGFVVFANLNLSINLSGGGM<br>NPTRSFGPAVVMNSWQDHWIYWVGPM<br>IGAIAAALIYNYVFAQRVYTKIAKTSYD<br>LNHDDVKAKKSGSVHSQEKINMDIQ |

#### Table S2. Aquaporin sequences with accession numbers and annotation notes.

| **Species** | Sequence name used in tree | Accezsion no. (original annotation) | Note | Amino acid sequence |
|---|---|---|---|---|
| *Corticium candelabrum* | XP_062508334.1_N | XP_062508334.1 | N-terminal fragment | MEEEDAIVHRPPSPTRRHETSIKWWAMLIRPCLAEYIGTTLFVFAGILSTFHSANEKGAVVN<br>VALGHGMALFVMVATTATVSGGHLNPAVTLGIFLTGTLAPVVALGFVGAQLLGGITGAGLA<br>KGLLNSTVTKSESGFALTEPPSYQDDVQAIFAEVLLTAILVLTVLMVAVETKGNNPIAPFAIG<br>MAVTAGIFAGGNVSGASMNPARSFGPAVVFNTWQRVWVYFVGCPVGGIIASIAYRLVLTDR<br>SLPIPIVSFFKRHKVGSC |
| *Corticium candelabrum* | XP_062508334.1_C | XP_062508334.1 | C-terminal fragment | MGESEPVSDVQNSQVGAVQSRSDKVVQTLPQEQSESYENSKYWTKVARPLLAEFIGTMFF<br>HCAGIITSVEANPVVGPLGISMTLLGLCGVFFPLSGAQFNPAVSLAAFLSGLMAFDMMILYIL<br>AQLAGGTASAAIAYGLINSTAVLRNPDGWATNAYNAAYQNEAQAFFAEILLTCILVTVILSSM<br>ADCDGKNQFTWIFMIFTVAGNTFTGAYISGAGMNPSRSFGPALVFNYWGQLWIYTFGPLIG<br>ACIAVVIYRFMTHHRKLRMPYLTFRDEEGWDDD |
| *Xenia* sp. Carnegie-2017 | XP_046851093.1_N | XP_062508334.1 | N-terminal fragment | MEDHRFTIEEQPPRSQFESSLIFKYIRPCIAEMFGTILIVYTDICLGWDNTFAHGYLMFALIAA<br>TSSVSGGCFNPVVTLALTIFRGIQPVLALLFIISQILGGIIGAALAQVAMSSADYKLNFGGVRT<br>LGGAHALYPYRTPKTEISPGQGILAEALATSILVLVVLMVLLDERNKSKLAPLAIGLSVVGGI<br>AANRWITGGSMNPVRSFGAAVVKNYWKHHYVYWVGPFLGGMLSTLMYGFVLASPNQLW<br>LVW |
| *Xenia* sp. Carnegie-2017 | XP_046851093.1_C | XP_062508334.1 | C-terminal fragment | MEEDNRFNIEEQPSKLRVYSTLFFKFIRPCVAELFGTTIFVFADLGGMNYISSTAFTHGLILFV<br>LIAGTATVSGGHFNPAVTLALTIVRGLQPVLAPLYILSQLLGGIIGAGLARAAMSSEQYTKTF<br>GGVHMLGADINPGQGILAEALATSLLVFVVLMVAFDERNKSEFAPLAIGFTVTGGILAIGVIS<br>GGSMNPARSFGPAVVKNYWKHHYVYWVGPFLGSILSTLMYGFVFATPSQLWLPLREPIT |

Table S3. Accession numbers for the *aqp* genes and their encoded proteins in vertebrate.

| | *Homo sapiens* | *Mus musculus* | *Xenopus tropicalis* | *Takifugu rubripes* | *Danio rerio* |
|---|---|---|---|---|---|
| *aqp0* | NM_198098.4<br>NP_036196.1 | NM_008600.5<br>NP_032626.2 | NM_001097347.1<br>NP_001090816.1 | (*0a*) XM_003963091.2<br>XP_003963140.1<br>(*0b*) XM_003963426.2<br>XP_003963475.1 | (*0a*) NM_001003534.1<br>NP_001003534.1<br>(*0b*) NM_001020520.1<br>NP_001018356.1 |
| *aqp1* | NM_198098.4<br>NP_932766.1 | NM_007472.2<br>NP_031498.1 | NM_001005829.1<br>NP_001005829.1 | (1*aa*) LC735283.1<br>BDT49272.1<br>(1*ab*) LC735284.1<br>BDT49273.1 | (1*aa*) NM_207059.1<br>NP_996942.1<br>(1*ab*) NM_001135682.1<br>NP_001129154.1 |
| *aqp2* | NM_000486.6<br>NP_000477.1 | NM_009699.3<br>NP_033829.3 | | | |
| *aqp3* | NM_004925.5<br>NP_004916.1 | NM_016689.2<br>NP_057898.2 | NM_001016845.2<br>NP_001016845.1 | AB610903.1<br>BBE28311.1 | (*3a*) NM_213468.1<br>NP_998633.1<br>(*3b*) NM_001166121.2<br>NP_001159593.1 |
| *aqp4* | NM_001317384.3<br>NP_001304313.1 | NM_009700.3<br>NP_033830.2 | (*4.1*) NM_001317845.1<br>NP_001304774.1<br>(*4.2*) XM_018094938.2<br>XP_017950427.1<br>(*4.3*) NM_001142111.1<br>NP_001135583.1 | LC735286.1<br>BDT49275.1 | NM_001003749.6<br>NP_001003749.1 |
| *aqp5* | NM_001651.4<br>NP_001642.1 | NM_009701.4<br>NP_033831.1 | (5L) NM_001310112.1<br>NP_001297041.1<br>(5) NM_001015749.1<br>NP_001015749.1 | | |
| *aqp6* | NM_001652.4<br>NP_001643.2 | NM_175087.5<br>NP_780296.1 | (6ub) XM_002935732.3<br>XP_002935778.1<br>(6vs) XM_012958316.3<br>XP_012813770.2 | | |
| *aqp7* | NM_001376192.1<br>NP_001363121.1 | NM_001378639.1<br>NP_001365568.1 | NM_001015726.1<br>NP_001015726.1 | LC735287.1<br>BDT49276.1 | NM_199910.2<br>NP_956204.2 |
| *aqp8* | NM_001169.3<br>NP_001160.2 | NM_007474.2<br>NP_031500.1 | NM_001114256.1<br>NP_001107728.1 | LC735288.1<br>BDT49277.1 | (*8aa*) NM_001004661.1<br>NP_001004661.1<br>(*8ab*) NM_001080182.1<br>NP_001073651.1<br>(*8bb*) NM_001114910.3<br>NP_001108382.2 |
| *aqp9* | NM_020980.5<br>NP_066190.2 | NM_022026.3<br>NP_071309.1 | XM_002937673.5<br>XP_002937719.2 | (*9a*) XM_003967709.3<br>XP_003967758.1<br>(*9b*) LC735290.1<br>BDT49279.1 | (*9a*) NM_001033096.1<br>NP_001028268.1<br>(*9b*) NM_001177744.1<br>NP_001171215.1 |
| *aqp10* | NM_080429.3<br>NP_536354.2 | | XM_018090358.2<br>XP_002943404.2 | (*10aa*) LC735291.1<br>BDT49280.1<br>(*10bb*) LC735292.1<br>BDT49281.1 | (*10a*) NM_001002349.2<br>NP_001002349.1<br>(*10b*) NM_001083047.1<br>NP_001076516.1 |
| *aqp11* | NM_173039.3<br>NP_766627.1 | NM_175105.3<br>NP_780314.1 | NM_001445251.1<br>NP_001432180.1 | (*11a*) XM_003968221.3<br>XP_003968270.2<br>(*11b*) LC735294.1<br>BDT49283.1 | NM_001327893.1<br>NP_001314822.1 |
| *aqp12* | NM_198998.3<br>NP_945349.1 | NM_177587.2<br>NP_808255.1 | XM_031902958.1<br>XP_031758818.1 | LC735295.1<br>BDT49284.1 | NM_001045862.2<br>NP_001039327.1 |
| *aqp13* | | | XM_002940465.5<br>XP_002940511.2 | | |
| *aqp14* | | | XM_031896224.1<br>XP_031752084.1 | LC735296.1<br>BDT49285.1 | XM_005174125.6<br>XP_005174182.1 |
| *aqp16* | | | XM_002937020.5<br>XP_002937066.2 | | |

### Table S4. Summary of genes and accession numbers used for *aqp4L1.1, 1.2, 1.3* synteny analysis.

| **#** | **Species** | **Chromosome (direction)** | **Upstream** | **Upstream** | **Upstream** | **Upstream** | **Upstream** | **Upstream** | **Upstream** | **Upstream** | **Upstream** | **Upstream** | **Upstream** | **Upstream** | **Upstream** | **Target gene** | **Target gene** | **Target gene** | **Target gene** |
|---|---|---|---|---|---|---|---|---|---|---|---|---|---|---|---|---|---|---|---|
| 1 | *Acropora millepora* | Chromosome 3 (Minus) | <*id2l*<br>XM_029346685.2<br>XP_029202518.2 | <LOC114949846<br>XM_044317998.1<br>XP_044173933.1 | <LOC122957674<br>XM_044317997.1<br>XP_044173932.1 | <*mcphl*<br>XM_029346259.2<br>XP_029202092.2 | >*glua1l*<br>XM_029345588.2<br>XP_029201421.2 | | <*glua2l*<br>XM_029345556.2<br>XP_029201389.2 | | <*prkar2a*<br>XM_044317996.1<br>XP_044173931.1 | | | | < LOC114965506<br>XM_029345207.2<br>XP_029201040.2 | **>*aqp4L1.1*<br>XM_029346894.2<br>XP_029202727.1** | **>*aqp4L1.2***<br>**XM_044318315.1<br>XP_044174250.1** | | **>*aqp4L1.3***<br>**XM_044318315.1<br>XP_044174250.1** |
| 2 | *Acropora digitifera* | NW_015441625.1 (Plus) | <*id2l*<br>XM_015920914.1<br>XP_015776400.1 | | <*mcphl*<br>XM_015920912.1<br>XP_015776398.1 | <*mcphl*<br>XM_015920897.1<br>XP_015776383.1 | >*glua1l*<br>XM_015920899.1<br>XP_015776385.1 | >*glua3l*<br>XM_015920911.1<br>XP_015776397.1 | <*glua2l*<br>XM_015920910.1<br>XP_015776396.1 | | <*prkar2a*<br>pseudo | | | >ncRNA<br>XR_001566821.1 | <LOC107354422<br>XM_015920904.1<br>XP_015776390.1 | **>*aqp4L1.1*<br>XM_015920905.1<br>XP_015776391.1** | **>*aqp4L1.2***<br>**XM_015920900.1<br>XP_015776386.1** | | **>*aqp4L1.3***<br>**XM_015920900.1<br>XP_015776386.1** |
| 3 | *Porites lutea* | Chromosome 11 (Minus) | >LOC140951923<br>XM_073401299.1<br>XP_073257400.1 | >LOC140951794<br>pseudo | <LOC140952293<br>XM_073401720.1<br>XP_073257821.1 | >LOC140951793<br>XM_073401142.1<br>XP_073257243.1 | >*mtnr1al*<br>XM_073401948.1<br>XP_073258049.1 | <*glua2l*<br>XM_073402811.1<br>XP_073258912.1 | <*glua2l*<br>XM_073401467.1<br>XP_073257568.1 | <LOC140952057<br>XM_073401469.1<br>XP_073257570.1 | <*prkar2a*<br>pseudo | | | | <LOC140953317<br>XM_073402810.1<br>XP_073258911.1 | **>*aqp4L1.1*<br>XM_073400850.1<br>XP_073256951.1** | **>*aqp4L1.2*<br>XM_073400849.1<br>XP_073256950.1** | | **>*aqp4L1.3*<br>XM_073400851.1<br>XP_073256952.1** |
| 4 | *Stylophora pistillata* | NW_019217930.1 (Minus) | <*thsd7a*<br>XM_022934943.1<br>XP_022790678.1 | >LOC111330137<br>pseudo | <*glua3l*<br>XM_022934982.1<br>XP_022790717.1 | <LOC111330177<br>XM_022935001.1<br>XP_022790736.1 | >*melanopsin-a-like*<br>XM_022935007.1<br>XP_022790742.1 | <*glua2l*<br>XM_022935000.1<br>XP_022790735.1 | <*glua2l*<br>XM_022934999.1<br>XP_022790734.1 | | <*prkar2a*<br>pseudo | | | | <LOC111330174<br>XM_022934998.1<br>XP_022790733.1 | **>*aqp4L1.1*<br>XM_022935003.1<br>XP_022790738.1** | **>*aqp4L1.2***<br>**XM_022934997.1<br>XP_022790732.1** | | **>*aqp4L1.3***<br>**XM_022934997.1<br>XP_022790732.1** |
| 5 | *Pocillopora damicornis* | NW_020843848.1 (Minus) | <*thsd7a*<br>XM_027204507.1<br>XP_027060308.1 | >ncRNA<br>XR_003447684.1 | <*glua3l*<br>XM_027204506.1<br>XP_027060307.1 | <*microcephalin*<br>XM_027204492.1<br>XP_027060293.1 | | <*glua2l*<br>XM_027204489.1<br>XP_027060290.1 | <LOC113686803<br>XM_027204474.1<br>XP_027060275.1 | <*glua2l*<br>XM_027204472.1<br>XP_027060273.1 | <*prkar2a*<br>pseudo | | | <ncRNA<br>XR_003447683.1 | <LOC113686824<br>XM_027204501.1<br>XP_027060302.1 | **>*aqp4L1.1*<br>XM_027204487.1<br>XP_027060288.1** | **>*aqp4L1.2***<br>**XM_027204471.1<br>XP_027060272.1** | | **>*aqp4L1.3***<br>**XM_027204471.1<br>XP_027060272.1** |
| 6 | *Orbicella faveolata* | NW_018150331.1 (Minus) | <*col2a1*<br>XM_020776793.1<br>XP_020632452.1 | <ncRNA<br>XR_002298869.1 | <*glua2l*<br>XM_020776732.1<br>XP_020632391.1 | <*glua4l*<br>XM_020776808.1<br>XP_020632467.1 | <*glua4l*<br>XM_020776735.1<br>XP_020632394.1 | <*glua3l*<br>XM_020776734.1 | <*glua2l*<br>XM_020776792.1 | | <*prkar2a*<br>pseudo | >*samd3*<br>XM_020776736.1<br>XP_020632395.1 | >LOC110069249<br>XM_020776743.1<br>XP_020632402.1 | <ncRNA<br>XR_002298870.1 | <LOC110069298<br>XM_020776803.1<br>XP_020632462.1 | **>*aqp4L1.1*<br>XM_020776758.1<br>XP_020632417.1** | **>*aqp4L1.2*<br>XM_020776756.1<br>XP_020632415.1** | | **>*aqp4L1.3*<br>XM_020776759.1<br>XP_020632419.1** |
| 7 | *Nematostella vectensis* | Chromosome.4 (Plus) | | >*afar*<br>XM_032382632.2<br>XP_032238523.2 | <*enoph1*<br>XM_032382628.2<br>XP_032238519.2 | >*vgat*<br>XM_032382630.2<br>XP_032238521.1 | <*glua3l*<br>XM_001633306.3<br>XP_001633356.3 | <*glua2l*<br>XM_048726930.1<br>XP_048582887.1 | <*glua2l*<br>XM_048726674.1<br>XP_048582631.1 | <*vgat*<br>XM_048726675.1<br>XP_048582632.1 | <*prkar2a*<br>XM_001633176.3<br>XP_001633226.2 | | | | >LOC116618530<br>XM_032382335.2<br>XP_032238226.2 | **>*aqp4L1.1*<br>XM_032382406.2<br>XP_032238297.1** | **<*aqp4L1.2*<br>XM_001633177.3<br>XP_001633227.2** | | **>*aqp4L1.3*<br>XM_032382425.2<br>XP_032238316.2** |
| 8 | *Exaiptasia diaphana* | NW_018385184.1 (Minus) | <*odc*<br>XM_021041954.2<br>XP_020897613.1 | <*klf10*<br>XM_021041969.2<br>XP_020897628.1 | | | <*qrfpr*<br>XM_021041958.2<br>XP_020897617.1 | >*slc2a12*<br>XM_028658294.1<br>XP_028514095.1 | <LOC110236435<br>XM_021041956.2<br>XP_020897615.1 | <*ectin*<br>XM_021041960.2<br>XP_020897619.1 | <*zfp809*<br>XM_021041961.2<br>XP_020897620.1 | >*abcd4*<br>XM_021041999.1<br>XP_020897658.1 | | | <LOC110236477<br>XM_021042008.1<br>XP_020897667.1 | **>*aqp4L1.1***<br>**XM_021042014.1<br>XP_020897673.1** | | | **>*aqp4L1.3*<br>XM_021042000.2<br>XP_020897659.1** |
| 9 | *Exaiptasia diaphana* | NW_018384893.1 (Minus) | | | | | | | | | | | | | | | **<*aqp4L1.2c*<br>XM_021039591.2<br>XP_020895250.1** | | |
| 10 | *Exaiptasia diaphana* | NW_018384158.1 (Minus) | | >LOC110234660<br>XM_021040051.2<br>XP_020895710.1 | >LOC110234855<br>XM_021040262.2<br>XP_020895921.2 | <*qrfpr*<br>XM_021040646.2<br>XP_020896305.1 | <*qrfpr*<br>XM_021040700.2<br>XP_020896359.1 | | | | | | <*polr1a*<br>XM_028657905.1<br>XP_028513706.1 | >*mutsl*<br>XM_028657904.1<br>XP_028513705.1 | >*mutsl*<br>XM_028657961.1<br>XP_028513762.1 | | **<*aqp4L1.2a***<br>**XM_021040589.2<br>XP_020896248.1** | **<*aqp4L1.2b***<br>**XM_021040712.2<br>XP_020896371.1** | |
| 11 | *Actinia tenebrosa* | NW_022259781.1 (Minus) | | | | | <LOC116294201<br>XM_031701748.1<br>XP_031557608.1 | <LOC116294200<br>XM_031701747.1<br>XP_031557607.1 | <LOC116294219<br>XM_031701773.1<br>XP_031557633.1 | <LOC116294220<br>XM_031701774.1<br>XP_031557634.1 | < LOC116294204<br>XM_031701754.1<br>XP_031557614.1 | >LOC116294203<br>XM_031701753.1<br>XP_031557613.1 | <*ggsl*<br>XM_031701757.1<br>XP_031557617.1 | >*adamts7l*<br>XM_031701751.1<br>XP_031557611.1 | <*zdhhc4*<br>XM_031701755.1<br>XP_031557615.1 | **>*aqp4L1.1a*<br>XM_031701769.1<br>XP_031557629.1** | | | |
| 12 | *Actinia tenebrosa* | NW_022258470.1 (Minus) | | <*klf10*<br>XM_031694596.1<br>XP_031550456.1 | <*tkr86c*<br>XM_031694560.1<br>XP_031550420.1 | >*slc2a10*<br>XM_031694555.1<br>XP_031550415.1 | <*gapr1*<br>XM_031694566.1<br>XP_031550426.1 | <LOC116287856<br>XM_031694549.1<br>XP_031550409.1 | <*gapr1*<br>XM_031694573.1<br>XP_031550433.1 | <*ectin*<br>XM_031694550.1<br>XP_031550410.1 | <*contactin-4-like*<br>XM_031694590.1<br>XP_031550450.1 | >*abcd4*<br>XM_031694587.1<br>XP_031550447.1 | <*contactin-4-like*<br>XM_031694590.1<br>XP_031550450.1 | <*hemicentin-2-like*<br>XM_031694589.1<br>XP_031550449.1 | <LOC116287866<br>XM_031694575.1<br>XP_031550435.1 | **>*aqp4L1.1b*<br>XM_031694576.1<br>XP_031550436.1** | <ncRNA<br>XR_004196121.1 | >*aqp*<br>XM_031694577.1<br>pseudo | |
| 13 | *Actinia tenebrosa* | NW_022257979.1 (Plus) | | | | | | | | | | | | | >ncRNA<br>XR_004196316.1 | | | | **>*aqp4L1.3*<br>XM_031696179.1<br>XP_031552039.1** |

Direction: The arrows (>) indicate the transcriptional orientation of each gene on the genomic scaffold.

Gene Symbols: Official symbols are used where available; otherwise, NCBI LOC identifiers are provided.

Asterisks (*): Indicate full-length sequences determined in this study (refer to Supplementary Table S1 for sequence data).

(Continued from the previous page)

| # | Species | Chromosome (direction) | Downstream | Downstream | Downstream | Downstream | Downstream | Downstream | Downstream | Downstream | Downstream |
|---|---|---|---|---|---|---|---|---|---|---|---|
| 1 | *Acropora millepora* | Chromosome 3 (Minus) | | <*haus7*<br>XM_044317994.1<br>XP_044173929.1 | >*washc5l*<br>XM_044317993.1<br>XP_044173928.1 | <*a3arl*<br>XM_029346877.2<br>XP_029202710.2 | >*glua4l*<br>XM_029346863.2<br>XP_029202696.2 | >ncRNA<br>XR_003826504.2 | >LOC114966929<br>XM_029346916.2<br>XP_029202749.2 | >LOC122957672<br>XM_044317992.1<br>XP_044173927.1 | |
| 2 | *Acropora digitifera* | NW_015441625.1 (Plus) | | | >*washc5l*<br>XM_015920901.1<br>XP_015776387.1 | <*a3arl*<br>XM_015920918.1<br>XP_015776404.1 | >*glua4l*<br>XM_015920902.1<br>XP_015776388.1 | <*malrd1*<br>XM_015920903.1<br>XP_015776389.1 | <*nfyb*<br>XM_015920917.1<br>XP_015776403.1 | | |
| 3 | *Porites lutea* | Chromosome 11 (Minus) | | <*haus7*<br>XM_073401026.1<br>XP_073257127.1 | >*washc5l*<br>XM_073401025.1<br>XP_073257126.1 | <*slc16a14*<br>XM_073402010.1<br>XP_073258111.1 | <*slc16a14*<br>XM_073402809.1<br>XP_073258910.1 | <*slc16a12*<br>pseudo | <*slc16a12*<br>XM_073402808.1<br>XP_073258909.1 | <*ctl*<br>XM_073402807.1<br>XP_073258908.1 | >*mrpl15l*<br>XM_073401353.1<br>XP_073257454.1 |
| 4 | *Stylophora pistillata* | NW_019217930.1 (Minus) | | <*haus7*<br>XM_022935005.1<br>XP_022790740.1 | >*washc5l*<br>XM_022934965.1<br>XP_022790700.1 | <*slc16a10*<br>XM_022934966.1<br>XP_022790701.1 | >LOC111330191<br>XM_022935015.1<br>XP_022790750.1 | <*chfrl*<br>XM_022935013.1<br>XP_022790748.1 | | | >*mrpl15l*<br>XM_022934981.1<br>XP_022790716.1 |
| 5 | *Pocillopora damicornis* | NW_020843848.1 (Minus) | | <*haus7*<br>XM_027204494.1<br>XP_027060295.1 | >*washc5l*<br>XM_027204481.1<br>XP_027060282.1 | | | | | | |
| 6 | *Orbicella faveolata* | NW_018150331.1 (Minus) | <*mfsd6l*<br>XM_020776751.1<br>XP_020632410.1 | <*haus7*<br>XM_020776752.1<br>XP_020632411.1 | >*washc5l*<br>XM_020776791.1<br>XP_020632450.1 | <*slc16a8*<br>XM_020776801.1<br>XP_020632460.1 | | | | | >*mrpl15l*<br>XM_020776790.1<br>XP_020632449.1 |
| 7 | *Nematostella vectensis* | Chromosome.4 (Plus) | | <*haus7*<br>XM_001633203.3<br>XP_001633253.2 | >*washc5l*<br>XM_048726307.1<br>XP_048582264.1 | >*lactb*<br>XM_048726256.1<br>XP_048582213.1 | >ccdc63<br>XM_048726249.1<br>XP_048582206.1 | | | | |
| 8 | *Exaiptasia diaphana* | NW_018385184.1 (Minus) | >*taar13c-like*<br>XM_021042013.2<br>XP_020897672.1 | <*haus7*<br>XM_021042022.1<br>XP_020897681.1 | >*washc5l*<br>XM_021041971.2<br>XP_020897630.1 | <*mamdc2*<br>XM_021041974.2<br>XP_020897633.1 | <LOC110236451<br>XM_021041973.2<br>XP_020897632.2 | <*mucin-like*<br>XM_028658297.1<br>XP_028514098.1 | | | |
| 9 | *Exaiptasia diaphana* | NW_018384893.1 (Minus) | | <*lrp4*<br>XM_028657621.1<br>XP_028513422.1 | | | | | | | |
| 10 | *Exaiptasia diaphana* | NW_018384158.1 (Minus) | >nxRNA<br>XR_003695228.1 | <*lrp4*<br>XM_021040230.2<br>XP_020895889.2 | <*otop2*<br>XM_021040730.2<br>XP_020896389.1 | <LOC110235058<br>XM_021040488.2<br>XP_020896147.1 | <*cyp17a1*<br>XM_021040960.2<br>XP_020896619.1 | >*smarca4*<br>XM_028658044.1<br>XP_028513845.1 | <*trim71*<br>XM_021040942.2<br>XP_020896601.2 | <*trim71-like*<br>XM_028657765.1<br>XP_028513566.1 | |
| 11 | *Actinia tenebrosa* | NW_022259781.1 (Minus) | | <LOC116294212<br>XM_031701764.1<br>XP_031557624.1 | <*ciao2al*<br>XM_031701768.1<br>XP_031557627.1 | <*gcnt1*<br>pseudo | <LOC116294213<br>XM_031701766.1<br>XP_031557626.1 | <*tex9l*<br>XM_031701771.1<br>XP_031557631.1 | <*fem1bl*<br>XM_031701772.1<br>XP_031557632.1 | <*eef2*<br>XM_031701745.1<br>XP_031557605.1 | <*frmd5l*<br>XM_031701759.1<br>XP_031557619.1 |
| 12 | *Actinia tenebrosa* | NW_022258470.1 (Minus) | | | | | | | | | |
| 13 | *Actinia tenebrosa* | NW_022257979.1 (Plus) | | < LOC116289272<br>XM_031696170.1<br>XP_031552030.1 | >*washc5l*<br>XM_031696203.1<br>XP_031552063.1 | >*entpd1*<br>XM_031696141.1<br>XP_031552001.1 | >*ttc28l*<br>XM_031696153.1<br>XP_031552013.1 | >LOC116289300<br>XM_031696193.1<br>XP_031552053.1 | <LOC116289272<br>XM_031696170.1<br>XP_031552030.1 | | |

Direction: The arrows (>) indicate the transcriptional orientation of each gene on the genomic scaffold.

Gene Symbols: Official symbols are used where available; otherwise, NCBI LOC identifiers are provided.

Asterisks (*): Indicate full-length sequences determined in this study (refer to Supplementary Table S1 for sequence data).

### Table S5. Summary of genes and accession numbers used for *aqp4L2* synteny analysis.

| **#** | **Species** | **Chromosome (direction)** | **Upstream** | **Upstream** | **Upstream** | **Upstream** | **Upstream** | **Upstream** | **Upstream** | **Target gene** | **Target gene** | **Downstream** | **Downstream** | **Downstream** | **Downstream** | **Downstream** | **Downstream** | **Downstream** | **Downstream** |
|---|---|---|---|---|---|---|---|---|---|---|---|---|---|---|---|---|---|---|---|
| 1 | *Acropora millepora* | NW_025323129.1<br>(Minus) | | *>lrp4l*<br>XM_044315524.1<br>XP_044171459.1 | *<nkain*<br>XM_029351031.2<br>XP_029206864.1 | | *>snrnp48*<br>XM_044315530.1<br>XP_044171465.1 | *<znt2l*<br>XM_044315531.1<br>XP_044171466.1 | *<rab-30-like*<br>XM_044315533.1<br>XP_044171468.1 | | ***>aqp4L2***<br>**XM_029351016.2**<br>**XP_029206849.2** | <ncRNA<br>XR_003827258.2 | *>naprtl*<br>XM_029351009.2<br>XP_029206842.2 | | *>hmgcl*<br>XM_029351011.2<br>XP_029206844.2 | < LOC114970552<br>XM_029351018.2<br>XP_029206851.2 | *>hmgcl*<br>pseudo | <LOC114970552<br>XM_029351018.2<br>XP_029206851.2 | *<stam2l*<br>XM_029351020.2<br>XP_029206853.2 |
| 2 | *Acropora digitifera* | NW_015441086.1<br>(Minus) | | | *<nkain*<br>XM_015896138.1<br>XP_015751624.1 | >LOC107331475<br>XM_015896064.1<br>XP_015751550.1 | *>snrnp48*<br>XM_015896124.1<br>XP_015751610.1 | *<znt2l*<br>XM_015896063.1<br>XP_015751549.1 | *<rab-30-like*<br>pseudo | ***>aqp***<br>**XM_015896062.1**<br>pseudo | ***>aqp4L2****<br>**XM_015896123.1**<br>**XP_015751609.1** | | *>naprtl*<br>XM_015896122.1<br>XP_015751608.1 | | *>hmgcl*<br>XM_015896119.1<br>XP_015751605.1 | < LOC107331522<br>XM_015896120.1<br>XP_015751606.1 | | | *<stam2l*<br>XM_015896061.1<br>XP_015751547.1 |
| 3 | *Porites lutea* | Chr.8<br>(Plus) | | | *<nkain*<br>XM_073395364.1<br>XP_073251465.1 | >LOC140946260<br>XM_073395353.1<br>XP_073251454.1 | *>snrnp48*<br>XM_073395353.1<br>XP_073251454.1 | *<znt2l*<br>XM_073395359.1<br>XP_073251460.1 | *<kiaa1958l*<br>XM_073395354.1<br>XP_073251455.1 | | ***>aqp4L2***<br>**XM_073395360.1**<br>**XP_073251461.1** | <ncRNA<br>XR_012166829.1 | *>naprtl*<br>XM_073395351.1<br>XP_073251452.1 | | *>hmgcl*<br>XM_073395357.1<br>XP_073251458.1 | <LOC140946272<br>XM_073395362.1<br>XP_073251463.1 | | | *<stam2l*<br>XM_073395349.1<br>XP_073251450.1 |
| 4 | *Stylophora pistillata* | NW_019217903.1<br>(Plus) | *>cdh23l*<br>XM_022933095.1<br>XP_022788830.1 | | *<nkain*<br>XM_022933145.1<br>XP_022788880.1 | | *>snrnp48*<br>XM_022933147.1<br>XP_022788882.1 | *<znt2l*<br>XM_022933149.1<br>XP_022788884.1 | | | ***<aqp4L2***<br>**XM_022933122.1**<br>**XP_022788857.1** | <ncRNA<br>XR_002694633.1 | *>naprtl*<br>XM_022933146.1<br>XP_022788881.1 | >LOC111328660<br>XM_022933138.1<br>XP_022788873.1 | *>hmgcl*<br>XM_022933123.1<br>XP_022788858.1 | <LOC111328657<br>XM_022933133.1<br>XP_022788868.1 | | | *<stam2l*<br>XM_022933108.1 |
| 5 | *Pocillopora damicornis* | NW_020845701.1<br>(Plus) | *>cdh23l*<br>XM_027190837.1<br>XP_027046638.1 | *>a2brl*<br>XM_027190856.1<br>XP_027046657.1 | *<nkain*<br>XM_027190824.1<br>XP_027046625.1 | | *>snrnp48*<br>XM_027190822.1<br>XP_027046623.1 | *<znt2l*<br>XM_027190823.1<br>XP_027046624.1 | | | ***<aqp4L2***<br>**XM_027190821.1**<br>**XP_027046622.1** | <ncRNA<br>XR_003446511.1 | *>naprtl*<br>XM_027190819.1<br>XP_027046620.1 | | *<hmgcl*<br>XM_027190820.1<br>XP_027046621.1 | <LOC113674351<br>XM_027190828.1<br>XP_027046629.1 | | | *<stam2l*<br>XM_027190829.1<br>XP_027046630.1 |
| 6 | *Orbicella faveolata* | NW_018150003.1<br>(Minus) | | *>octn4l*<br>XM_020769731.1<br>XP_020625390.1 | *> octn4l*<br>XM_020769730.1<br>XP_020625389.1 | *>slc22a5*<br>XM_020769723.1<br>XP_020625382.1 | *>snrnp48*<br>XM_020769722.1<br>XP_020625381.1 | *<znt2l*<br>XM_020769721.1<br>XP_020625380.1 | | | ***<aqp4L2***<br>**XM_020769737.1**<br>**XP_020625396.1** | | *>naprtl*<br>XM_020769734.1<br>XP_020625393.1 | | | >ncRNA<br>XR_002298151.1 | *>mshrl*<br>XM_020769738.1<br>XP_020625397.1 | >ncRNA<br>XR_002298155.1 | *<cdh23l*<br>XM_020769720.1<br>XP_020625379.1 |
| 7 | *Nematostella vectensis* | Chr. 11<br>(Minus) | | >ncRNA<br>XR_007314165.1 | <acyl-lipid (8-3)-desaturase<br>XM_048734376.1<br>XP_048590333.1 | *>acsf2*<br>XM_032368610.2<br>XP_032224501.2 | *>pask*<br>XM_048734457.1<br>XP_048590414.1 | *<znt2l*<br>XM_032369689.2<br>XP_032225580.2 | | | ***>aqp4L2***<br>**XM_001622599.3**<br>**XP_001622649.3** | *>ak2*<br>XM_048734283.1<br>XP_048590240.1 | *>edil3*<br>XM_048734282.1<br>XP_048590239.1 | *>edil3*<br>XM_048734556.1<br>XP_048590513.1 | *>edil3*<br>XM_048734555.1<br>XP_048590512.1 | >ncRNA<br>XR_007314102.1 | | | |
| 8 | *Exaiptasia diaphana* | NW_018387499.1<br>(Minus) | | *<dydc1*<br>XM_021053247.1<br>XP_020908906.1 | *>lrmda*<br>XM_021053244.2<br>XP_020908903.1 | *>fam241b*<br>XM_021053246.2<br>XP_020908905.1 | *<wapl*<br>XM_028661436.1<br>XP_028517237.1 | *<znt2l*<br>XM_021053242.2<br>XP_020908901.1 | | ***>aqp4L2.1***<br>**XM_021053240.2**<br>**XP_020908899.1** | ***>aqp4L2.2***<br>**XM_021053239.2**<br>**XP_020908898.1** | | *<naprtl*<br>XM_021053235.2<br>XP_020908894.1 | | *>hmgcl*<br>XM_028661430.1<br>XP_028517231.1 | *>jmjd6*<br>XM_028661435.1<br>XP_028517236.1 | *<dzip3*<br>XM_028661434.1<br>XP_028517235.1 | *<nlrc5l*<br>XM_028661433.1<br>XP_028517234.1 | |
| 9 | *Actinia tenebrosa* | NW_022258903.1<br>(Plus) | | *<dydc1*<br>XM_031697465.1<br>XP_031553325.1 | *>lrmda*<br>XM_031697443.1<br>XP_031553303.1 | *>fam241b*<br>XM_031697462.1<br>XP_031553322.1 | *<wapl*<br>XM_031697434.1<br>XP_031553294.1 | *<znt2l*<br>XM_031697428.1<br>XP_031553288.1 | | | ***>aqp4L2***<br>**XM_031697449.1**<br>**XM_031697449.1** | | *<naprtl*<br>XM_031697444.1<br>XP_031553304.1 | | *>hmgcl*<br>XM_031697427.1<br>XP_031553287.1 | *>slc12a8*<br>XM_031697441.1<br>XP_031553301.1 | <LOC116290432<br>XM_031697463.1<br>XP_031553323.1 | *<zdhhc6l*<br>XM_031697429.1<br>XP_031553289.1 | |

Direction: The arrows (>) indicate the transcriptional orientation of each gene on the genomic scaffold.

Gene Symbols: Official symbols are used where available; otherwise, NCBI LOC identifiers are provided.

Asterisks (*): Indicate full-length sequences determined in this study (refer to Supplementary Table S1 for sequence data).

#### Table S6. Summary of genes and accession numbers used for *aqp4L3* synteny analysis.

| # | Species | Chromosome (direction) | Upstream gene | Upstream gene | Upstream gene | Upstream gene | Upstream gene | Upstream gene | Upstream gene | Upstream gene | Upstream gene | Upstream gene | Upstream gene | Upstream gene | Target gene | Downstream gene | Downstream gene | Downstream gene | Downstream gene | Downstream gene |
|---|---|---|---|---|---|---|---|---|---|---|---|---|---|---|---|---|---|---|---|---|
| 1 | *Acropora millepora* | Chromosome 7<br>(Minus) | >*klhl*<br>XM_044323504.1<br>XP_044179439.1 | <*arf1*<br>XM_029351846.2<br>XP_029207679.1 | >ncRNA<br>XR_003827517.2 | >*phyh*<br>XM_044322962.1<br>XP_044178897.1 | <ncRNA<br>XR_006396203.1 | >*phyh*<br>XM_044322961.1<br>XP_044178896.1 | <*mmtag2*<br>XM_029351779.2<br>XP_029207612.2 | >*cdk5r1l*<br>XM_044323472.1<br>XP_044179407.1 | | | | <*myo1dl*<br>XM_029351749.2<br>XP_029207582.2 | ***>aqp4L3***<br>**XM_044322967.1**<br>**XP_044178902.1** | <*rsu-1l*<br>XM_044322966.1<br>XP_044178901.1 | >*stk11ipl*<br>XM_044322966.1<br>XP_044178901.1 | >ncRNA<br>XR_006396247.1 | <*tsl*<br>XM_029351429.2<br>XP_029207262.2 | >ncRNA<br>XR_003827332.2 |
| 2 | *Acropora digitifera* | NW_015441086.1<br>(Plus) | >*klhl*<br>XM_015923553.1<br>XP_015779039.1 | <*arf1*<br>XM_015923559.1<br>XP_015779045.1 | | | | >*phyh*<br>XM_015923557.1<br>XP_015779043.1 | | >*cdk5r1l*<br>XM_015923562.1<br>XP_015779048.1 | | | | <*myo1dl*<br>XM_015923561.1<br>XP_015779047.1 | ***>aqp4L3****<br>**XM_015923564.1**<br>**XP_015779050.1** | <*rsu-1l*<br>XM_015923563.1<br>XP_015779049.1 | >*stk11ipl*<br>XM_015923545.1<br>XP_015779031.1 | <*rnase inhibitor*<br>XM_015923546.1<br>XP_015779032.1 | < LOC107356917<br>XM_015923547.1<br>XP_015779033.1 | >*nod1l*<br>XM_015923566.1<br>XP_015779052.1 |
| 3 | *Porites lutea* | Chromosome 3<br>(Plus) | >*klhl*<br>XM_073380088.1<br>XP_073236189.1 | <*arf1*<br>XM_073380089.1<br>XP_073236190.1 | <*arf1*<br>XM_073380090.1<br>XP_073236191.1 | <*arf1*<br>XM_073380091.1<br>XP_073236192.1 | | >*phyh*<br>XM_073379153.1<br>XP_073235254.1 | <LOC140930406<br>pseudo | >*cdk5r1l*<br>XM_073380092.1<br>XP_073236193.1 | | | | <*myo1dl*<br>XM_073380093.1<br>XP_073236194.1 | ***>aqp4L3***<br>**XM_073381337.1**<br>**XP_073237438.1** | <*rsu-1l*<br>XM_073380094.1<br>XP_073236195.1 | >*stk11ipl*<br>XM_073380095.1<br>XP_073236196.1 | >ncRNA<br>XR_012164719.1 | >*slr1101l*<br>XM_073379155.1<br>XP_073235256.1 | >*slr1101l*<br>XM_073379156.1<br>XP_073235257.1 |
| 4 | *Stylophora pistillata* | NW_019218147.1<br>(Minus) | >*klhl*<br>XM_022944741.1<br>XP_022800476.1 | <*arf1*<br>XM_022944739.1<br>XP_022800474.1 | | | >*phyh*<br>XM_022944742.1<br>XP_022800477.1 | >*phyh*<br>XM_022944738.1<br>XP_022800473.1 | <*mmtag2*<br>XM_022944750.1<br>XP_022800485.1 | >*cdk5r1l*<br>XM_022944737.1<br>XP_022800472.1 | >*ox2r*<br>XM_022944735.1<br>XP_022800470.1 | < LOC111338309<br>XM_022944769.1<br>XP_022800504.1 | >LOC111338286<br>pseudo | <*myo1dl*<br>XM_022944753.1<br>XP_022800488.1 | ***>aqp4L3***<br>**XM_022944767.1**<br>**XP_022800502.1** | <*rsu-1l*<br>XM_022944768.1<br>XP_022800503.1 | >*stk11ipl*<br>XM_022944749.1<br>XP_022800484.1 | >*rya-r1l*<br>XM_022944766.1<br>XP_022800501.1 | <*sifar*<br>XM_022944765.1<br>XP_022800500.1 | <*npffr1*<br>XM_022944755.1<br>XP_022800490.1 |
| 5 | *Pocillopora damicornis* | NW_020844967.1<br>(Plus) | >*klhl*<br>XM_027186255.1<br>XP_027042056.1 | <*arf1*<br>XM_027186271.1<br>XP_027042072.1 | | | | >*phyh*<br>XM_027186244.1<br>XP_027042045.1 | <*mmtag2*<br>XM_027186242.1<br>XP_027042043.1 | >*cdk5r1l*<br>XM_027186281.1<br>XP_027042082.1 | | | | <*myo1dl*<br>XM_027186265.1<br>XP_027042066.1 | | <*rsu-1l*<br>XM_027186280.1<br>XP_027042081.1 | >*stk11ipl*<br>XM_027186245.1<br>XP_027042046.1 | >*rya-r1l*<br>XM_022944766.1<br>XP_022800501.1 | | |
| 6 | *Orbicella faveolata* | NW_018149991.1<br>(Minus) | >*klhl*<br>XM_020769433.1<br>XP_020625092.1 | <*arf1*<br>XM_020769386.1<br>XP_020625045.1 | <*arf1*<br>XM_020769431.1<br>XP_020625090.1 | | | >*phyh*<br>XM_020769469.1<br>XP_020625128.1 | <*mmtag2*<br>XM_020769468.1<br>XP_020625127.1 | >*cdk5r1l*<br>XM_020769402.1<br>XP_020625061.1 | | | | <*myo1dl*<br>XM_020769430.1<br>XP_020625089.1 | ***>aqp4L3***<br>**XM_020769429.1**<br>**XP_020625088.1** | <*rsu-1l*<br>XM_020769428.1<br>XP_020625087.1 | >*stk11ipl*<br>XM_020769406.1<br>XP_020625065.1 | >LOC110062481<br>XM_020769407.1<br>XP_020625066.1 | >*ankyrin-1-like*<br>XM_020769404.1<br>XP_020625063.1 | >LOC110062476<br>XM_020769403.1<br>XP_020625062.1 |
| 7 | *Nematostella vectensis* | Chromosome 2<br>(Plus) | | | | | <ncRNA<br>XR_004294073.2 | <*nkx-2.8*<br>XM_032373617.2<br>XP_032229508.1 | >*nkx-2.2*<br>XM_032373618.2<br>XP_032229509.1 | <ncRNA<br>XR_007306963.1 | <LOC116613034<br>XM_032373619.2<br>XP_032229510.1 | >*nkx-2.2*<br>XM_001625997.3<br>XP_001626047.2 | <ncRNA<br>XR_004294072.2 | > *nkx-2.2*<br>XM_001625998.3<br>XP_001626048.1 | ***>aqp4L3***<br>**XM_032373635.2**<br>**XP_032229526.1** | <*rsu-1l*<br>XM_001626014.3<br>XP_001626064.1 | <*tpr*<br>XM_032373632.2<br>XP_032229523.1 | >*kcnk3*<br>XM_048723782.1<br>XP_048579739.1 | >*kcnk3l*<br>XM_032373634.2<br>XP_032229525.1 | <*pla2g15*<br>XM_001626016.3<br>XP_001626066.1 |
| 8 | *Exaiptasia diaphana* | NW_018384631.1<br>(Plus) | | | | | <*aamp*<br>XM_021037541.2<br>XP_020893200.1 | >*tpgs1*<br>XM_021037542.2<br>XP_020893201.1 | >*copz1*<br>XM_021037420.2<br>XP_020893079.1 | <*usp37*<br>XM_028657079.1<br>XP_028512880.1 | <ncRNA<br>XR_002334293.2 | <*mgat1*<br>XM_021037480.2<br>XP_020893139.1 | >LOC110232347<br>XM_021037524.2<br>XP_020893183.1 | >*b4galnt3*<br>XM_028657106.1<br>XP_028512907.1 | | < *rsu-1l*<br>XM_021037406.2<br>XP_020893065.1 | >*stk11ipl*<br>XM_021037401.2<br>XP_020893060.1 | >*slr1101l*<br>XM_021037535.2<br>XP_020893194.1 | <LOC110232349<br>XM_021037525.2<br>XP_020893184.1 | <*catip*<br>XM_021037567.2<br>XP_020893226.1 |
| 9 | *Actinia tenebrosa* | NW_022257960.1<br>(Plus) | <*klhl*<br>XM_031694077.1<br>XP_031549937.1 | | | | | | | >*hlx*<br>XM_031694332.1<br>XP_031550192.1 | >*fp595*<br>XM_031694291.1<br>XP_031550151.1 | >*fp595*<br>XM_031694320.1<br>XP_031550180.1 | >*abracl*<br>XM_031694159.1<br>XP_031550019.1 | >*mettl24*<br>XM_031694239.1<br>XP_031550099.1 | ***>aqp4L3***<br>**XM_031694249.1**<br>**XP_031550109.1** | < *rsu-1l*<br>XM_031694109.1<br>XP_031549969.1 | <LOC116287416<br>XM_031694098.1<br>XP_031549958.1 | >*stk11ipl*<br>XM_031694087.1<br>XP_031549947.1 | >*slr1101l*<br>XM_031694148.1 | |

Direction: The arrows (>) indicate the transcriptional orientation of each gene on the genomic scaffold.

Gene Symbols: Official symbols are used where available; otherwise, NCBI LOC identifiers are provided.

Asterisks (*): Indicate full-length sequences determined in this study (refer to Supplementary Table S1 for sequence data).

### Table S7. Summary of genes and accession numbers used for *glps* synteny analysis.

| # | Species | Chromosome (direction) | Upstream gene | Upstream gene | Upstream gene | Upstream gene | Upstream gene | Upstream gene | Upstream gene | Upstream gene | Upstream gene | Upstream gene | Upstream gene | Upstream gene | Upstream gene | Upstream gene | Upstream gene | Target gene | Target gene | Intervening gene |
|---|---|---|---|---|---|---|---|---|---|---|---|---|---|---|---|---|---|---|---|---|
| 1 | *Acropora millepora* | Chromosome 4<br>(Minus) | | <*ethe1l*<br>XM_029343600.2<br>XP_029199433.2 | >*hnmtl*<br>XM_044318848.1<br>XP_044174783.1 | >*hnmtl*<br>XM_029343599.2<br>XP_029199432.1 | >*hnmtl*<br>XM_029343605.2<br>XP_029199438.2 | >*tctn3l*<br>XM_029343604.2<br>XP_029199437.2 | | | | <*myosin-15-like*<br>XM_029343643.2<br>XP_029199476.2 | <LOC114964268<br>XM_029343612.2<br>XP_029199445.2 | > LOC114964259<br>XM_029343601.2<br>XP_029199434.2 | >LOC114964259<br>XM_029343601.2<br>XP_029199434.2 | >LOC114964243<br>XM_029343584.2<br>XP_029199417.2 | >*taf8l*<br>XM_029343582.2<br>XP_029199415.1 | ***>glp1***<br>**XM_029343581.2**<br>**XP_029199414.2** | | |
| 2 | *Acropora digitifera* | NW_015441274.1<br>(Minus) | | | | | >*hnmtl*<br>XM_015909511.1<br>XP_015764997.1 | >*tctn3l*<br>XM_015909541.1<br>XP_015765027.1 | >*tctn3l*<br>XM_015909513.1<br>XP_015764999.1 | | <*ccdc18*<br>XM_015909508.1<br>XP_015764994.1 | <*spc110l*<br>XM_015909540.1<br>XP_015765026.1 | >ncRNA<br>XR_001564327.1 | >LOC107343917<br>XM_015909507.1<br>XP_015764993.1 | >ncRNA<br>XR_001564325.1 | >LOC107343912<br>XM_015909504.1<br>XP_015764990.1 | >*taf8l*<br>XM_015909505.1<br>XP_015764991.1 | ***>glp1****<br>**XM_015909506.1**<br>**XP_015764992.1** | | |
| 3 | *Porites lutea* | Chromosome 2<br>(Minus) | >*hsdl1*<br>XM_073378740.1<br>XP_073234841.1 | | | | <LOC140928934<br>XM_073378738.1<br>XP_073234839.1 | >*tctn3l*<br>XM_073378736.1<br>XP_073234837.1 | <LOC140927152<br>XM_073376803.1<br>XP_073232904.1 | >LOC140928932<br>XM_073378735.1<br>XP_073234836.1 | >LOC140928849<br>XM_073378650.1<br>XP_073234751.1 | >*gatml*<br>XM_073376109.1<br>XP_073232210.1 | >LOC140928930<br>XM_073378733.1<br>XP_073234834.1 | >LOC140926354<br>XM_073376107.1<br>XP_073232208.1 | <LOC140927151<br>XM_073376802.1<br>XP_073232903.1 | > LOC140927150<br>XM_073376801.1<br>XP_073232902.1 | >*taf8l*<br>XM_073376799.1<br>XP_073232900.1 | ***>glp1***<br>**XM_073378729.1**<br>**XP_073234830.1** | | |
| 4 | *Stylophora pistillata* | NW_019217958.1<br>(Minus) | >*hsdl1*<br>XM_022936773.1<br>XP_022792508.1 | | | <LOC111331619<br>XM_022936769.1<br>XP_022792504.1 | >*cenpe*<br>XM_022936764.1<br>XP_022792499.1 | < *tctn3l*<br>XM_022936768.1<br>XP_022792503.1 | >*ethe1l*<br>XM_022936757.1<br>XP_022792492.1 | >*lactadherin-like*<br>XM_022936756.1<br>XP_022792491.1 | | | | >ncRNA<br>XR_002694939.1 | <LOC111331596<br>XM_022936745.1<br>XP_022792480.1 | >LOC111331580<br>XM_022936723.1<br>XP_022792458.1 | >*taf8l*<br>XM_022936755.1<br>XP_022792490.1 | ***>glp1****<br>**XM_022936741.1**<br>**XP_022792476.1** | | |
| 5 | *Pocillopora damicornis* | NW_020844727.1<br>(Plus) | >*hsdl1*<br>XM_027184335.1<br>XP_027040136.1 | | | <LOC113668434<br>XM_027184334.1<br>XP_027040135.1 | >*cep290*<br>XM_027184326.1<br>XP_027040127.1 | < *tctn3l*<br>XM_027184331.1<br>XP_027040133.1 | >*ethe1l*<br>XM_027184338.1<br>XP_027040139.1 | >*lactadherin-like*<br>XM_027184366.1<br>XP_027040167.1 | >*nrp1*<br>XM_027184352.1<br>XP_027040153.1 | >*edil3*<br>XM_027184353.1<br>XP_027040154.1 | >*lactadherin-like*<br>XM_027184354.1<br>XP_027040155.1 | >*lactadherin-like*<br>XM_027184355.1<br>XP_027040156.1 | < LOC113668459<br>XM_027184359.1<br>XP_027040160.1 | >LOC113668464<br>XM_027184362.1<br>XP_027040163.1 | >*taf8l*<br>XM_027184361.1<br>XP_027040162.1 | ***>glp1***<br>**XM_027184360.1**<br>**XP_027040161.1** | | |
| 6 | *Pocillopora damicornis* | NW_020843631.<br>(Plus) | | | | | | | | | | | | | | | | | | |
| 7 | *Orbicella faveolata* | NW_018149354.1<br>(Plus) | | | | | | <*enpp5l*<br>XM_020758286.1<br>XP_020613945.1 | < *enpp5l*<br>XM_020758105.1<br>XP_020613764.1 | <*nfatc2ip*<br>XM_020757997.1<br>XP_020613656.1 | >*vkorc1l*<br>XM_020758003.1<br>XP_020613662.1 | >*mapk7*<br>XM_020757996.1<br>XP_020613655.1 | <*gtf2h3l*<br>XM_020758001.1<br>XP_020613660.1 | >LOC110051866<br>XM_020757947.1<br>XP_020613606.1 | >*hectd4*<br>XM_020757945.1<br>XP_020613604.1 | >ncRNA<br>XR_002297023.1 | <LOC110051988<br>XM_020758106.1<br>XP_020613765.1 | ***>glp1***<br>**XM_020758242.1**<br>**XP_020613901.1** | | < LOC110051857<br>XM_020757937.1<br>XP_020613596.1 |
| 8 | *Orbicella faveolata* | NW_018149303.1<br>(Plus) | | | | | | | | | | | | | | | | | | |
| 9 | *Nematostella vectensis* | Chromosome 3<br>(Plus) | | | | | | | | | <*dnajc25*<br>XM_048725024.1<br>XP_048580981.1 | >*hsd17b14*<br>XM_048725027.1<br>XP_048580984.1 | <*tspan33*<br>XM_032386956.2<br>XP_032242847.2 | > LOC5517057<br>XM_001637043.3<br>XP_001637093.3 | <*chymotrypsinogen*<br>XM_001636975.3<br>XP_001637025.3 | <*oct*<br>XM_048725049.1<br>XP_048581006.1 | >*taf8l*<br>XM_032387071.2<br>XP_032242962.1 | ***>glpα1.1***<br>**XM_001637041.3**<br>**XP_001637091.1** | ***>glp1.2***<br>**XM_048725185.1**<br>**XP_048581142.1** | |
| 10 | *Exaiptasia diaphana* | NW_018388314.1<br>(Plus) | | | | | | | | | | | | | | >LOC110253744<br>XM_021060697.2<br>XP_020916356.1 | >*taf8l*<br>XM_021060699.1<br>XP_020916358.1 | ***>glpα1.1***<br>**XM_021060680.2**<br>**XP_020916339.1** | | |
| 11 | *Exaiptasia diaphana* | NW_018387866.1<br>(Plus) | | | | | | | | | | | | | <*csmd1*<br>XM_021054317.2<br>XP_020909976.2 | <ncRNA<br>XR_003695810.1 | < *csmd3*<br>XM_028661781.1<br>XP_028517582.1 | ***>glpα1.2***<br>**XM_021054314.2**<br>**XP_020909973.1** | ***>glp1.3***<br>**XM_028661779.1**<br>**XP_028517580.1** | |
| 12 | *Actinia tenebrosa* | NW_022258437.1<br>(Plus) | | | | | | >LOC116287638<br>XM_031694323.1<br>XP_031550183.1 | >*grhpr*<br>XM_031694321.1<br>XP_031550181.1 | <LOC116287636<br>XM_031694319.1<br>XP_031550179.1 | >*vps18*<br>XM_031694329.1<br>XP_031550189.1 | <*motl*<br>XM_031694331.1<br>XP_031550191.1 | > LOC116287646<br>XM_031694330.1<br>XP_031550190.1 | >*psmd2l*<br>XM_031694335.1<br>XP_031550195.1 | >LOC116287630<br>XM_031694313.1<br>XP_031550173.1 | <*cela2a*<br>XM_031694342.1<br>XP_031550202.1 | >*taf8l*<br>XM_031694344.1<br>XP_031550204.1 | ***>glpα1.1***<br>**XM_031694314.1**<br>**XP_031550174.1** | ***>glpα1.2***<br>**XM_031694336.1**<br>**XP_031550196.1** | >ncRNA<br>XR_004196091.1 |

(Continued from the previous page)

| # | Species | Chromosome (direction) | Target gene | Intervening gene | Intervening gene | Intervening gene | Intervening gene | Intervening gene | Target gene | Target gene | Downstream gene | Downstream gene | Downstream gene | Downstream gene | Downstream gene | Downstream gene | Downstream gene | Downstream gene | Downstream gene |
|---|---|---|---|---|---|---|---|---|---|---|---|---|---|---|---|---|---|---|---|
| 1 | *Acropora millepora* | Chromosome 4<br>(Minus) | | | | | | | **>*glp2.1***<br>**XM_029343579.2**<br>**XP_029199412.2** | **< *glp2.2***<br>**XM_044318845.1**<br>**XP_044174780.1** | < LOC114964277<br>XM_044318843.1<br>XP_044174778.1 | < LOC114964278<br>XM_029343625.2<br>XP_029199458.2 | < LOC114964253<br>XM_029343596.2<br>XP_029199429.2 | <ncRNA<br>XR_003826048.2 | | >*sf3a1l*<br>XM_044318842.1<br>XP_044174777.1 | | | |
| 2 | *Acropora digitifera* | NW_015441274.1<br>(Minus) | | | | | | | **> *glp2.1****<br>**XM_015909539.1**<br>**XP_015765025.1** | **< *glp2.2****<br>**XM_015909538.1**<br>**XP_015765024.1** | < LOC107343911<br>XM_015909503.1<br>XP_015764989.1 | < LOC107343910<br>XM_015909502.1<br>XP_015764988.1 | <LOC107343909<br>XM_015909501.1<br>XP_015764987.1 | | | >*sf3a1l*<br>XM_015909537.1<br>XP_015765023.1 | | | |
| 3 | *Porites lutea* | Chromosome 2<br>(Minus) | | | | | | < *gpxl*<br>XM_073378731.1<br>XP_073234832.1 | **> *glp2.1a***<br>**XM_073378726.1**<br>**XP_073234827.1** | **> *glp2.1b***<br>**XM_073378730.1**<br>**XP_073234831.1** | >LOC140926353<br>XM_073376106.1<br>XP_073232207.1 | **< *glp2.2***<br>**XM_073376798.1**<br>**XP_073232899.1** | < LOC140927147<br>XM_073376797.1<br>XP_073232898.1 | < LOC140928924<br>XM_073378724.1<br>XP_073234825.1 | < LOC140926352<br>XM_073376105.1<br>XP_073232206.1 | > *sf3a1l*<br>XM_073378723.1<br>XP_073234824.1 | | | |
| 4 | *Stylophora pistillata* | NW_019217958.1<br>(Minus) | | | < LOC111331587<br>XM_022936730.1<br>XP_022792465.1 | < LOC111331586<br>XM_022936728.1<br>XP_022792463.1 | < LOC111331588<br>XM_022936731.1<br>XP_022792466.1 | < *gpxl*<br>XM_022936754.1<br>XP_022792489.1 | **> *glp2.1***<br>**XM_022936732.1**<br>**XP_022792467.1** | **< *glp2.2***<br>**XM_022936739.1**<br>**XP_022792474.1** | >ncRNA<br>XR_002694941.1 | >ncRNA<br>XR_002694940.1 | <*fx*<br>XM_022936727.1<br>XP_022792462.1 | <ncRNA<br>XR_002694944.1 | >LOC111331595<br>pseudo | > LOC111331614<br>XM_022936762.1<br>XP_022792497.1 | | | |
| 5 | *Pocillopora damicornis* | NW_020844727.1<br>(Plus) | | | >ncRNA<br>XR_003445998.1 | < *slc37a4l*<br>LOC113668465<br>pseudo | >trace amine-associated receptor 13c-like<br>XM_027184356.1 | | | | | | | | | | | | |
| 6 | *Pocillopora damicornis* | NW_020843631.<br>(Plus) | | | | | | < *gpxl*<br>XM_022936754.1<br>XP_022792489.1 | **> *glp2.1***<br>**XM_027191852.1**<br>**XP_027047653.1** | **< *glp2.2***<br>**XM_027191820.1**<br>**XP_027047621.1** | >LOC113675513<br>XM_027192109.1<br>XP_027047910.1 | <LOC113675524<br>XM_027192121.1<br>XP_027047922.1 | > LOC113675489<br>XM_027192091.1<br>XP_027047892.1 | > LOC113675133<br>XM_027191670.1<br>XP_027047471.1 | < LOC113675454<br>XM_027192055.1<br>XP_027047856.1 | >LOC113675368<br>pseudo | | | |
| 7 | *Orbicella faveolata* | NW_018149354.1<br>(Plus) | | <LOC110052161<br>XM_020758292.1<br>XP_020613951.1 | < LOC110051953<br>XM_020758065.1<br>XP_020613724.1 | < LOC110052015<br>XM_020758133.1<br>XP_020613792.1 | <*morc2*<br>XM_020758256.1<br>XP_020613915.1 | <ncRNA<br>XR_002297009.1 | | | | | | | | | | | |
| 8 | *Orbicella faveolata* | NW_018149303.1<br>(Plus) | | | | | | < *gpxl*<br>XM_020756059.1<br>XP_020611718.1 | **> *glp2.1***<br>**XM_020756053.1**<br>**XP_020611712.1** | **< *glp2.2***<br>**XM_020756051.1**<br>**XP_020611710.1** | <*hs3st3b1*<br>XM_020756047.1<br>XP_020611706.1 | <*hs3st3b1*<br>XM_020756048.1<br>XP_020611707.1 | < LOC110050165<br>XM_020756049.1<br>XP_020611708.1 | <*hs3st3b1*<br>XM_020756050.1<br>XP_020611709.1 | <*hs3st3b1*<br>XM_020756046.1<br>XP_020611705.1 | | | | |
| 9 | *Nematostella vectensis* | Chromosome 3<br>(Plus) | **< *glp1.3***<br>**XM_048725306.1**<br>**XP_048581263.1** | | | | | | | | >*pak*<br>XM_048724778.1<br>XP_048580735.1 | <*epha5*<br>XM_001633007.3<br>XP_001633057.3 | <*pex10*<br>XM_032382130.2<br>XP_032238021.2 | >*mct14*<br>XM_032382212.2<br>XP_032238103.1 | > LOC5512772<br>XM_048725711.1<br>XP_048581668.1 | >*pla2*<br>XM_032382138.2<br>XP_032238029.2 | | | |
| 10 | *Exaiptasia diaphana* | NW_018388314.1<br>(Plus) | | | | | | | | | < *pecr*<br>XM_021060688.1<br>XP_020916347.1 | > *fabl*<br>XM_021060698.1<br>XP_020916357.1 | <chymotrypsin-like elastase family member 2A<br>XM_021060687.2 | <XM_021060681.<br>2 | >XM_028663584.<br>1 | <XM_021060682.<br>2 | >ncRNA<br>XR_003696134.1 | >XM_028663583.<br>1 | <XM_028663593.<br>1 |
| 11 | *Exaiptasia diaphana* | NW_018387866.1<br>(Plus) | | | | | | | | | <*ethe1l*<br>XM_028661776.1<br>XP_028517577.1 | <*nk2r*<br>XM_028661778.1<br>XP_028517579.1 | >*tctn3l*<br>XM_028661780.1<br>XP_028517581.1 | >*tctn1l*<br>XM_028661777.1<br>XP_028517578.1 | >LOC110247841<br>XM_021054316.2<br>XP_020909975.1 | > LOC110247838<br>XM_021054313.1<br>XP_020909972.1 | | | |
| 12 | *Actinia tenebrosa* | NW_022258437.1<br>(Plus) | **> *glp1.3***<br>**XM_031694341.1**<br>**XP_031550201.1** | | | | | | | | <*ethe1l*<br>XM_031694339.1<br>XP_031550199.1 | >*tctn3l*<br>XM_031694338.1<br>XP_031550198.1 | >*kpr95*<br>XM_031694325.1<br>XP_031550184.1 | <*sglt1*<br>XM_031694326.1<br>XP_031550186.1 | <*sglt1*<br>XM_031694327.1<br>XP_031550187.1 | >*hsdl1*<br>XM_031694328.1<br>XP_031550188.1 | >*lrrc15*<br>XM_031694315.1<br>XP_031550175.1 | >LOC116287629<br>XM_031694312.1<br>XP_031550172.1 | |

Direction: The arrows (>) indicate the transcriptional orientation of each gene on the genomic scaffold.

Gene Symbols: Official symbols are used where available; otherwise, NCBI LOC identifiers are provided.

Asterisks (*): Indicate full-length sequences determined in this study (refer to Supplementary Table S1 for sequence data).

#### Table S8. Summary of genes and accession numbers used for *aqp12L1* synteny analysis.

| **#** | **Species** | **Chromosome (direction)** | **Upstream gene** | **Upstream gene** | **Upstream gene** | **Upstream gene** | **Upstream gene** | **Upstream gene** | **Upstream gene** | **Target gene** | **Downstream gene** | **Downstream gene** | **Downstream gene** | **Downstream gene** | **Downstream gene** | **Downstream gene** | **Downstream gene** | **Downstream gene** | **Downstream gene** |
|---|---|---|---|---|---|---|---|---|---|---|---|---|---|---|---|---|---|---|---|
| 1 | *Acropora millepora* | Chromosome 7 (Plus) | | | *>gabbr2*<br>XM_029332113.2<br>XP_029187946.2 | | *<mars*<br>XM_029332111.2<br>XP_029187944.2 | *>sgt*<br>XM_029332076.2<br>XP_029187909.1 | *>tnfsf10*<br>XM_029332078.2<br>XP_029187911.1 | ***>aqp12L1***<br>**XM_029332077.2**<br>**XP_029187910.2** | | *<aladin*<br>XM_029332075.2<br>XP_029187908.2 | | | *>mcm9*<br>XM_029332074.2<br>XP_029187907.2 | *<mchr2*<br>XM_029332166.2<br>XP_029187999.2 | *>ddx42l*<br>XM_029332123.2<br>XP_029187956.2 | | |
| 2 | *Acropora digitifera* | NW_015441081.1 (Minus) | | | *>gabbr2*<br>XM_015895542.1<br>XP_015751028.1 | | *<mars*<br>XM_015895466.1<br>XP_015750952.1 | *>sgt*<br>XM_015895464.1<br>XP_015750950.1 | *>tnfsf10*<br>XM_015895540.1<br>XP_015751026.1 | ***>aqp12L1***<br>**XM_015895537.1**<br>**XP_015751023.1** | | *<aladin*<br>XM_015895538.1<br>XP_015751024.1 | | | *>mcm9*<br>XM_015895541.1<br>XP_015751027.1 | *<mchr2*<br>XM_015895536.1<br>XP_015751022.1 | *>ddx42l*<br>XM_015895455.1<br>XP_015750941.1 | | |
| 3 | *Porites lutea* | Chromosome 3 (Plus) | | | *>gabbr2*<br>XM_073379848.1<br>XP_073235949.1 | | *<mars*<br>XM_073379847.1<br>XP_073235948.1 | *>sgt*<br>XM_073379067.1<br>XP_073235168.1 | | ***>aqp12L1***<br>**XM_073379068.1**<br>**XP_073235169.1** | | *<aladin*<br>XM_073379069.1<br>XP_073235170.1 | | | *>mcm9*<br>XM_073379850.1<br>XP_073235951.1 | <LOC140930187<br>XM_073379849.1<br>XP_073235950.1 | *>ddx42l*<br>XM_073379851.1<br>XP_073235952.1 | | |
| 4 | *Stylophora pistillata* | NW_019217885.1 (Minus) | | | *>gabbr2*<br>XM_022931711.1<br>XP_022787446.1 | | *<mars*<br>XM_022931709.1<br>XP_022787444.1 | *>sgt*<br>XM_022931721.1<br>XP_022787456.1 | | ***>aqp12L1***<br>**XM_022931722.1**<br>**XP_022787457.1** | *>ttc28*<br>XM_022931734.1<br>XP_022787469.1 | *<aladin*<br>XM_022931760.1<br>XP_022787495.1 | *>irc3*<br>XM_022931747.1<br>XP_022787482.1 | >irc3<br>XM_022931733.1<br>XP_022787468.1 | *>mcm9*<br>XM_022931753.1<br>XP_022787488.1 | <LOC111327544<br>XM_022931754.1<br>XP_022787489.1 | *>ddx42l*<br>XM_022931724.1<br>XP_022787459.1 | | |
| 5 | *Pocillopora damicornis* | NW_020847128.1 (Plus) | | | *>gabbr2*<br>XM_027199984.1<br>XP_027055785.1 | | *<mars*<br>XM_027199992.1<br>XP_027055793.1 | *>sgt*<br>XM_027199994.1<br>XP_027055795.1 | | ***>aqp12L1***<br>**XM_027199985.1**<br>**XP_027055786.1** | | *<aladin*<br>XM_027199997.1<br>XP_027055798.1 | | | *>mcm9*<br>XM_027199977.1<br>XP_027055778.1 | | *>ddx42l*<br>XM_027199990.1<br>XP_027055791.1 | | |
| 6 | *Orbicella faveolata* | NW_018149303.1 (Minus) | | | *>gabbr2*<br>XM_020765211.1<br>XP_020620870.1 | *<mars*<br>XM_020765237.1<br>XP_020620896.1 | *<mars*<br>XM_020765236.1<br>XP_020620895.1 | *>sgt*<br>XM_020765274.1<br>XP_020620933.1 | | ***>aqp12L1***<br>**XM_020765273.1**<br>**XP_020620932.1** | >ncRNA<br>XR_002297671.1 | *<aladin*<br>XM_020765272.1<br>XP_020620931.1 | | | *>mcm9*<br>XM_020765271.1<br>XP_020620930.1 | <LOC110058673<br>XM_020765327.1<br>XP_020620986.1 | *>ddx42l*<br>XM_020765270.1<br>XP_020620929.1 | | |
| 7 | *Nematostella vectensis* | Chromosome 2 (Minus) | | | *<phf10*<br>XM_048721517.1<br>XP_048577474.1 | *<fzd1*<br>XM_001647490.3<br>XP_001647540.1 | *<dstyk*<br>XM_048721511.1<br>XP_048577468.1 | <LOC116614550<br>XM_048721567.1<br>XP_048577524.1 | *<egln1*<br>XM_048721547.1<br>XP_048577504.1 | ***>aqp12L1***<br>**XM_032375666.2**<br>**XP_032231557.1** | *<inp*<br>XM_032375661.2<br>XP_032231552.1 | *>syt6*<br>XM_048724150.1<br>XP_048580107.1 | *>dnajb6*<br>XM_048724007.1<br>XP_048579964.1 | >LOC5499010<br>XM_001620607.3<br>XP_001620657.2 | *<hdac4*<br>XM_048723762.1<br>XP_048579719.1 | <LOC116614542<br>XM_032375640.2<br>XP_032231531.2 | <ncRNA<br>XR_007307033.1 | >adora2a<br>XM_032375696.2<br>XP_032231587.2 | |
| 8 | *Exaiptasia diaphana* | NW_018384872.1 (Plus) | | *>tra2b*<br>XM_021039476.2<br>XP_020895135.1 | >LOC110234112<br>XM_028657588.1<br>XP_028513389.1 | *>clasp1*<br>XM_021039473.2<br>XP_020895132.1 | | <XM_021039449.2 →<br>XP_020895108.1 | *<egln1*<br>XM_021039423.2<br>XP_020895082.1 | ***>aqp12L1***<br>**XM_021039447.2**<br>**XP_020895106.1** | | *<aladin*<br>XM_021039446.2<br>XP_020895105.1 | | | *>mcm9*<br>XM_021039413.2<br>XP_020895072.2 | <LOC110234059<br>XM_021039408.2<br>XP_020895067.1 | *>psmd11*<br>XM_021039466.2<br>XP_020895125.2 | *<mchr2*<br>XM_021039460.2<br>XP_020895119.1 | *>prrt1*<br>XM_021039463.2<br>XP_020895122.1 |
| 9 | *Actinia tenebrosa* | NW_022261460.1 (Plus) | | | <ncRNA<br>XR_004197778.1 | *>clasp1*<br>XM_031710928.1<br>XP_031566788.1 | <LOC116301816<br>XM_031710937.1<br>XP_031566797.1 | <LOC116301813<br>XM_031710932.1<br>XP_031566792.1 | >LOC116301815<br>XM_031710936.1<br>XP_031566796.1 | ***>aqp12L1***<br>**XM_031710936.1**<br>**XP_031566796.1** | | *<aladin*<br>XM_031710933.1<br>XP_031566793.1 | | | *>mcm9*<br>XM_031710931.1<br>XP_031566791.1 | <LOC116301806<br>XM_031710924.1<br>XP_031566784.1 | | | |

Direction: The arrows (>) indicate the transcriptional orientation of each gene on the genomic scaffold.

Gene Symbols: Official symbols are used where available; otherwise, NCBI LOC identifiers are provided.

Asterisks (*): Indicate full-length sequences determined in this study (refer to Supplementary Table S1 for sequence data).

Table S9. Water and solute permeabilities of *Acropora millepora* Aqp4Ls and Glps shown in Fig. 6.

| Protein | $P_{water}$ (×10$^{-6}$ cm/s, 100 mosM inside osmotic gradient) | $P_{glycerol}$ (×10$^{-6}$ cm/s, 180 mM outside solute gradient) | $P_{urea}$ (×10$^{-6}$ cm/s, 180 mM outside solute gradient) | $P_{boric\ acid}$ (×10$^{-6}$ cm/s, 180 mM outside solute gradient) |
|---|---|---|---|---|
| Control | 1.0 ± 0.36 (20) | 0.16 ± 0.16 (20) | 0.23 ± 0.26 (18) | 0.16 ± 0.14 (19) |
| Aqp4L1.1 | 8.3 ± 4.7 (9) | 0.82 ± 0.66 (11) | 0.64 ± 0.44 (9) | 0.67 ± 0.43 (9) |
| Aqp4L1.2 | 7.1 ± 2.3 (9) | 2.5 ± 0.94 (8) | 18 ± 11 (10) | 18 ± 8.4 (9) |
| Aqp4L1.3 | 3.3 ± 0.99 (12) | 0.26 ± 0.28 (13) | 0.10 ± 0.11 (12) | 0.40 ± 0.22 (12) |
| Aqp4L2 | 0.76 ± 0.42 (8) | 0.19 ± 0.23 (8) | 0.20 ± 0.19 (7) | 0.20 ± 0.058 (6) |
| Aqp4L3 | 0.91 ± 0.27 (12) | 0.29 ± 0.27 (11) | 0.35 ± 0.19 (11) | 0.19 ± 0.18 (11) |
| Glp1 | 1.6 ± 0.75 (11) | 4.7 ± 2.1 (10) | 0.21 ± 0.26 (10) | 0.22 ± 0.19 (10) |
| Glp2.1 | 6.4 ± 2.8 (12) | 54 ± 19 (15) | 1.9 ± 0.57 (14) | 2.5 ± 0.88 (13) |
| Glp2.2 | 7.4 ± 3.2 (12) | 43 ± 22 (11) | 3.2 ± 1.3 (11) | 3.3 ± 1.4 (12) |

Values are expressed as the mean ± standard deviation. Numbers in parentheses indicate the total number of oocytes tested.

Table S10. List of plots data of water and solute permeabilities shown in Fig. 6.

| | control | AmiAqp4L1.1 | AmiAqp4L1.2 | AmiAqp4L1.3 | AmiAqp4L2 | AmiAqp4L3 | AmiGlp1 | AmiGlp2.1 | AmiGlp2.2 |
|---|---|---|---|---|---|---|---|---|---|
| $P_{water}$ | 1.61 | 9.70 | 6.10 | 3.60 | 0.50 | 1.00 | 2.00 | 11.60 | 5.90 |
| (×$10^{-6}$ cm/s, 100 mosM inside osmotic gradient) | 0.63 | 12.90 | 9.90 | 3.90 | 0.60 | 1.20 | 1.80 | 5.40 | 14.50 |
| | 1.01 | 10.30 | 7.60 | 2.20 | 0.40 | 1.00 | 0.50 | 11.50 | 8.40 |
| | 1.26 | 8.50 | 3.60 | 2.30 | 0.10 | 1.20 | 0.60 | 6.90 | 6.20 |
| | 1.40 | 17.90 | 7.10 | 4.10 | 0.80 | 0.90 | 2.40 | 7.50 | 4.20 |
| | 1.32 | 3.90 | 10.60 | 4.00 | 1.10 | 0.90 | 2.70 | 6.38 | 9.50 |
| | 2.00 | 5.30 | 3.40 | 3.50 | 1.40 | 0.50 | 2.30 | 3.45 | 8.70 |
| | 0.85 | 3.60 | 7.20 | 4.90 | 1.20 | 1.00 | 2.30 | 3.86 | 8.50 |
| | 1.00 | 2.60 | 8.40 | 3.50 | | 1.30 | 0.90 | 2.72 | 7.90 |
| | 1.06 | | | 4.00 | | 0.90 | 1.10 | 3.84 | 1.10 |
| | 0.62 | | | 1.80 | | 0.50 | 1.20 | 6.39 | 8.90 |
| | 0.72 | | | 1.70 | | 0.50 | | 6.86 | 5.10 |
| | 1.07 | | | | | | | | |
| | 1.02 | | | | | | | | |
| | 0.45 | | | | | | | | |
| | 0.97 | | | | | | | | |
| | 0.62 | | | | | | | | |
| | 0.87 | | | | | | | | |
| | 1.06 | | | | | | | | |
| | 0.80 | | | | | | | | |
| $P_{glycerol}$ | 0.00 | 0.30 | 2.60 | 0.10 | 0.00 | 0.00 | 5.30 | 75.40 | 83.50 |
| (×$10^{-6}$ cm/s, 180 mM outside solute gradient) | 0.41 | 1.00 | 4.10 | 0.10 | 0.00 | 0.20 | 6.30 | 73.50 | 67.00 |
| | 0.42 | 0.00 | 2.70 | 0.00 | 0.50 | 0.10 | 3.90 | 55.90 | 41.40 |
| | 0.32 | 0.80 | 3.00 | 0.30 | 0.60 | 0.30 | 6.50 | 53.20 | 38.20 |
| | 0.20 | 1.10 | 3.00 | 1.10 | 0.30 | 0.20 | 6.60 | 72.50 | 34.20 |
| | 0.30 | 0.20 | 2.10 | 0.40 | 0.00 | 0.20 | 6.40 | 44.40 | 39.60 |
| | 0.10 | 2.30 | 0.60 | 0.00 | 0.00 | 0.00 | 2.90 | 48.70 | 56.20 |
| | 0.60 | 1.40 | 2.00 | 0.10 | 0.10 | 0.70 | 7.00 | 99.60 | 34.20 |
| | 0.00 | 0.10 | | 0.40 | | 0.20 | 1.50 | 46.43 | 9.50 |
| | 0.20 | 0.50 | | 0.40 | | 0.90 | 1.00 | 39.03 | 64.10 |
| | 0.00 | 1.30 | | 0.30 | | 0.40 | | 28.27 | 9.50 |
| | 0.20 | | | 0.10 | | | | 55.25 | |
| | 0.10 | | | 0.10 | | | | 39.86 | |
| | 0.10 | | | | | | | 29.37 | |
| | 0.00 | | | | | | | 41.11 | |
| | 0.10 | | | | | | | | |
| | 0.04 | | | | | | | | |
| | 0.04 | | | | | | | | |
| | 0.06 | | | | | | | | |
| | 0.07 | | | | | | | | |
| $P_{urea}$ | 0.17 | 0.70 | 14.30 | 0.20 | 0.10 | 0.10 | 0.10 | 2.10 | 2.70 |
| (×$10^{-6}$ cm/s, 180 mM outside solute gradient) | 0.54 | 0.50 | 8.80 | 0.10 | 0.60 | 0.10 | 0.00 | 1.50 | 0.90 |
| | 1.06 | 0.70 | 8.80 | 0.10 | 0.30 | 0.20 | 0.10 | 2.20 | 2.90 |
| | 0.00 | 0.70 | 27.90 | 0.00 | 0.10 | 0.40 | 0.10 | 1.60 | 2.20 |
| | 0.50 | 1.80 | 11.40 | 0.10 | 0.00 | 0.60 | 0.30 | 3.50 | 3.30 |
| | 0.10 | 0.20 | 14.50 | 0.00 | 0.20 | 0.10 | 0.00 | 1.10 | 4.40 |
| | 0.00 | 0.50 | 10.00 | 0.10 | 0.10 | 0.50 | 0.00 | 2.20 | 4.80 |
| | 0.20 | 0.40 | 10.70 | 0.00 | | 0.50 | 0.30 | 2.26 | 4.30 |
| | 0.30 | 0.30 | 29.10 | 0.10 | | 0.60 | 0.30 | 1.81 | 2.20 |
| | 0.00 | | 41.00 | 0.10 | | 0.30 | 0.90 | 1.89 | 2.20 |
| | 0.00 | | | 0.00 | | 0.40 | | 2.10 | 5.30 |
| | 0.40 | | | 0.40 | | | | 1.59 | |
| | 0.20 | | | | | | | 1.23 | |
| | 0.00 | | | | | | | 1.46 | |
| | 0.07 | | | | | | | | |
| | 0.24 | | | | | | | | |
| | 0.09 | | | | | | | | |
| | 0.24 | | | | | | | | |
| $P_{boric\ acid}$ | 0.05 | 0.40 | 29.80 | 0.50 | 0.30 | 0.10 | 0.00 | 2.00 | 5.60 |
| (×$10^{-6}$ cm/s, 180 mM outside solute gradient) | 0.07 | 0.40 | 6.70 | 0.90 | 0.20 | 0.60 | 0.40 | 3.00 | 5.20 |
| | 0.08 | 0.20 | 31.50 | 0.70 | 0.20 | 0.20 | 0.20 | 3.90 | 3.80 |
| | 0.37 | 1.30 | 25.10 | 0.20 | 0.20 | 0.20 | 0.70 | 2.80 | 4.60 |
| | 0.00 | 0.10 | 8.00 | 0.30 | 0.10 | 0.20 | 0.20 | 1.60 | 1.90 |
| | 0.30 | 1.40 | 14.10 | 0.50 | 0.20 | 0.40 | 0.20 | 2.40 | 4.00 |
| | 0.40 | 0.70 | 19.60 | 0.30 | | 0.00 | 0.10 | 2.10 | 3.00 |
| | 0.40 | 0.70 | 15.20 | 0.40 | | 0.00 | 0.10 | 0.72 | 4.20 |
| | 0.10 | 0.80 | 14.90 | 0.10 | | 0.10 | 0.10 | 1.77 | 0.50 |
| | 0.10 | | | 0.20 | | 0.30 | 0.20 | 3.91 | 2.00 |
| | 0.00 | | | 0.40 | | 0.00 | | 2.12 | 2.50 |
| | 0.00 | | | 0.30 | | | | 2.46 | 2.60 |
| | 0.10 | | | | | | | 3.32 | |
| | 0.10 | | | | | | | | |
| | 0.40 | | | | | | | | |
| | 0.22 | | | | | | | | |
| | 0.01 | | | | | | | | |
| | 0.10 | | | | | | | | |
| | 0.18 | | | | | | | | |
