## Supplementary material for "Solute selectivity and functional divergence of aquaporins in reef-building corals": FigS1-S6

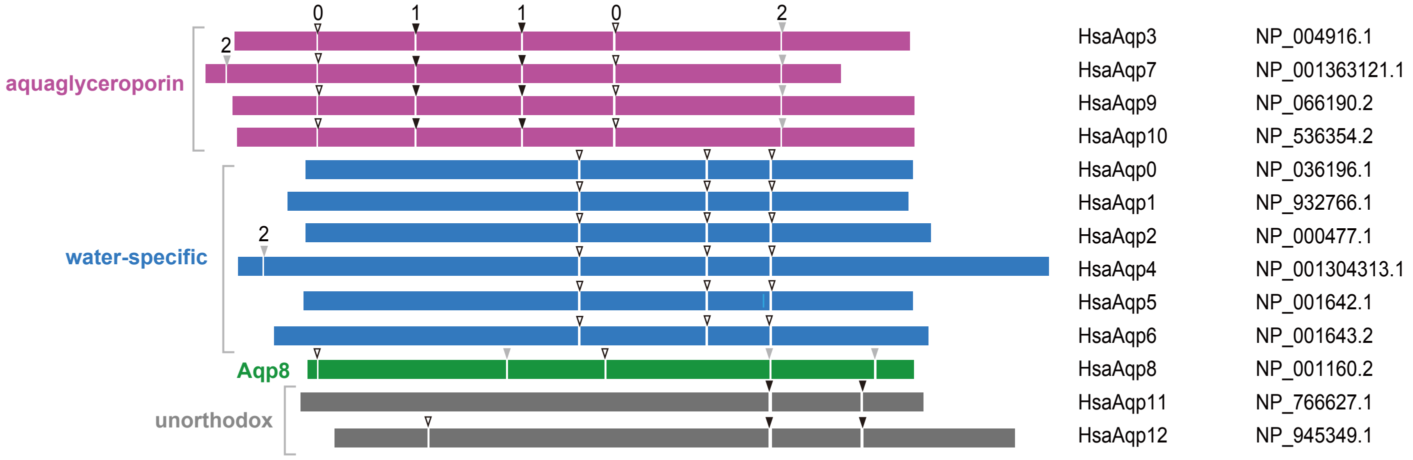


**Supplementary Fig. S1.** Exon–intron structures of human *aqp*s.
Exons are shown as colored boxes. Arrowheads indicate intron positions, and numbers denote intron phases (0, 1, or 2).

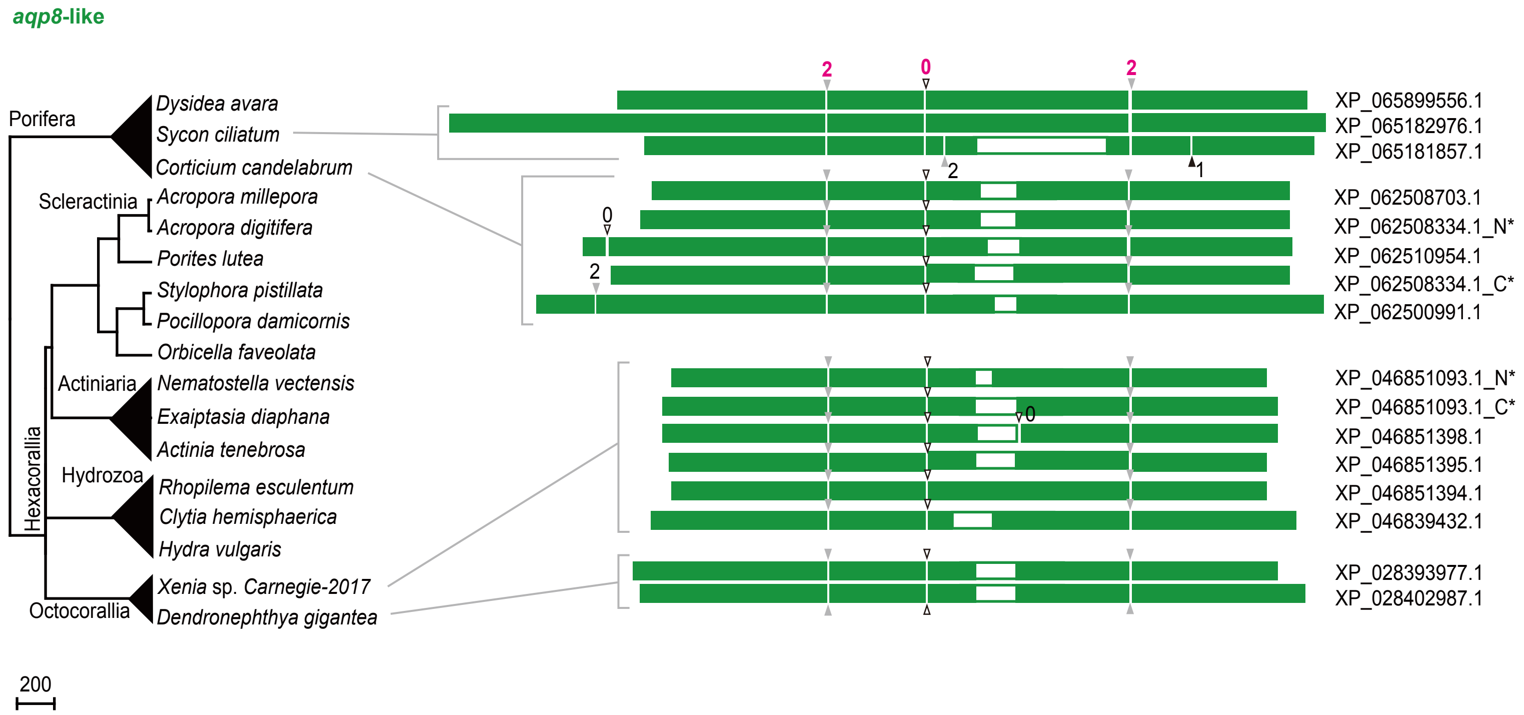


**Supplementary Fig. S2.** Exon–intron structures of *aqp8L* subfamilies.
Exons are shown as colored boxes. White regions within exons represent alignment gaps. Arrowheads indicate intron positions, and numbers denote intron phases (0, 1, or 2). Colored numbers indicate intron positions and phases conserved with the corresponding human AQP subfamilies (Fig. S1). Alignments are shown in Fig. S6. Species phylogenies are shown on the left. Asterisks (*) indicate full-length sequences determined in this study (Table S2).

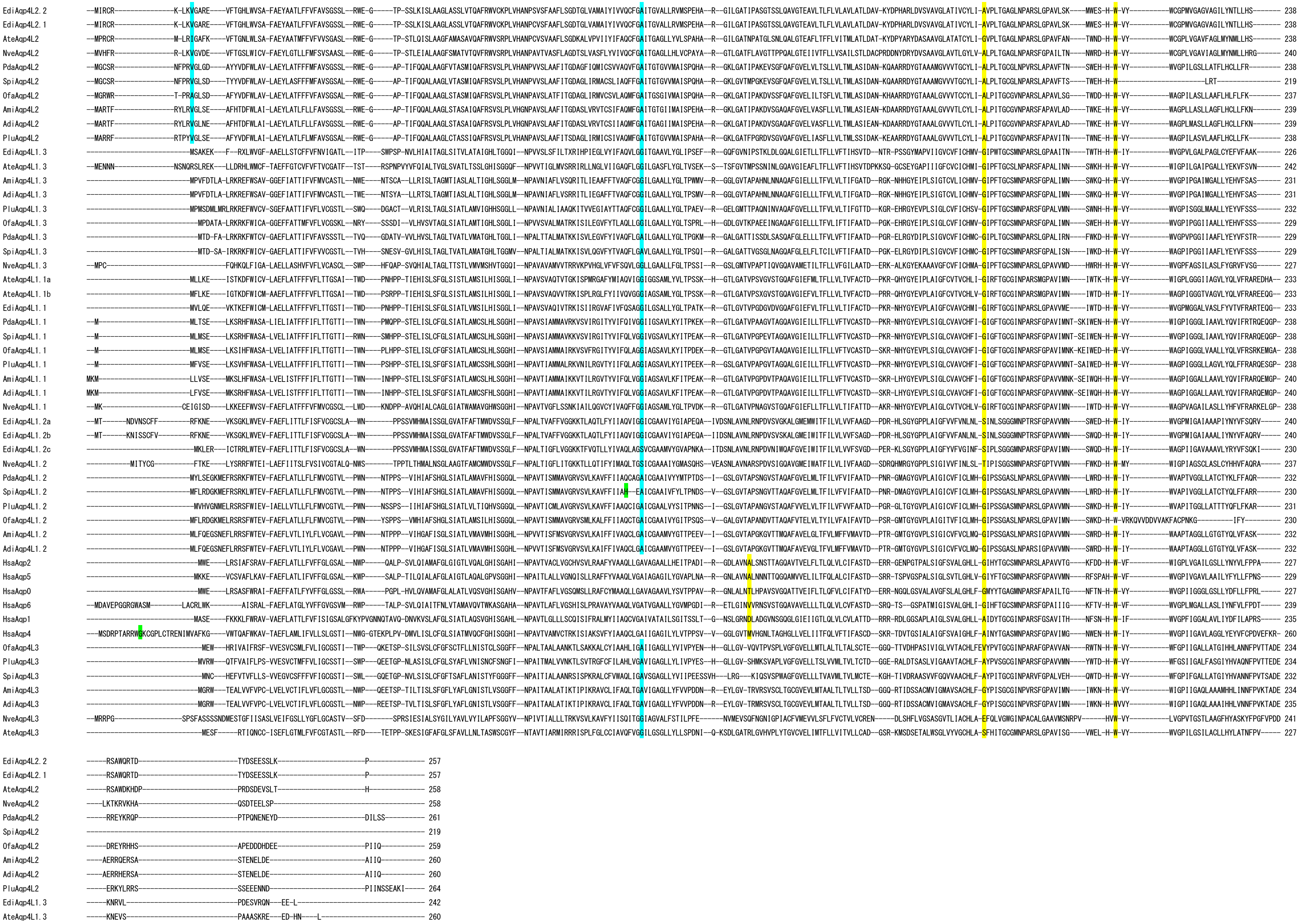
EdiAqp4L2.2
EdiAqp4L2.1
AteAqp4L2
NveAqp4L2
PdaAqp4L2
SpiAqp4L2
OfaAqp4L2
AmiAqp4L2
AdiAqp4L2
PluAqp4L2
EdiAqp4L1.3
AteAqp4L1.3
AmiAqp4L1.3
AdiAqp4L1.3
PluAqp4L1.3
OfaAqp4L1.3
PdaAqp4L1.3
SpiAqp4L1.3
NveAqp4L1.3
AteAqp4L1.1a
AteAqp4L1.1b
EdiAqp4L1.1
PdaAqp4L1.1
SpiAqp4L1.1
OfaAqp4L1.1
PluAqp4L1.1
AmiAqp4L1.1
AdiAqp4L1.1
NveAqp4L1.1
EdiAqp4L1.2a
EdiAqp4L1.2b
EdiAqp4L1.2c
NveAqp4L1.2
PdaAqp4L1.2
SpiAqp4L1.2
PluAqp4L1.2
OfaAqp4L1.2
AmiAqp4L1.2
AdiAqp4L1.2
HsaAqp2
HsaAqp5
HsaAqp0
HsaAqp6
HsaAqp1
HsaAqp4
OfaAqp4L3
PluAqp4L3
SpiAqp4L3
AmiAqp4L3
AdiAqp4L3
NveAqp4L3
AteAqp4L3

```
AmiAqp4L1.3    -----KMAVT----------------------------PA--------------------------------------------- 238
AdiAqp4L1.3    -----KMAVT----------------------------PA--------------------------------------------- 238
PluAqp4L1.3    -----KVTIS----------------------------PA--------------------------------------------- 239
OfaAqp4L1.3    -----KTEVV----------------------------PNVT------------------------------------------- 238
PdaAqp4L1.3    -----SAGVS----------------------------PA--------------------------------------------- 236
SpiAqp4L1.3    -----KTGVS----------------------------PS--------------------------------------------- 236
NveAqp4L1.3    -----SVAVE----------------------------QNDTNATELA------------------------------------- 242
AteAqp4L1.1a   ------SITE----------------------------PGGIVLNSL--MS-H-------------------------------- 249
AteAqp4L1.1b   ------SITT----------------------------PADIVLSNITPTN-HL----N-------------------------- 253
EdiAqp4L1.1    -----QSEGN----------------------------MDAVVLSNVTINN-NK------------------------------- 253
PdaAqp4L1.1    -----TA-GS----------------------------VNELDNY----QD-SK----I-------------------------- 254
SpiAqp4L1.1    -----TA-GS----------------------------VSELDNY----PD-SK----L-------------------------- 254
OfaAqp4L1.1    -----AA-ES----------------------------VNELDLY----AD-SK----L-------------------------- 254
PluAqp4L1.1    -----TA-GS----------------------------VNDLEMYH---PD-SK----M-------------------------- 255
AmiAqp4L1.1    -----TASAN----------------------------TNDLEMN----HD-AK----L-------------------------- 257
AdiAqp4L1.1    -----TASAT----------------------------TNDLEMN----HD-AK----L-------------------------- 257
NveAqp4L1.1    ---NADKAVG----------------------------PEDVELH---------------------------------------- 252
Aqp4L1.2a      ----YNSLLS----------------------------QPQ-------QSENST--------SNH-----DLDNENIQFQP---- 269
Aqp4L1.2b      ----YTKIAK----------------------------TSYDLNHDDVKAKKSG--------SVH-----SQEKINMDIQ----- 275c
Aqp4L1.2c      ----LQKYQA----------------------------P------DDRYDEGLP--------TVI-----QLNTYPI-------- 256
NveAqp4L1.2    ----INARIS----------------------------SAEEMPSKSHQNEGFNCLEVVCNKSQI-----SENGQETNVCRL--- 282
PdaAqp4L1.2    -----RNNRN----------------------------TLEDDNYNVELND-ND----TSK---------FMISPL--------- 261
SpiAqp4L1.2    -----RINQN----------------------------TLEDDNYNVELND-ND----TSK---------FMVSPL--------- 259
PluAqp4L1.2    -----KMTSNNN-------------------------INQENELYTVEVKD-NN----ESK---------SLVSSS--------- 263
OfaAqp4L1.2    --------------------------------------PSANVFYGILLSN-LS----FSECVFF----CYILGTCWPFERGFY- 267
AmiAqp4L1.2    -----RSNHD----------------------------SDNNMIHSMEVTEKDD----ATK---------FLDSTL--------- 262
AdiAqp4L1.2    -----RSNHD----------------------------SDNNMINSMEVMDKDD----ASK---------FLDSTL--------- 262
HsaAqp2        ------KSLSERLAVLKG------------------------LEPDTDWEEREV----RRRQSV------ELHSPQ-SLPRGTKA 271
HsaAqp5        ------LSLSERVAIIKGT-----------------------YEPDEDWEEQRE----ERKKTM------ELTTR---------- 265
HsaAqp0        ------KSISERLSVLKG------------------------AKPDVS-NGQPE----VTGEPV------ELNTQ--AL------ 263
HsaAqp6        ------KTLAQRLAILTGTV--------EVGTG-----AGAGAEPLKK-ESQP------GSGAV------EMESV---------- 282
HsaAqp1        ------SDLTDRVKVWTS---------------------GQVEEYDLDADD--------INSRV------EMKPK---------- 269
HsaAqp4        ---RFKEAFSKAAQQTKGSYMEVEDNRSQVETDDLILKPGVVHVIDVDRGEEKK----GKDQSG------EVLSSV--------- 323
OfaAqp4L3      -LVITEQELNDRAEPDKK-----EDTG-QT--------PRKYNIQTQT--------------------PYELKTTV--------- 275
PluAqp4L3      -LVITGKEIKNEAKSQGF-----ELEGPQH--------PSQADMQTQT--------------------PYELKTTV--------- 276
SpiAqp4L3      -LVIIGKELHGKAESAKR-----PDYQGEL--------PQSCDIQTQT--------------------PSELKTTV--------- 274
AmiAqp4L3      -LEVRGRQLKDKVEMPSFGL-NISEHHPPI--------PTQVTTQGDS--------------------PSELATFL--------- 280
AdiAqp4L3      -LEVRGRQLKDKVEMPSFGL-NISEHHPPI--------PTQVTTQGDS--------------------PSELATFL--------- 281
NveAqp4L3      TFLIKGEAIN----------------LPEVKRC-----ESGMGLESVIVEKPQP---VSPRFDRM-QIPDEMKTNL--------- 292
AteAqp4L3      ---ITDEDMA-----VKGEGIDL-DMDLEVVSDKIVIDNDDDHLSDKNMPS-----PILPDSPRLDQVPTEMKTYM--------- 289
```

**Supplementary Fig. S3.** Multiple sequence alignment of Aqp4Ls showing the conservation of intron positions. Intron insertion sites are indicated as follows: yellow, phase 0; blue, phase 1; and, green phase 2.

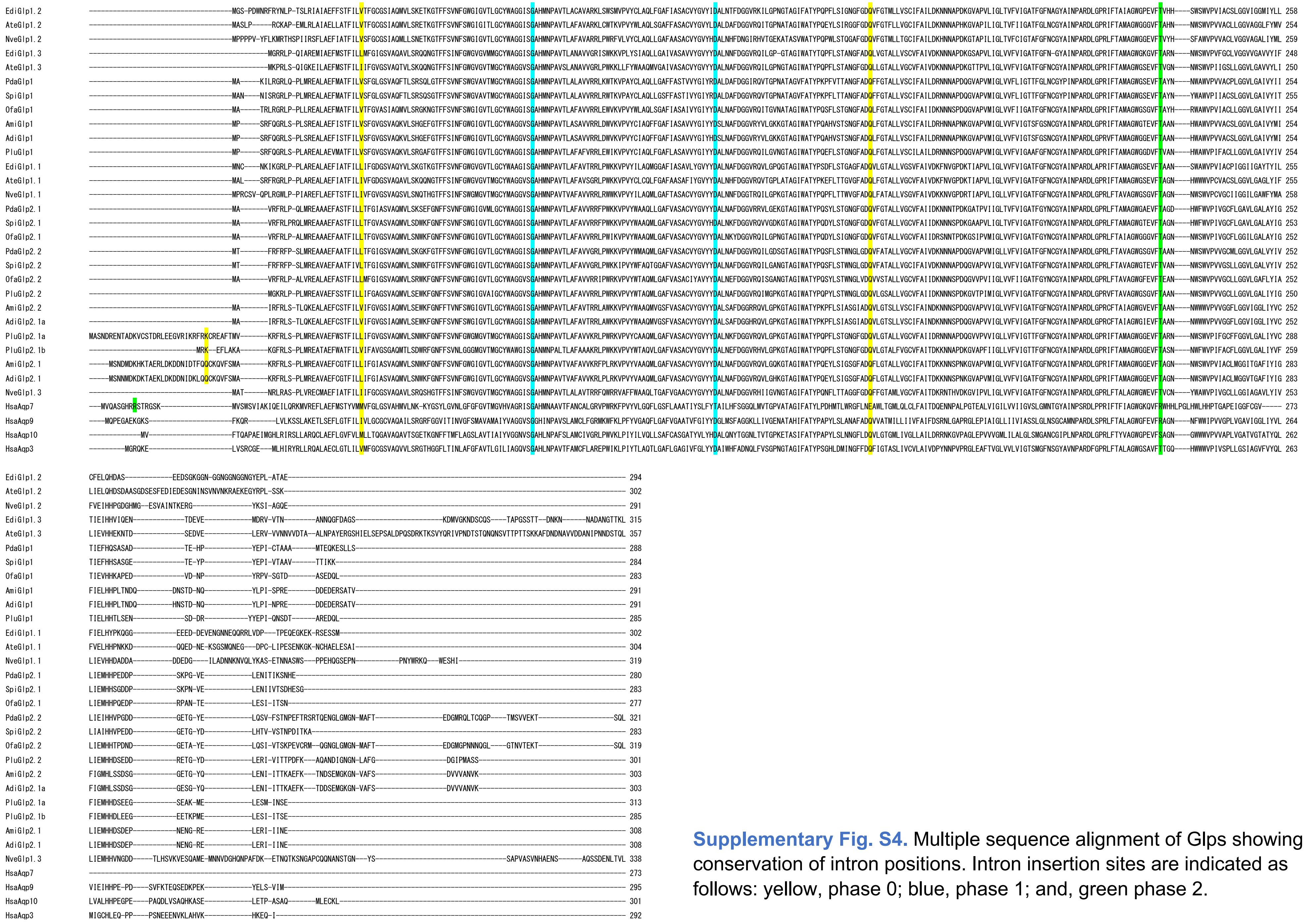

**Supplementary Fig. S4.** Multiple sequence alignment of Glps showing the conservation of intron positions. Intron insertion sites are indicated as follows: yellow, phase 0; blue, phase 1; and, green phase 2.

```
PdaAqp12L1     -----------MGFLWSLAAIFAGYFIGEIVRLALR-RVLPEV--YYPYANELISTFQLSVCVFEISVIGRFYNPW-IALWCSFVLLSVKNLGFVFEGALANPCG-----LLEDLVYD--QG-WLT-DNLAKIVFQVLGALLSFPFIQMLWLSTWSELHYQQ--VKKGLRSTLDVSLLYGFSIEILATFVSTTSDFLSRG-TLRRFNPV-------IRSVVCVLLSFFLSETTGTWMNPALATAQTFVYCSRKEVITEHLFVFWLGPIIGTLAAIEMNTLF--------LKPSVNTQRTK 258
SpiAqp12L1     -----------MGFLWSLVAISAGYFLGEIVRLALR-RILPRV--YYPYANELISTFQLSVCVFEISVIGRFYNSW-IALLCSFVLLTVKNVGFIFEGAFANPCG-----LLEDLVYE--KGYWLS-DNIAKIFFQVLGALLSFPFIQMLWLSTWSELHYQQ--IKRGLRSTLDVSLLYGFSIEILATFVSTASDFLSRG-TLRRFNPV-------IRSAICVLLSFFLSETTGTWMNPALATAQTFVYCSRNEVITEHLFVFWLGPIIGTLAAIEMNTLF--------LKPSVITQGRE 259
OfaAqp12L1     -----------MGFHWSLAAIAAGYLVGEFLREILR-RLLPKT--YYIYANELISTFQLSVCVFEISVIGRFYNVW-IALFCSFCLLTLKNAEYIFEGAFANPCG-----LMEDFVYK--KGYWIA-DNVTKIVFQLFGALLSFPFIQMLWKSTWSDLHHQQ--VRKGLRSTLEVSLLYGISIEILGTFVTTISDFLSRG-GLKKFNPV-------IRSALCVLVSYFLAETTGTWMNPALATAHSFVFCTKKEVIWEHLFVFWLGPIVGTLAAIEVNTRL--------LKPE--DKRKK 257
PluAqp12L1     -----------MVFHWSIGAISAGFLVGELFRIALK-RLLPSG--YYAYANELISTFQLSVCVFEISVIGKFYSSW-IALSCTFVLLALKHAEFIFEGAFANPCG-----MLEDIVYK--KAYWVK-ENIAKVIFQLIGGLLSFPFIQFLWKSTWSEFHFQQ--VKKGLRSTLEVSLLYGISIEIMATFVTVMSDFMSRGHALKKFNSV-------IRAASAVVVSYALAETTGTWMNPALATAQTFVYCTRKEVISEHLFVFWLGPVIGTMAAIVLNSLLTWNPKPVEMKPE------K 262
AmiAqp12L1     -----------MGFHWSLATVLAGFVLGELLRIVLK-RVLPKK--FHPYAKELISTFQLSVCAFEINVVGKFYSAW-VGLSCSFVLLTLKNTELILEGALANPCA-----IWEGVIDK--KGYFFTRENVAKITCQVIGGILSFPFTQVIWQAKWSDFHEKQ--VERGLQSTLEVSLMYGFFIETVGTFITTMCDFLSRGPPLKNFNAV-------IRAASVVVVCYFLAETTGVWMNPALATAQTFVFCTRQEAIHEHLFVFWLGPTIGTLIAIQLKSLLSM------SQPQ------- 256
AdiAqp12L1     -----------MGLHWSLATVLPGFFLGELLRFVLK-RVLPKK--FHPYAKELISTFQLSVYAFEINVVGKFYSAW-VGLSCSFVLLTLKNTELILEGALANPCA-----IWEGVIDK--KGYFFTRENVAKITCQVIGGILSFPFTQVIWQAKWSDFHEKQ--VERGLQSTLEVSLMYGFLIETLGTFITTMCDFLSRGPALKNFNAV-------IRAASIVVVCYFLAETTGVWMNPALATAQTFVFCTRQEAIHEHLFVFWLGPTIGTLIAIQVKSLLSM------SQPQ------- 256
EdiAqp12L1     -----------MAVIWSFSAIIGGFFIGELARLLLK-KLLTRKSRHYHYAGELISAFEFSAAVFEIAVIGKYYSTA-LGFVCSFFLLCLKNTEYILDGLSANPCG-----LLEGVVKRNARKSWMV-DFVTKVCSQLTGALLAFPFMQFVWKSTSSELHFKQ--LNQGLSSSLEVPLFVGFIVEITTTFLLTMTDFLTRG-GLRLFNPY-------IRAIACIVICYALTGTTGVWMNPVFATVHSFLFTKAXEDVIEHVFVFWAGPIVGTLIAISIDSKV--------RSPE----RKP 259
AteAqp121L     -----------MPLAWSFTAIIGAFVLGETVRFLLK-KIIPRRSPHHHYVGELISSFEFSAGVFEIAVIGKYYSNL-LGFIFSFTFLFLKNSEYILDRKAANPCG-----LLEGIVRRDVRRSWVV-DFVTKVSSQLTGALLAFPFMKLVWQSTASEIHFKQ--LSRGLSSSLEVSVLVGFSIEITATFVLTMIDFLTRG-GLRKFNPL-------VRAGACVLICYALTGTTGVWMNPVFATVHTFLFTDAKEDILEHFFVFWIAPIFGTLIAIAIDSKL--------HEPE----RKS 259
NveAqp12L1     -----------MDILWSVAAIIVVFLLGEIGRMIVKTNMLPR---HYKYASELISSFQFSACIFELVVIGRFYSPY-VGIACSFILLFLKNSEYIFDGSAANPCG-----LLEGVVCK-LKKPWFL-NFVLTVLFQFSGALLSFPVMRFMWQKTNSEMHFKQ--LNSELGTTLQVPIIIGFAIEVFTTFVVTMVDFVTRG-EMRRFNPI-------LRSTTCIAVCYALTGTTGVWMNPVFATVHTFLFTTGKELLLEHLFVFWVGPILGTLVAIGIDFKT--------HKHE---VQKP 257
HsaAqp11Homo   MSPLLGLRSELQDTCTSLGLMLSVVLLMGLARVVAR-QQLHRPV-AHAFVLEFLATFQLCCCTHELQLLSEQHPAH-PTWTLTLVYFFSLVHGLTLVGTSSNPCGVMMQMMLGGMSPE---------TGAVRLLAQLVSALCSRYCTSALWSLGLTQYHVSE--RSFACKNPIRVDLLKAVITEAVCSFL-----FHS---ALLHFQEVRTKLRIHLLAALITFLVYAGGSLTGAVFNPALAL--SLHFMCFDEAFPQFFIVYWLAPSLGILLMILMFSFF----LPW----------- 260
HsaAqp12Homo   ----------MAGLNVSLSFFFATFALCEAARRASK-ALLPVGA-YEVFAREAMRT---------LVELGPWAGDFGPDLLLTLLFLLFLAHGVTLDGASANPTVSLQEFLMAEQSLP---------GTLLKLAAQGLGMQAACTLMRLCWAWELSDLHLLQSLMAQSCSSALRTSVPHGALVEAACAFC-----FHLTLLHLRHSPPAYSGP---AVALLVTVTAYTAGPFTSAFFNPALAA--SVTFACSGHTLLEYVQVYWLGPLTGMVLAVLLHQGR----LPHLFQRNLFYGQK- 255
```

```
PdaAqp12L      NSLAKKIKQSSAK----NH-------------RNRAVNSNSEQKERLAKNA-------- 292
SpiAqp12L      KNLAKKIKQSSAK----NH-------------RDGAVNSNRNQKERFAKNA-------- 293
OfaAqp12L      KSLAKKIKKSSAK----NH-------------KNGTLKPSSEMKKRHAKNA-------- 291
PluAqp12L      KYWGKKDSRSSVE----NH-------------KNEELNPSELKKRRKGTKS-------- 296
AmiAqp12L      KAKDKKTNERWNG----NHLSGKT--------SKNEETNDGSEVRKRHTAKA-------- 296
AdiAqp12L      KAKDKQTNKRWNG----NHLSGKT--------SKNEETNDGSEVRKRHTAKA-------- 296
EdiAqp12L      TPISKTIRRKL------NKQLSKNG-IANGXHSMNG-TVQNCNKYEQKQTFSKKHK---- 307
AteAqp12L      PAIAKTILKKRKKHAVSNEAASQNGKVTNG--KANGVSVSNTKARQRKTNYGAAKK---- 313
NveAqp12L      KPIARTIKKHLQK----N-------------STNGRSLKQRNSRKLKQTSTGVR----- 294
HsaAqp11       --------------LHNNH------------TINKKE---------------------- 271
HsaAqp12       -----------------NKYRAPRGKPAP---ASGDTQTPAKGSSVREPGRSGVEGPHSS 295
```

**Supplementary Fig. S5.** Multiple sequence alignment of Aqp12Ls showing the conservation of intron positions. Intron insertion sites are indicated as follows: yellow, phase 0; blue, phase 1; and, green phase 2.

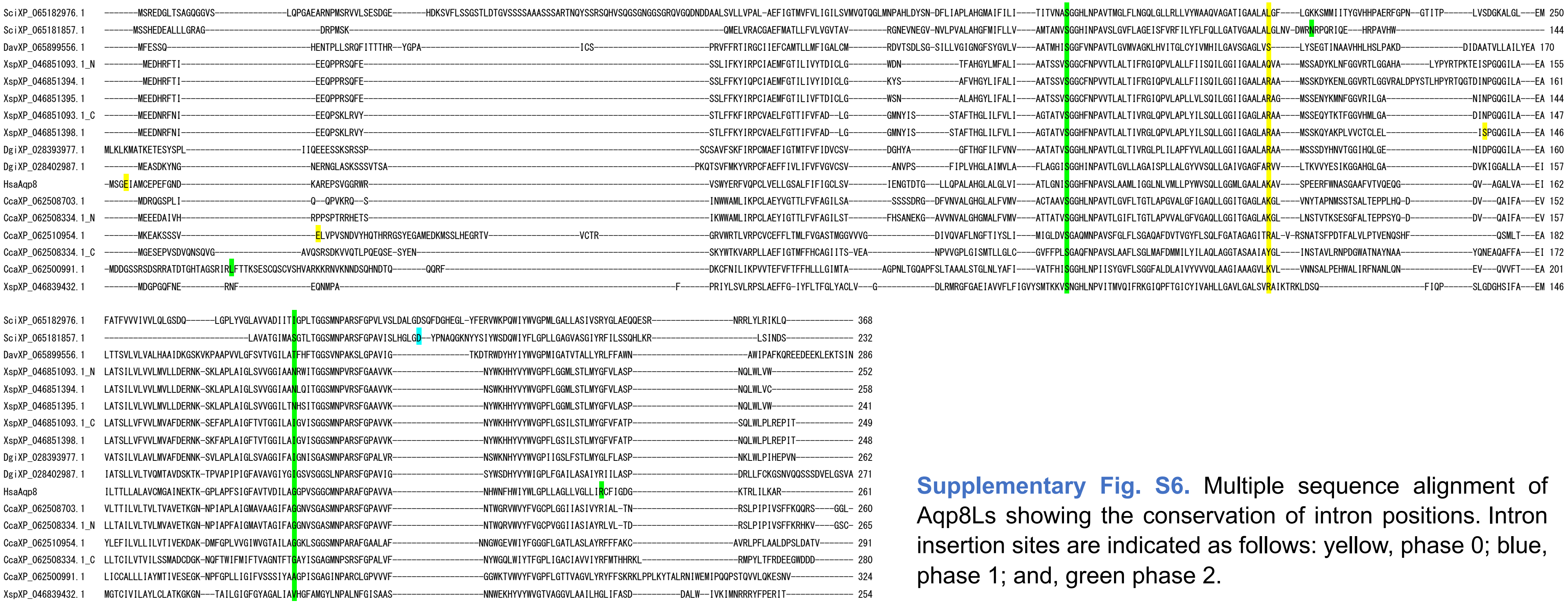

**Supplementary Fig. S6.** Multiple sequence alignment of Aqp8Ls showing the conservation of intron positions. Intron insertion sites are indicated as follows: yellow, phase 0; blue, phase 1; and, green phase 2.
